# The haplotype-resolved reference genome of lemon (*Citrus limon* L. Burm f.)

**DOI:** 10.1101/2021.07.14.452308

**Authors:** M. Di Guardo, M. Moretto, M. Moser, C. Catalano, M. Troggio, Z. Deng, A. Cestaro, M. Caruso, G. Distefano, S. La Malfa, L. Bianco, A. Gentile

## Abstract

Lemon (*Citrus limon* (L.) Burm. f.) is an evergreen tree belonging to the genus *Citrus*. The fruits are particularly prized for their organoleptic and nutraceutical properties of the juice.

Herein we report, for the first time, the release of a high-quality reference genome of the two haplotypes of lemon. The sequencing has been carried out coupling Illumina short reads and Oxford Nanopore data leading to the definition of a primary and an alternative assembly characterized by a genome size of 312.8 Mb and 324.74 Mb respectively. The analysis of the long terminal repeat (LTR) allowed the identification of 1921 regions on the primary and 1911 on the alternative assembly distributed across the nine chromosomes. Furthermore, an *in-silico* analysis of the microRNA genes was carried out using 246 mature miRNA and the respective pre-miRNA hairpin sequences of *C. sinensis*. Such analysis highlighted a high conservation between the two species with 233 mature miRNAs and 51 pre-miRNA stem-loops aligning with perfect match on the lemon genome.

In parallel, total RNA was extracted from fruit, flower, leaf and root enabling the detection of 38,205 and 37,753 predicted transcripts on primary and alternative assemblies respectively. Among those, the highest and lowest number of tissue-specific transcripts were detected in flower (2.73% and 2.71% in primary and alternative assemblies respectively) and leaf (0.7% and 0.68%) while gene ontology analysis enables a more precise characterization of the expressed genes based on their function.

The availability of a reference genome is an important prerequisite both for the set-up of high-throughput genotyping analysis and for functional genomic approaches toward the characterization of the genetic determinism of traits of agronomic interest.

## 1 Introduction

Lemon (*Citrus limon* (L.) Burm. f.) is ranked third for cultivated area in the world among the *Citrus* species and, together with lime, their global harvested area is more than one million hectares (FAOSTAT, 2019). Lemon cultivation has long been restricted to the coastal areas of Spain, Greece, and Italy while, nowadays, lemons are cultivated all over the world in areas characterized by Mediterranean-type climate (mild winters and cool summers) such as Argentina, Brazil, China, Mexico, Southern California, South Africa etc. Lemon is grown for fresh consumption and for processing. Fruits are widely priced for their quality in terms of flavor and nutraceutical value thanks to the high content in components such as: citric acid, vitamin C, flavonoids, and minerals (Sun et al., 2019) and for the essential oils in the peel (Mehl et al., 2014).

Even though the center of origin of lemon is still unknown, the species likely originated in an area comprising Eastern Himalaya, Middle East and India (Singh, 1981). This hypothesis is supported by the fact that such an area is characterized by the occurrence of lemons growing in a wild state (and bearing high quality fruit) and that most of the known *Citrus* species originated from the same area as well.

The use of molecular markers and whole genome sequencing (WGS) approaches provided fundamental insights to decipher the phylogeny of the genus *Citrus*. Cultivated citrus species are interspecific hybrid and/or admixture of three founder species (i.e., accessions without interspecific admixture): citron (*Citrus medica* L.), pummelo (*Citrus maxima* (Burm.) Merri) and mandarin (*Citrus reticulata* Blanco) (Wu et al., 2014). Lemon is a sour orange × citron hybrid, with the female parent being a F_1_ hybrid between pummelo and a pure mandarin. Analysis of cpDNA highlighted that lemon shares the chloroplast of pummelo through sour orange (Nicolosi et al., 2000; Gulsen and Roose, 2001; Carbonell-Caballero et al., 2015). WGS approaches confirmed the hybrid origin of lemon and sour orange, both characterized by high heterozygosity coupled with low interspecific diversity (Wu et al., 2018). The same study further elucidated the contribution of the three founders on the genetic makeup of lemon with citron, mandarin and pummelo characterizing the 50%, 19% and 31% of the lemon genome (Wu et al., 2018).

Even though several *Citrus* species were sequenced and assembled in the last years, like in the case of *Citrus* × clementina (Wu et al., 2014), pummelo, citron (Wang et al., 2017) and mandarin (Wang et al., 2018), the reference genome of lemon is not yet publicly available. In this paper we present the *de novo* sequencing of the cv ‘Femminello Siracusano’ lemon. The group of lemons named ‘Femminello’ represents the oldest, and most cultivated, Italian lemon variety (Barry et al., 2020). ‘Femminello’ cultivars usually show a marked ever-blooming and ever-bearing habit, which is particularly priced for the out-of-season production of *verdelli* lemons during the summer months (Barry et al., 2020). Among the ‘Femminello’ group, ‘Femminello Siracusano’ is one of the most widely cultivated thanks to the quality of the fruits coupled with the high yield.

The *de-novo* assembly of Illumina short reads and Oxford Nanopore data resulted in nine pseudomolecules that were built by synteny with the *Citrus maxima* genome (Wang et al., 2017), the only chromosome-scale reference genome among the relatives of lemon. The assembly was complemented by a *de-novo* annotation of the genes and of the non-coding regions (long terminal repeats and sRNA) of the genome.

## 2 Materials and Methods

### 2.1 DNA extraction and genome sequencing

Young leaves of the *Citrus limon* cultivar ‘Femminello Siracusano’ were sampled at the experimental farm of the University of Catania (Italy) in 2020. Total genomic DNA was isolated using the NucleoSpin Plant II Midi kit (Macherey Nagel, Germany). The same plant was also employed to collect leaf, flower, fruit and root tissue for RNA-seq analysis. Total RNA was extracted using the RNeasy Plant Power kit (Qiagen, Hilden, Germany) following the manufacturer protocol and tested for quality. As for the fruit, the total RNA was extracted separately from flavedo, albedo and pulp and then combined for mRNA sequencing. Library preparation for Illumina sequencing was performed using standard Illumina protocols and Illumina paired-ends adapters to generate reads of 2 × 150 nucleotides.

Long reads sequencing was performed using a modified version of the Oxford Nanopore (ONT) Ligation sequencing kit (SQK-LSK109). Briefly, 2 μg of extracted DNA were diluted to a final volume of 47 μl using nuclease free water. The DNA repair and end-prep step mix was then prepared as described in the 1D Lambda Control Experiment protocol (ONT) with an incubation time of 30’ at 20°C followed by 10’ at 65°C. Then 50 μl of AMPure XP beads were added to the 60 μl reaction and incubated 40’ at RT under gentle mixing. The washing steps were performed as described in the ONT protocol, but the elution step was extended to 20’. Similarly, the ligation was performed as described but the incubation extended to 20’. Subsequently, 40 μl of AMPure XP beads were added and, again, the mix incubated under gentle mixing for 20’. The remaining steps were performed as described in the ONT protocol with the final elution step extended to 20’. The preparation of the sample for the loading on the flow cell (R10.3 and R9.4.3) was performed as described by ONT.

### 2.2 Kmer analysis and genome size estimation

The illumina reads were analyzed to estimate the genome size and the level of heterozygosity. In particular, the k-mer spectrum with k-mer size of 21 was computed from the reads with Jellyfish v.2.3.0 (commands ‘count’ and ‘histo’) and then fed to the online tool genomescope (http://qb.cshl.edu/genomescope/; Vurture et al., 2017) with maximum kmer coverage set to 6000.

### 2.3 Genome assembly and curation

The genome sequence assembly was initially performed using MaSuRCA v.3.4.1 (Zimin et al., 2017) as it is an hybrid approach that directly combines the benefits of long reads with the accuracy of short reads. An alternative assembly was then performed by using falcon build 180808 (Chin et al., 2016) and afterwards polished with Polca (Zimin and Salzberg, 2020) bundled with MaSuRCA v.3.4.1, racon v1.4.17 (Vaser et al., 2017) and Pilon v.1.23 (Walker et al., 2014) to take full advantage of both types of reads also in this case.

To improve the definition of the two haplotypes, the information about the available parentals has been used. In particular, the assembled contigs have been aligned with BLAST v 2.11 (Altschul et al., 1990) against the sequences of pummelo and mandarin as representatives of the maternal haplotype (H1) and to citron as the representative of the paternal haplotype (H2). The alignments were filtered with a minimum similarity threshold of 95% and subsequently parsed to assign each contig to the most similar haplotype. The sequences of the two haplotypes were then de-duplicated to correct potential positioning errors by using Purge Haplotigs v.1.1.1 (Roach et al., 2018). The final set of contigs of the primary haplotype was built as the primary sequence of H1 plus the alternative sequence of H2. Similarly, the primary sequence of H2 and the alternative sequence of H1 was used for the alternative haplotype.

The pseudo-chromosomes were finally built by RaGOO v.1.11 (Alonge et al., 2019). The software was fed with pummelo as the guiding sequence because, among the currently available genomes, it is the only one that is organized into chromosomes. The completeness of the gene space was then assessed with BUSCO (Simão et al., 2015)

Finally, the synteny with other published citrus genomes (i.e. *Citrus maxima, Citrus medica* and *Citrus reticulata*) was visually inspected with the standalone version of D-Genies v.1.2.0 (Cabanettes and Klopp, 2018).

### 2.4 miRNA and LTR prediction and annotation

The sequences of known mature miRNA and the respective hairpin sequences of sweet orange (*Citrus sinensis* (L.) Osbeck) were retrieved from the miRBase repository (version 22.1). The mature sequences were aligned against the primary and alternative assemblies using bowtie (version 1.2.2; Langmead et al., 2009) with the option end-to-end (-v) with 0 mismatches. The miRNA hairpin sequences were blastn (Altschul et al., 1990) against the primary and alternative assemblies. Filters were adopted to assign miRNA loci with high confidence on the genome. Only miRNAs characterized by one locus on each genome assembly were considered for further filtering. The regions comprising the mature miRNA overlapping to the alignment of the associated hairpin, with a p-identity ≤ 95% and a ratio between the alignment length and the hairpin length ≥ 0.95, were retrieved and the coordinates reported for each assembly. In case that 5p and 3p mature miRNA were available then both had to be covered by the hairpin sequence alignment. miRNA that did not fulfill the above-mentioned conditions were classified as low confident loci. A manual inspection was performed on this set to select those miRNAs that could be clearly recognized but that did not pass the pident or the ratio cutoffs. All the others were reported in a separate list.

The presence of regions containing long terminal repeat (LTR) elements was assessed using LTR finder (Xu and Wang, 2007) with the parameters -L=5000 and -D=25000 (Du et al., 2018).

### 2.5 Gene prediction and annotation

Gene prediction was performed using AUGUSTUS (Stanke et al., 2008) on both primary and alternative assemblies trained with assembled transcripts from Bridger (version 2014-12-01; Chang et al., 2015). RNA-seq reads obtained from 4 different tissues: leaf, fruit, flower and root. Reads were filtered by quality, trimmed using Trimmomatic (version 0.39; Bolger et al., 2014) and independently assembled using Bridger. The four assembled datasets were then clustered together with CD-HIT (version 4.8.1; Li and Godzik, 2006) using a threshold of 99% identity. GeneMarkS-T (GeneMark version 3.20;GMST; Tang et al., 2015) was then employed to retain transcripts with high coding sequence potential. The resulting transcripts were translated into the corresponding amino acids sequences and aligned using BLAST+ (version 2.11; Camacho et al., 2009) to the Uniprot Uniref100 Viridiplantae dataset (The UniProt Consortiumx, 2019; Bateman et al., 2021). Only the transcripts with alignment similarity greater than 80% and alignment length greater than 90% (compared to the shortest sequence) were kept. Those high-quality sets of assembled transcripts were then aligned on the Primary assembly using GenomeThreader (GTH version 1.7.1; Gremme et al., 2005) and a GeneBank file comprising 1k upstream and downstream of genomic sequence was created for training AUGUSTUS.

RNA-seq alignments using GSNAP (version 2020-12-16; Wu et al., 2016) were used to create exon and intron hints files to be included in the final prediction of AUGUSTUS as specified in the AUGUSTUS online documentation. Only predictions with biological evidence from such alignments were considered and annotated using eggNOG (Huerta-Cepas et al., 2018), InterproScan (version 5.48.83; Jones et al., 2014) and BLAST+ alignment on the Uniprot Uniref100 Viridiplantae dataset.

The complete gene prediction pipeline has been created using Singularity (version 2.6) and Nextflow (version 20.10; Di Tommaso et al., 2017). All the scripts used to produce these results are available upon request.

## 3 Results and Discussion

### 3.1 Lemon DNA sequencing

The genomic DNA of the lemon cv. ‘Femminello Siracusano’ was extracted from young leaves and sequenced coupling Illumina paired end and ONT sequencing platforms. A total of 40.2 Gb Illumina short reads were generated (Table 1) and employed first to estimate the genome size and heterozygosity rate and then to perform hybrid assembly and polishing of the assembled sequence. As for the long reads sequencing, a total of 42.2 Gb reads were generated with the ONT platform with a N50 equal to 13.7, min and max read length equal to 0.1 and 139.9 Kbp and a median length of the reads of 4.4 Kbp (Table 1, Supplementary Figure 1).

**Table 1:** Illumina and Oxford Nanopore Technology (ONT) sequencing statistics.

|  | Number of reads (M) | Trimmed reads (M) | Number of Bases (Gbp) |
| --- | --- | --- | --- |
| DNA (Illumina) | 267 | - | 40.2 |
| DNA (ONT) | 5.04 | - | 42.2 |
| RNA (Flower) | 23.42 | 23.25 | 3.51 |
| RNA (Fruit) | 22.41 | 22.28 | 3.36 |
| RNA (Leaf) | 20.03 | 19.84 | 3.00 |
| RNA (Root) | 23.18 | 23.01 | 3.48 |

### 3.2 Kmer analysis and genome size estimation

The haploid genome size was estimated to be about 312 Mb with a heterozygosity level of 3.56% and about 182Mb of non-repetitive sequence (the 21-mer spectrum is reported in Supplementary Figure 2).

### 3.3 Genome assembly and curation

The genome assembly of Illumina and ONT reads has been carried out testing different assembly tools. A preliminary test was carried out employing the MaSuRCA genome assembler (Zimin et al., 2013). Sequences were assembled in 1,600 contigs with contig N50 and N80 equal to 2,548,919 (with 63 contigs) and 628,658 (with 232 contigs) respectively. The size of the assembled sequence was equal to 667,777 Mb, far larger than the genome size of the known ancestors of lemon ranging from 302 Mb (pummelo) to 405 Mb (citron) and to the estimated haploid genome size of ‘Femminello Siracusano’ of 312Mb. The genome assembly and annotation completeness were assessed using the BUSCO software (Simão et al., 2015). Results confirmed both the high quality of the sequenced data and the genome inflation since the complete sequence of the 99% of the genes tested was retrieved, but only a limited fraction, 15.4%, was detected in single copy. The relevant incidence of duplicated genes is largely a consequence of the high heterozygosity of the genome hampering an efficient detection of the homolog haplotypes. This issue, if not adequately tackled, often results in the duplication of the region.

In light of this, ONT and Illumina reads were assembled using FALCON while allelic contigs were reassigned using the BLAST (Altschul et al., 1990) alignment against the parental genomes in combination with the Purge Haplotigs pipeline (Roach et al., 2018). This approach led to the definition of a primary and an alternative haplotype: the first showed a genome size of 312.7986 Mb divided into 811 scaffolds with a N50 of 27.1 Mb. The alternative haplotype was instead characterized by a genome size of 324.74 Mb (divided into 799 scaffolds) with a N50 of 28.4 Mb. The genome assembly allowed the identification of nine pseudomolecules in each of the two assemblies (Figure 1A). The shortest and longest chromosomes were Chr. 8 (18.72 Mb and 20.09 Mb in the primary and the alternative assembly respectively) and Chr. 2 (51.41 Mb and 49.87 Mb) (Figure 1A, Table 2). The size of unanchored sequences was 39.31 Mb (divided into 802 contigs) and 38.81 Mb (in 790 contigs) in the two haplotypes (Supplementary Figure 3). The correlation between the chromosome length in lemon and pummelo was highly consistent (corr = 0.97 and 0.98 for the primary and alternative assemblies respectively), with chromosomes eight and two being the longest and shortest chromosomes in *Citrus Maxima* as well (Chr. 8 = 20.992 Mb; Chr. 2 = 52.984 Mb) (Supplementary Figure 3).

**Figure 1.**
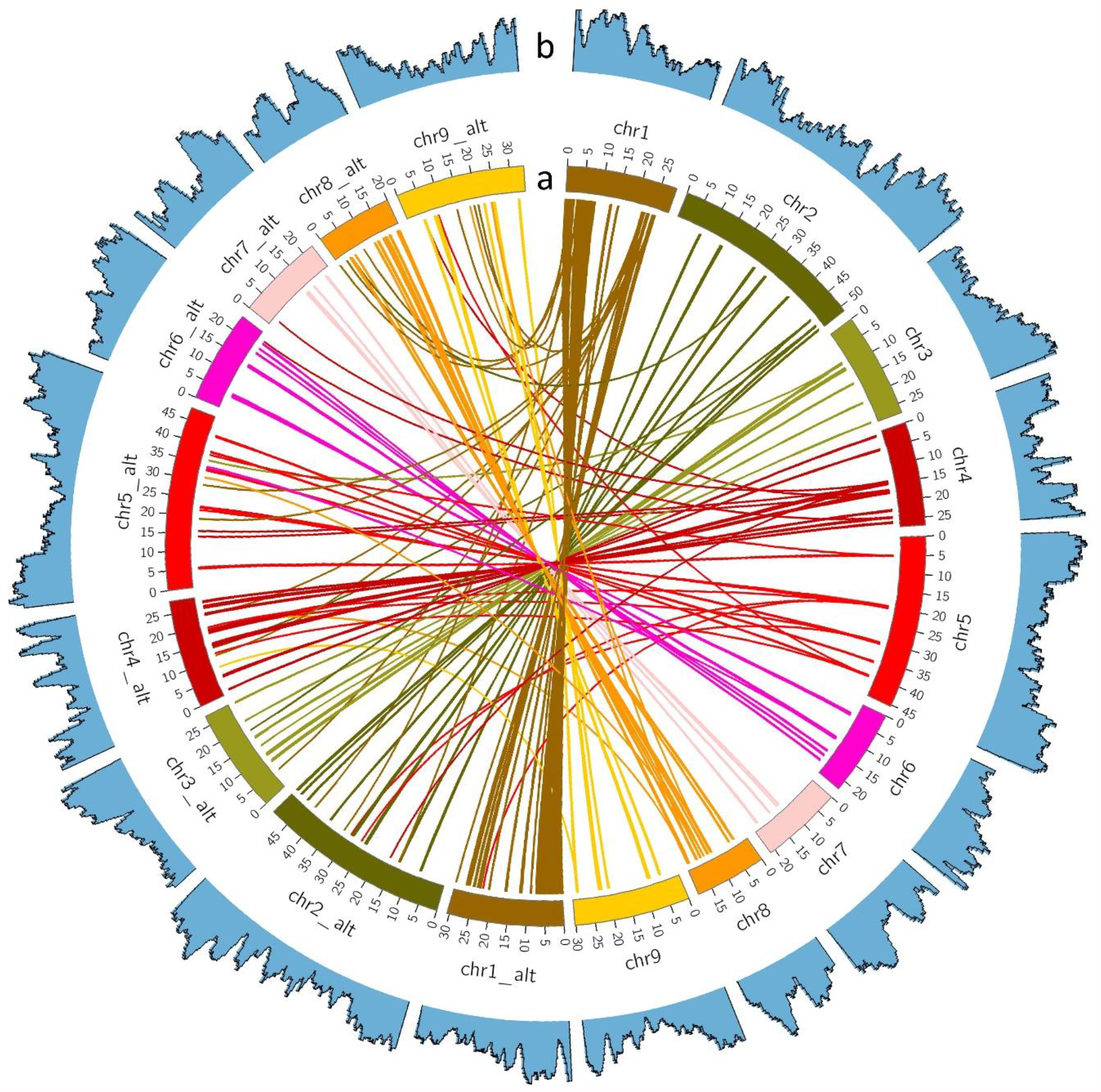
**(A)** Primary and alternative assemblies of the lemon genome. Chromosomes are represented as colored blocks, the two haplotypes of the same chromosome are represented by the same color. The position of genes showing a BLAST+ sequence identity of at least 96% and an alignment length of at least 60% compared to the smallest sequence are linked by straight lines. **(B)** Gene density distribution (blue histogram) across the primary and alternative assembly: the number of genes is calculated on a sliding window of 1 Mb with a 200 kb step.

**Table 2:** summary statistics of the primary and the alternative genome assembly. For each chromosome, the length in Mb is reported, as well as the number of genes, long terminal repeats (LTR) and the miRNA that were found with high confidence level detected.

| Chromosome | Length (Mb) |  | Genes |  | LTR |  | Mature MiRNA |  |
| --- | --- | --- | --- | --- | --- | --- | --- | --- |
|  | Primary | Alternative | Primary | Alternative | Primary | Alternative | Primary | Alternative |
| chr1 | 29.11929 | 30.652384 | 3428 | 3409 | 115 | 206 | 5 | 5 |
| chr2 | 51.414874 | 49.879738 | 5785 | 5648 | 305 | 295 | 51 | 29 |
| chr3 | 27.512417 | 27.616602 | 3374 | 3156 | 139 | 171 | 9 | 6 |
| chr4 | 27.165673 | 28.424913 | 3209 | 3146 | 141 | 175 | 11 | 10 |
| chr5 | 45.399278 | 48.255654 | 5816 | 5743 | 198 | 407 | 14 | 14 |
| chr6 | 22.460738 | 24.280274 | 2882 | 2957 | 107 | 155 | 10 | 10 |
| chr7 | 21.365374 | 22.905193 | 3032 | 3054 | 73 | 107 | 12 | 11 |
| chr8 | 18.720588 | 20.093211 | 2550 | 2510 | 642 | 173 | 7 | 6 |
| chr9 | 30.330071 | 33.825048 | 3186 | 3193 | 201 | 222 | 8 | 7 |

Based on the estimated genome size (312.073 Mb) it is likely that the alternative haplotypes might contain a small portion of contigs (i.e., around 5-6 Mb) that should be present in the primary sequence but that with the available information it was not possible to place correctly. The BUSCO analysis showed a significant number of complete sequences (95.9% and 94.8% in primary and alternative haplotypes respectively) with a slightly increased number of duplicated sequences in the alternative haplotype (8.8% and 10.0% for primary and alternative haplotypes respectively).

### 3.4 miRNA and LTR prediction and annotation

The annotation of microRNA genes on the genome was carried out performing an *in-silico* analysis using the mature miRNA and the respective pre-miRNA hairpin sequences of *Citrus sinensis* present in the miRBase repository. The dataset was selected being the most complete in terms of the number of miRNA annotated for a citrus species and considering the relatively low genetic distance between our genotype and sweet orange. The alignment of the 246 mature miRNA sequences showed that the miRNAs are well conserved between *Citrus sinensis* and *Citrus limon* with 233 aligning with perfect match on the genome, whereas 13 showed differences at nucleotide level. The alignments of the hairpin sequences produced hits with perfect identity for 54 out of 151 miRNA hairpins, 28 with identity between 99% and <100%, 27 with identity between 98% and <99%, 11 with identity between 97% and <98%, 8 with identity between 96% and <97% and 4 with identity between 95% and <96%. Among the remaining hairpins, 10 were characterized by an identity between 90% and <95% and the remaining by an identity <90% with several mismatches and gaps or only with partial alignments. However, most of the hairpins (115 out of 151 unique hairpins in the miRBase dataset corresponding to 76%) were positioned univocally (Table 2, Supplementary Table 1) on the chromosomes intersecting the alignment information associated with the mature miRNA alignments. In addition, 22 loci characterized by the miRNA hairpin/mature association were assigned with lower confidence (Supplementary Table 2). A minor number of miRNA hairpin/mature pairs did not pass the filters applied in the analysis and for these either no information could be retrieved due to the poor alignment quality (csi-miR3950, csi-miR3952, csi-miR3954) or the annotation was not confident enough to univocally assign the position on the genome (csi-miR169a, csi-miR395a, miR395c, csi-miR396b, csi-miR399c, csi-miR399a, csi-miR3948). The results indicate that the miRNAs are well conserved in *C. limon* compared to other members of the *Citrus* genus. Although this was expected for the mature miRNAs it is noteworthy that almost one third of the presented 100% identity with those of *Citrus sinensis*, while 131 miRNA hairpins showed a range of identity higher or equal to 95%.

The analysis on the LTR elements identified 1,921 regions on the primary assembly and 1,911 on the alternative assembly (Supplementary Table 3, Supplementary Table 4). The distributions on the chromosomes varied from 73 on Chr. 7 to 642 on Chr. 8 for the primary assembly and from 107 on Chr. 7 to 407 on Chr. 5 for the alternative assembly (Table 2). The scores associated with the LTR regions ranged from 6 (default minimum threshold) to 8 indicating that not all the known elements forming a complete LTR retrotransposon were found in any of the identified LTR regions on our genome.

### 3.5 Gene prediction and annotation

Raw paired-ends RNA-seq reads coming from four tissues: 20,026,773 from leaf, 23,182,013 from root, 22,414,235 from flower and 23,416,205 from fruit, were filtered and trimmed using Trimmomatic (Bolger et al., 2014) resulting in 19,843,227, 23,007,618, 22,277,820 and 23,250,076 sequences respectively. The four datasets were independently assembled using Bridger (Chang et al., 2015) resulting in 66,802 leaf, 66,937 root, 88,196 flower and 81,260 fruit transcripts. Those datasets were merged using CD-HIT (Li and Godzik, 2006) with a similarity threshold of 99% sequence identity and lead to a unified dataset of 205,757 transcripts. GMST (Tang et al., 2015) identified 96,700 of such transcripts to be more likely to code for proteins. The alignment of those transcripts against the UniProt Uniref100 Viridiplantae dataset allowed to further filter 24,880 putative complete transcripts. GenomeThreader (Gremme et al., 2005) was able to completely align 11,964 transcripts on the primary assembly and a genomic region comprising 1kbps upstream and downstream the alignment were retained for each transcript in order to create a GeneBank file to train AUGUSTUS (Stanke et al., 2008). After the training and optimization process, AUGUSTUS was used to predict genes on both primary and alternative assemblies using RNA-seq alignment results as biological hints to further enhance the prediction results. The final prediction consists of 38,205 and 37,753 predicted transcripts on primary and alternative assembly respectively (Table 2, Figure 1B).

Among those, 24,603 and 24,385 predicted transcripts were detected in all four tissues in the primary and alternative assembly respectively (Figure 2A). Overall, the number of predicted transcripts detected in the two assemblies was highly consistent in all the sets depicted in the Venn diagrams (average difference equal to 4.8% with standard deviation = 0.05). The set composed by the transcripts in common between leaf and fruit showed the highest relative difference (19.2%) between primary (130 transcripts) and the alternative (109 transcripts). The highest number of tissue-specific transcripts were detected in Flower (963 and 938 in the primary and alternative assembly) while the lowest value was detected in Leaf (116 and 114, Figure 2) with a relative frequency compared to all the other predicted transcripts equal to 2.73% and 2.71% for Flower and 0.7% and 0.68% for Leaf in the primary and alternative assembly respectively (Fig. 2B).

**Figure 2.**
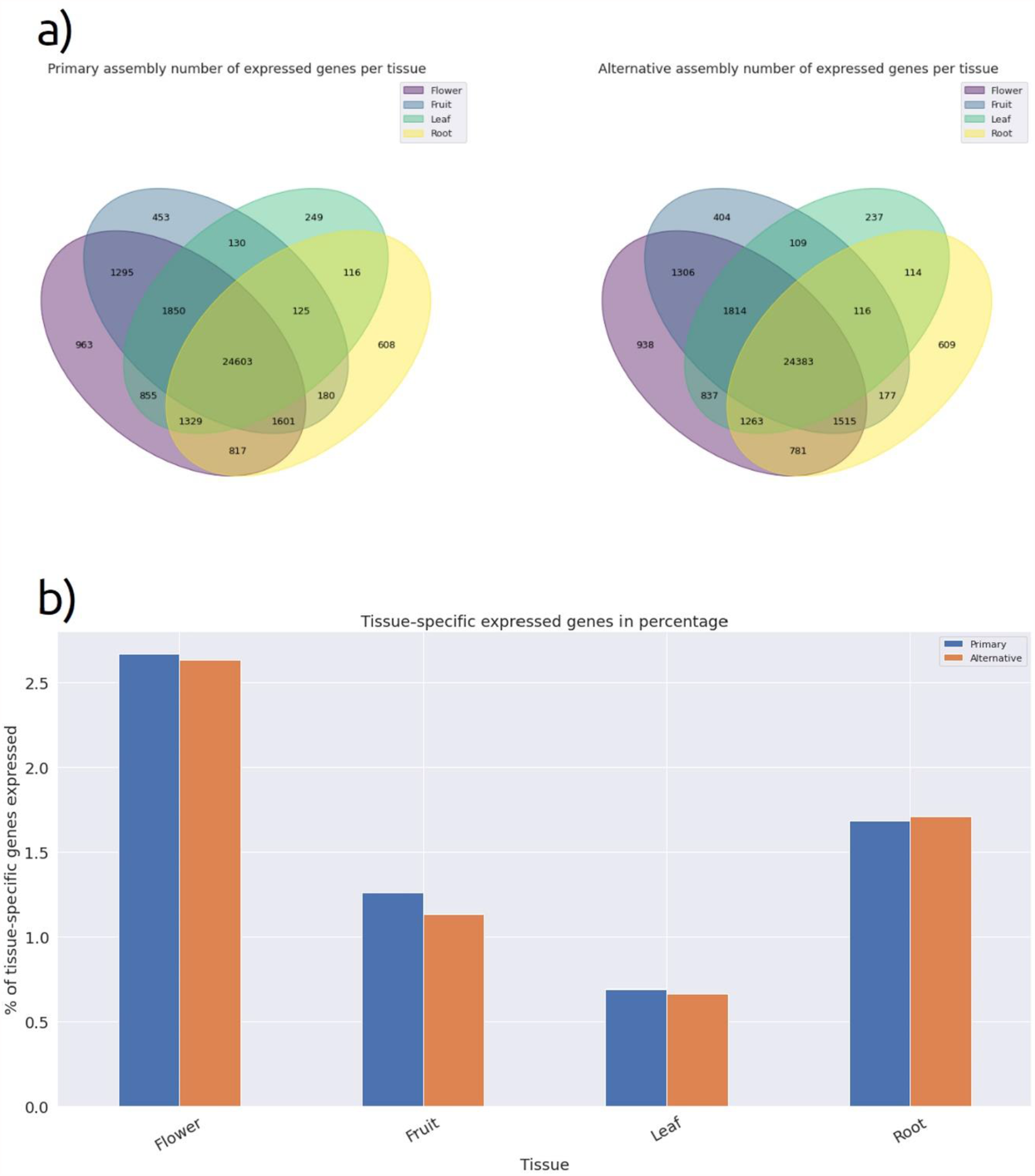
**(A)** Venn diagram with number of expressed genes in flower, fruit, leaf and root for the two haplotypes. **(B)** Bar-plot representing the percentages of genes uniquely expressed in flower, fruit, leaf and root in for the two haplotypes.

Gene Ontology functional annotation was performed using eggNOG and resulted in 28,338 and 27,895 annotated transcripts respectively for primary and alternative assembly. InterProScan assigned 29,117 and 28,645 transcripts at least one annotation term while the BLAST+ alignment on the Viridiplantae Uniprot Uniref100 dataset resulted in 30,213 and 29,737 hits with a putative known protein. The relative frequency of the genes according to the GO categories is depicted in Supplementary Figure 4

In addition, an enrichment analysis was performed using GOATOOLS (Klopfenstein et al., 2018) on the four tissues and for each of the three GO aspects (Supplementary figure 5). To provide a smaller and more readable GO graph, figure 3 shows a smaller version of the GO enrichment for leaf and root, containing only the most significant GO categories. As shown in Fig. 3A, several GO terms related to ‘biological process’ were significantly more represented in flower compared to the other tissues, notably significant differences were detected for GO terms related to reproductive system development, reproductive structure development, plant organ development, developmental process involved in reproduction, anatomical structure morphogenesis for a total of 185 genes. As for the root tissue, several GO terms were significantly more represented compared to all the annotated genes. Some of these GO terms were related to response to exogenous or endogenous stimuli (response to chemical, 60 genes; response to hormone, 39 genes; response to endogenous stimulus, 60 genes) while other were related to metabolic process (organic substance metabolic process, 60 genes; primary metabolic process, 43 genes; cellular metabolic process, 55 genes), figure 3B.

**Figure 3.**
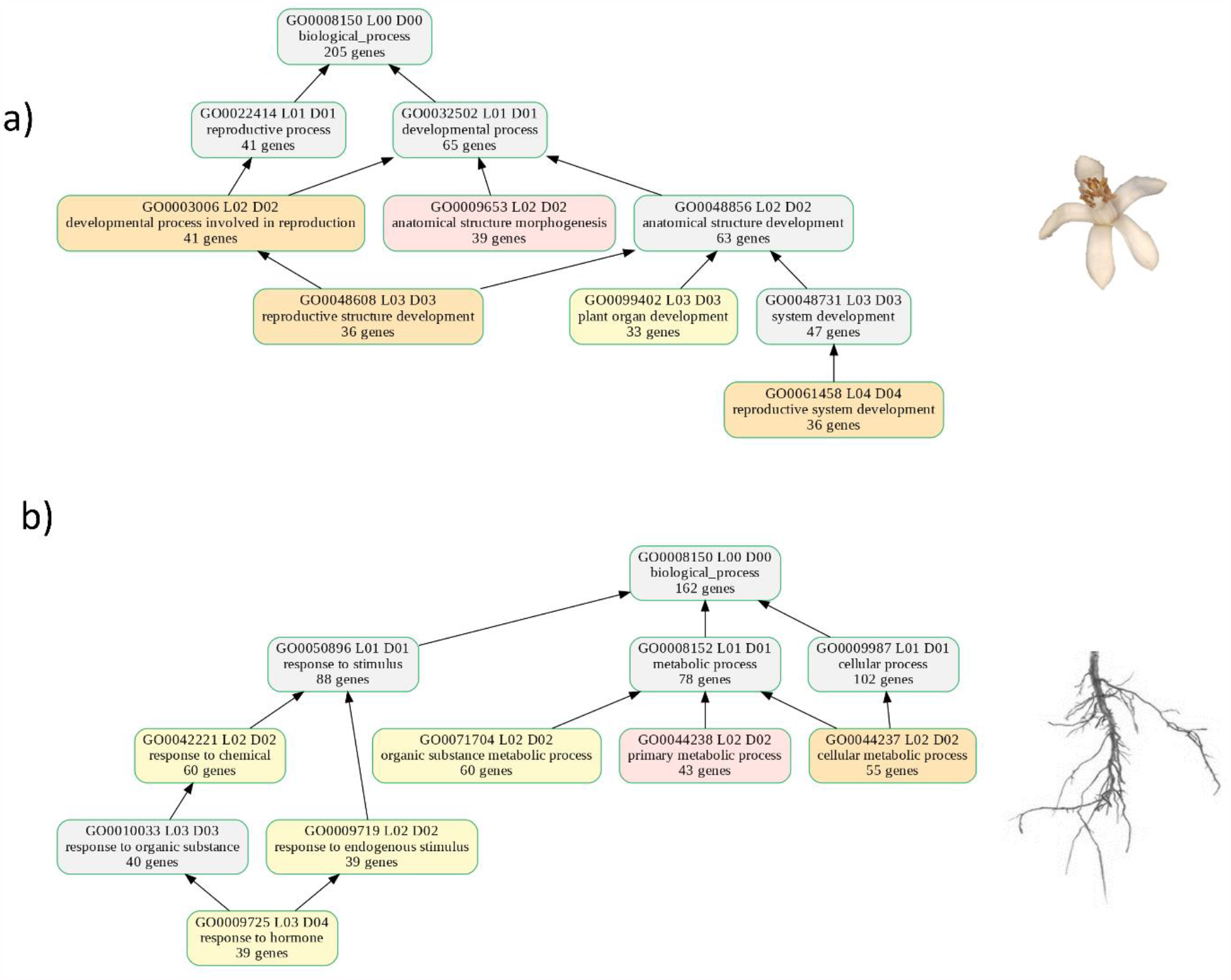
Enriched GO terms for the genes involved in ‘biological process’ and mapped in the alternative assembly in flower **(A)** and root **(B)**. The different color indicates different p-values, such as light red corresponds to a p-value < 0.005, light orange corresponds to a p-value < 0.01 and yellow corresponds to a p-value < 0.05. Small enrichments are created using only the best significant GO terms.

## 4 Conclusions

Here we present the first reference genome of lemon. Sequencing has been carried out combining short (Illumina) and long (Oxford Nanopore) sequencing approaches while genomic annotation was based on the analysis of four tissues to provide a comprehensive overview of the genes expressed in lemon. The high-quality draft genome will represent a milestone toward the identification of the genetic determinism of traits of agronomical interest and the set-up of marker-assisted breeding effort to select novel lemon selections characterized by superior characteristics (i.e., enhanced fruit quality and/or resistance to severe disease such as mal secco) (Catalano et al., 2021).

## Supporting information

all suplementary files

## 6 Author Contributions

**M. Di Guardo:** Conceptualization, Investigation, Writing - original draft, **Moretto M**.: Conceptualization, Formal analysis, Investigation, Writing - original draft. **Moser M**.: Conceptualization, Formal analysis, Investigation, Writing - original draft**; Catalano C**.: Investigation, **Troggio M**.: Conceptualization, **Deng Z**.: Investigation, **Cestaro A:** Investigation **Caruso M**, Investigation, Writing - review & editing, **Distefano G**.: Investigation, Writing - review & editing, Funding acquisition; **La Malfa S**.: Conceptualization, Writing - review & editing, Funding acquisition; **Bianco L**.: Conceptualization, Formal analysis, Investigation, Writing - original draft, **Gentile A**.: Conceptualization, Writing - review & editing, Funding acquisition

## 8 Data Availability Statement

The genome assembly sequences, and gene predictions will be submitted to the citrus genome database. Raw data have been submitted to NCBI’s SRA under the bioproject id PRJNA732837.

