## Supplementary material for "The haplotype-resolved reference genome of lemon (*Citrus limon* L. Burm f.)": all suplementary files: Supplementary_Figure.pdf

**Supplementary Figure 1.** Relative length distribution for the Oxford Nanopore (ONT) reads.

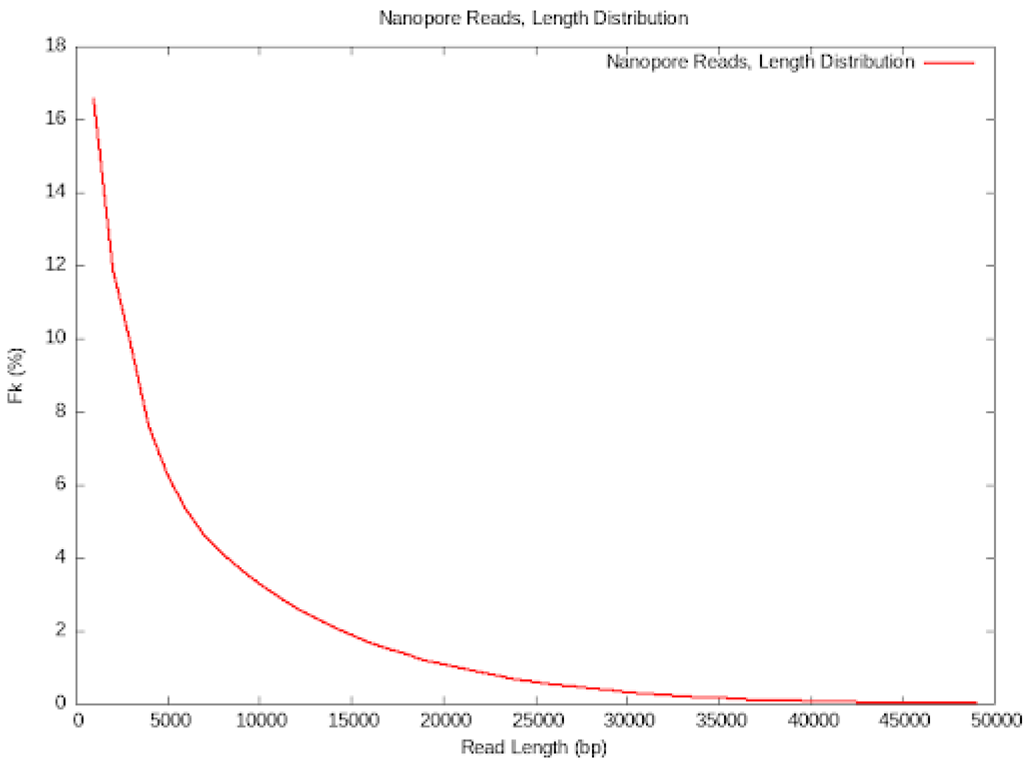

**Supplementary Figure 2.** GenomeScope K-mer profile plot (x axis = coverage of the K-mer, y axis = k-mer counts, for k-mer length = 21) of the Illumina sequencing of the lemon genome. Black line (full model) showed the fit of the GenomeScope model to the observed k-mer frequencies (blue, observed)

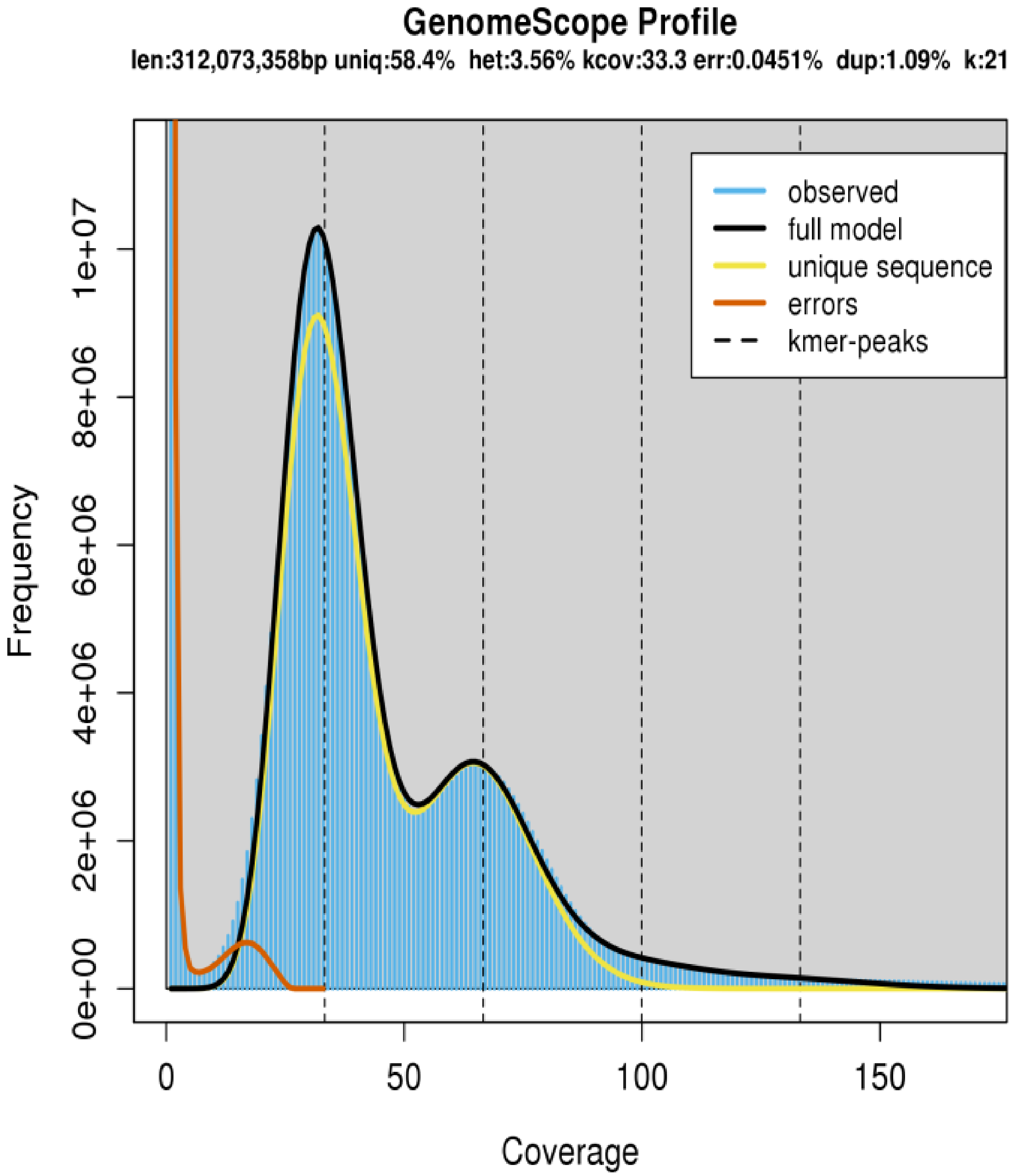

**Supplementary Figure 3.** Synteny analysis of the nine chromosomes of the assembled *Citrus limon* genome (X axis, (A): primary assembly, (B): alternative assembly) versus the reference genome of *Citrus maxima* (Y axis).

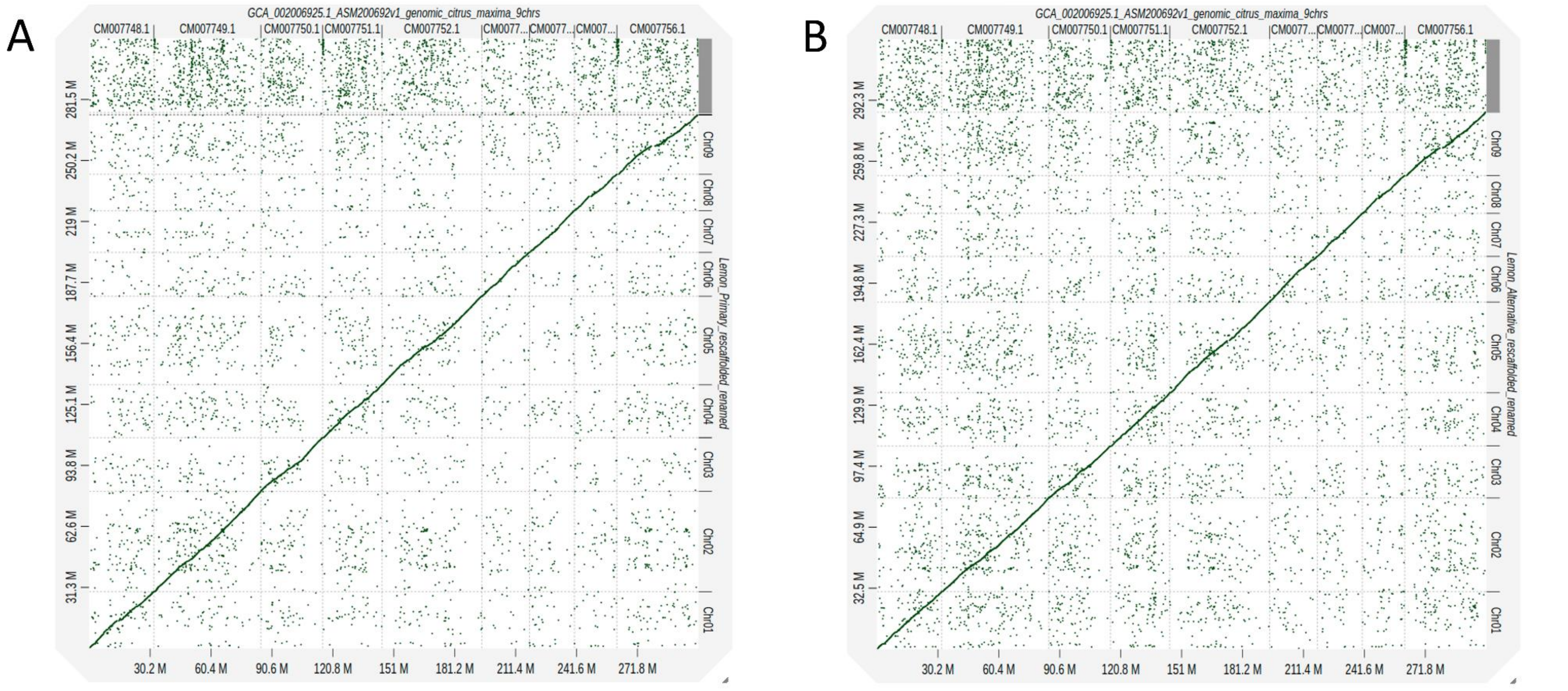

**Supplementary Figure 4. Bar-plot representing the distribution of main GO categories between the two haplotypes (colors are specified in legend).**

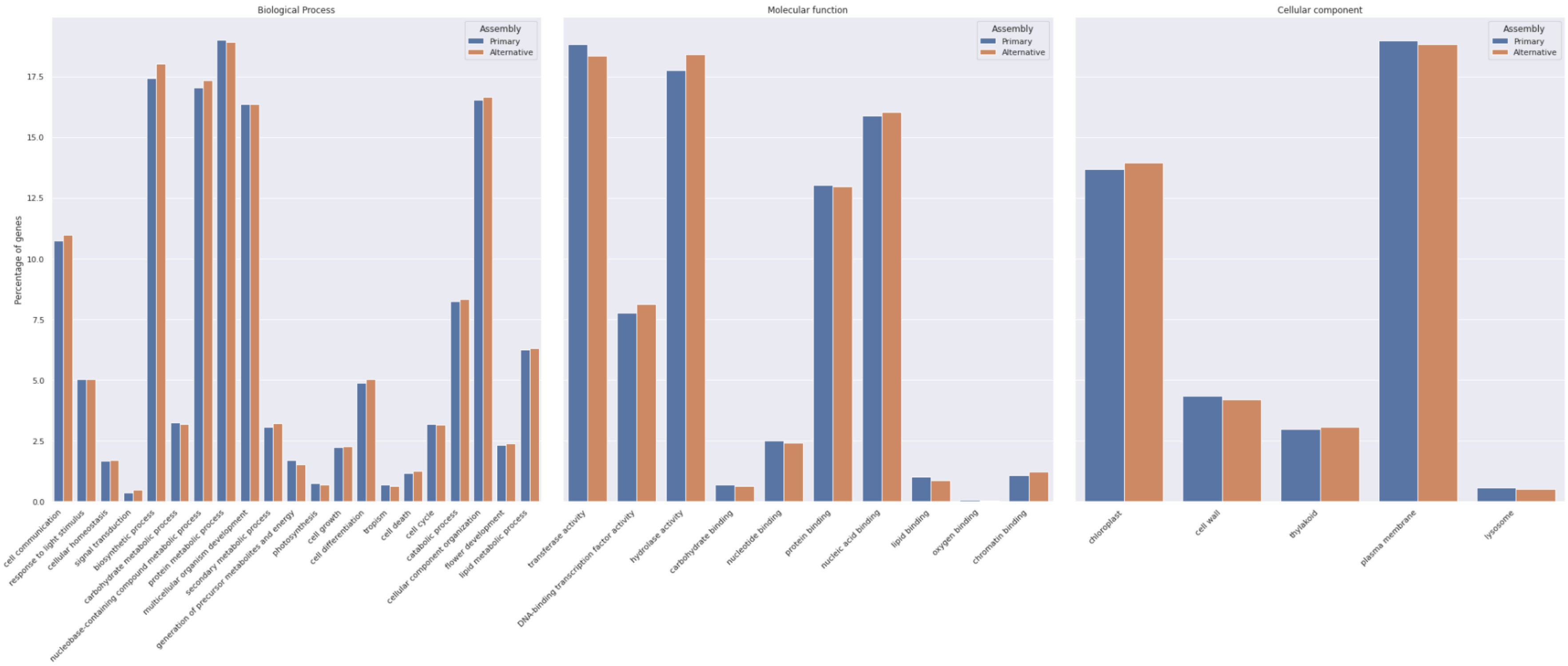

**Supplementary Figure 5.** Enriched GO terms for the genes involved in the three GO aspects (molecular function, biological process, cellular component) showing at least one GO term showing statistical significance. Data were depicted both for the primary and alternative assembly.

Tissue: **Root**, Assembly: **Primary**, GO: **Molecular Function**

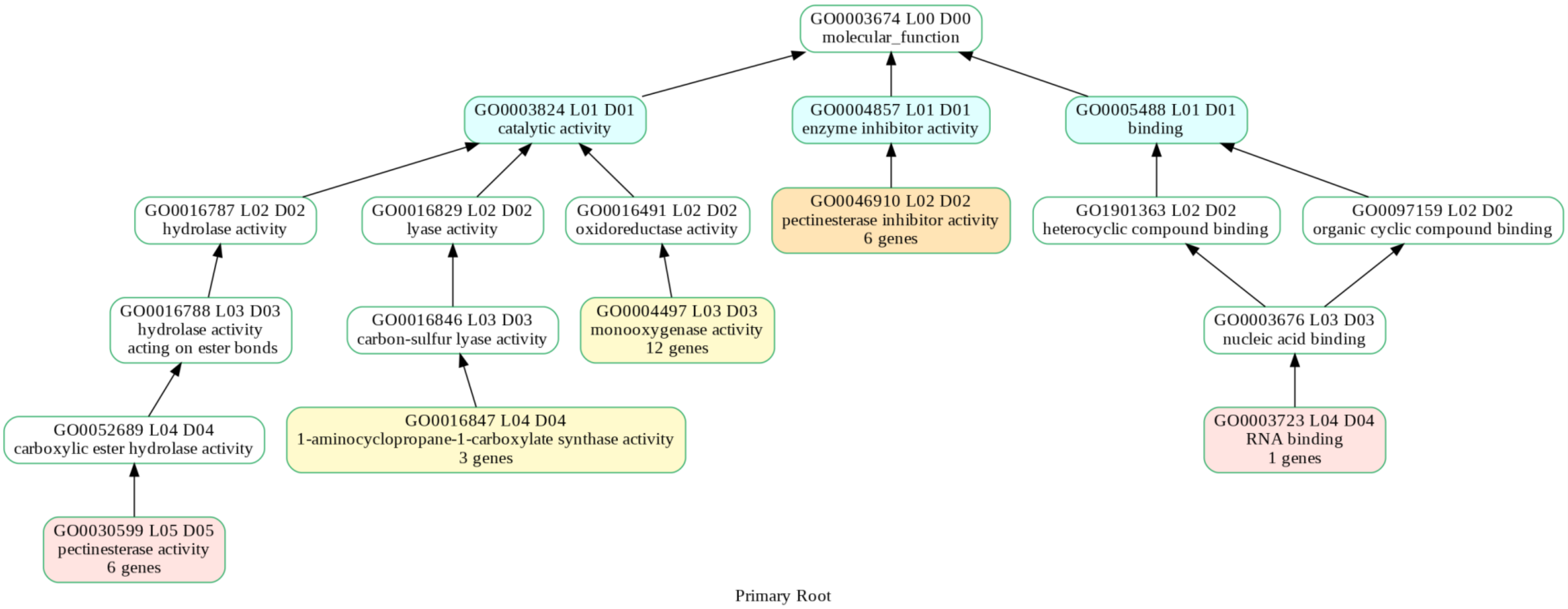

Tissue: **Root**, Assembly: **Primary**, GO: **Cellular Component**

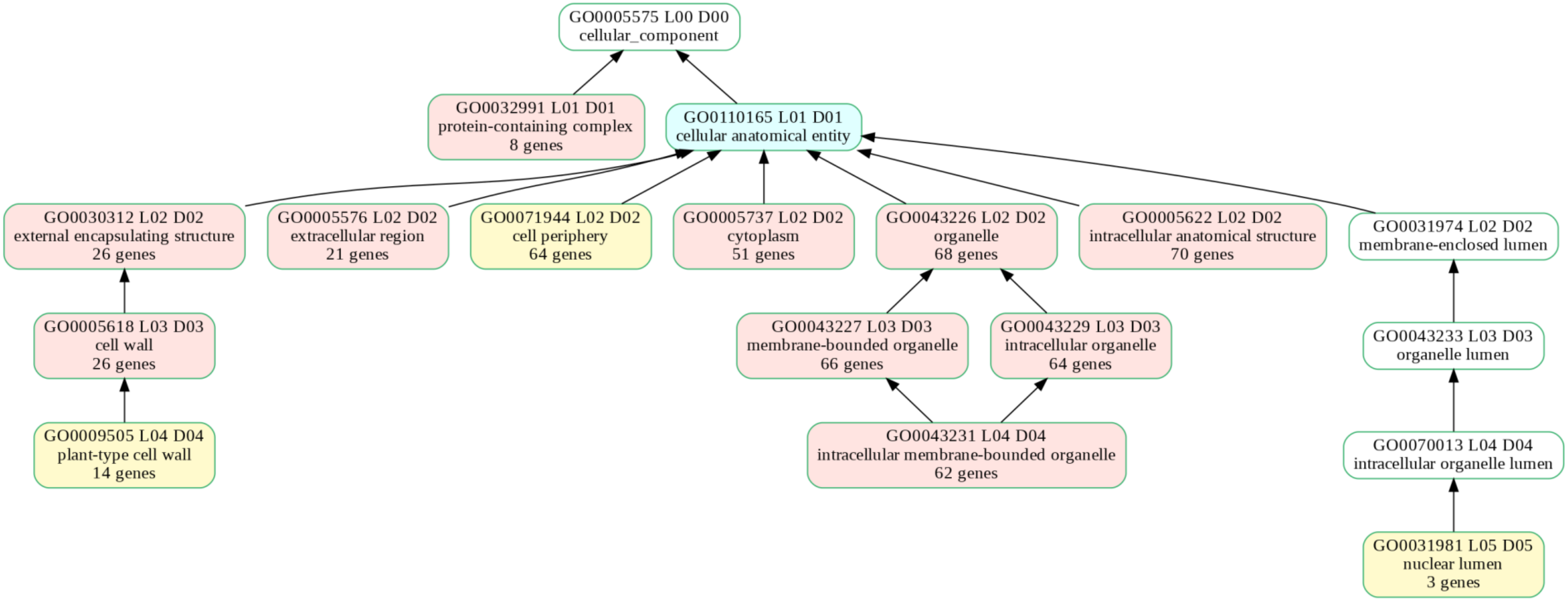

Primary Root

Tissue: **Root**, Assembly: **Primary**, GO: **Biological process**

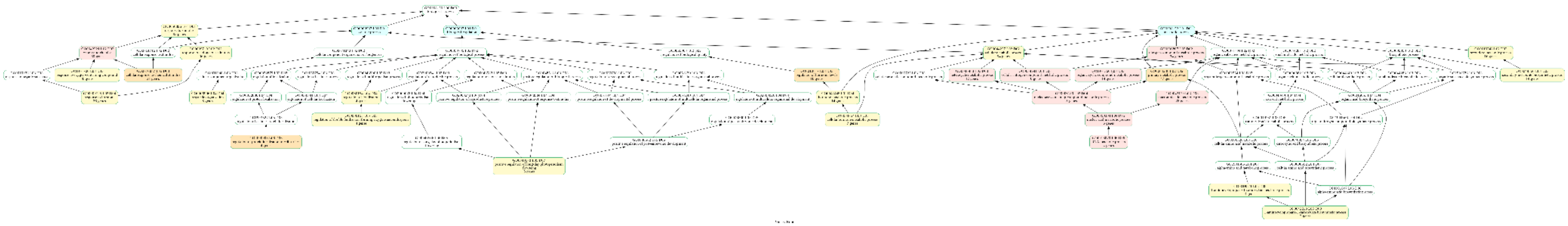

Tissue: **Fruit**, Assembly: **Primary**, GO: **Biological process**

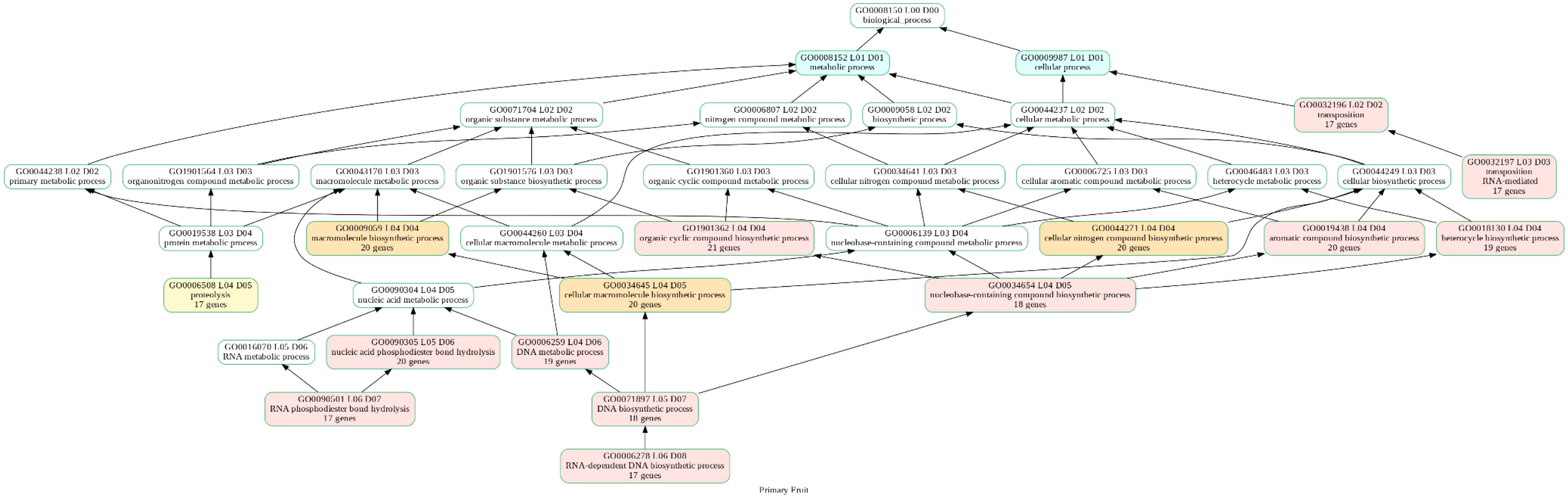

Tissue: **Fruit**, Assembly: **Primary**, GO: **Cellular Component**

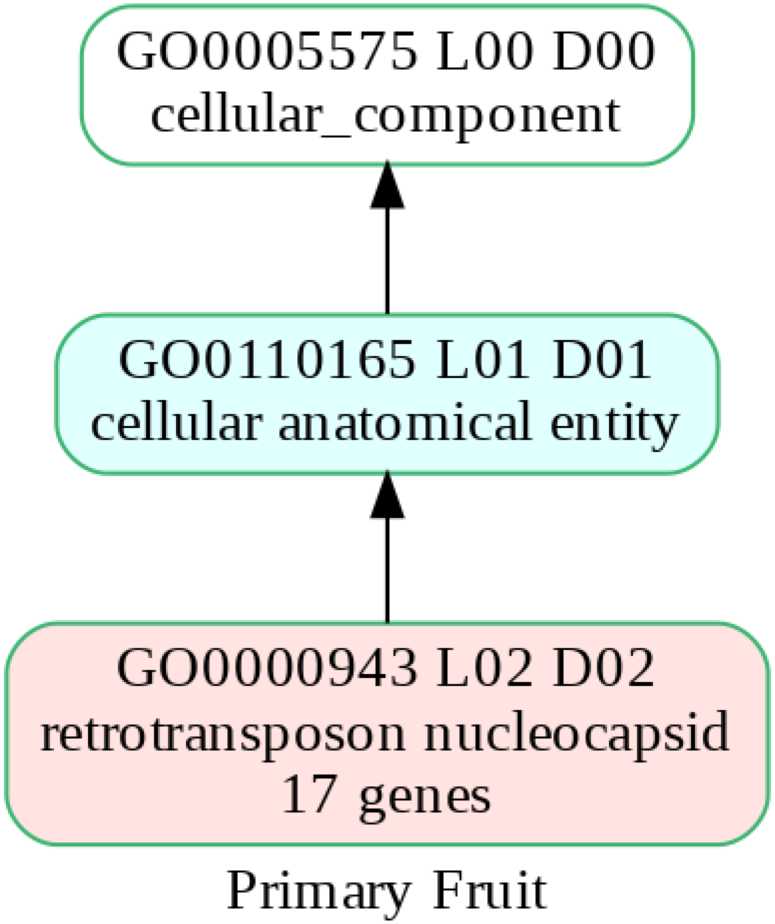

Tissue: **Fruit**, Assembly: **Primary**, GO: **Molecular Function**

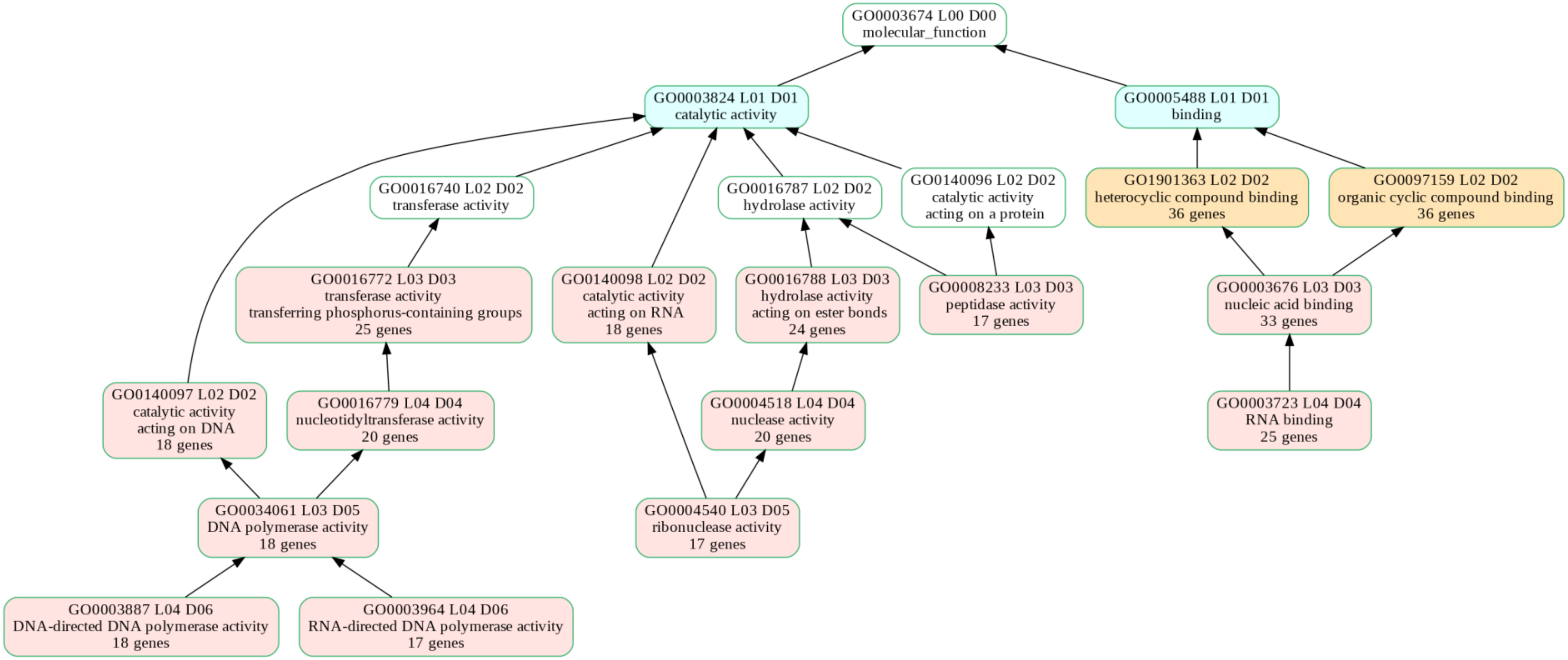


Primary Fruit

Tissue: **Flower**, Assembly: **Primary**, GO: **Biological process**

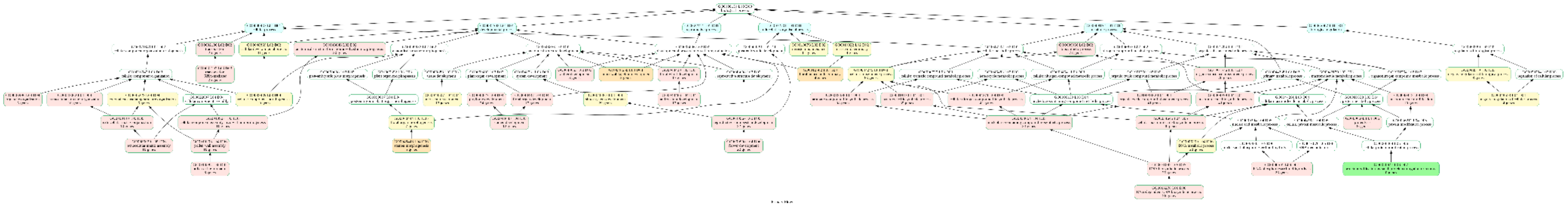

Tissue: **Flower**, Assembly: **Primary**, GO: **Cellular Component**

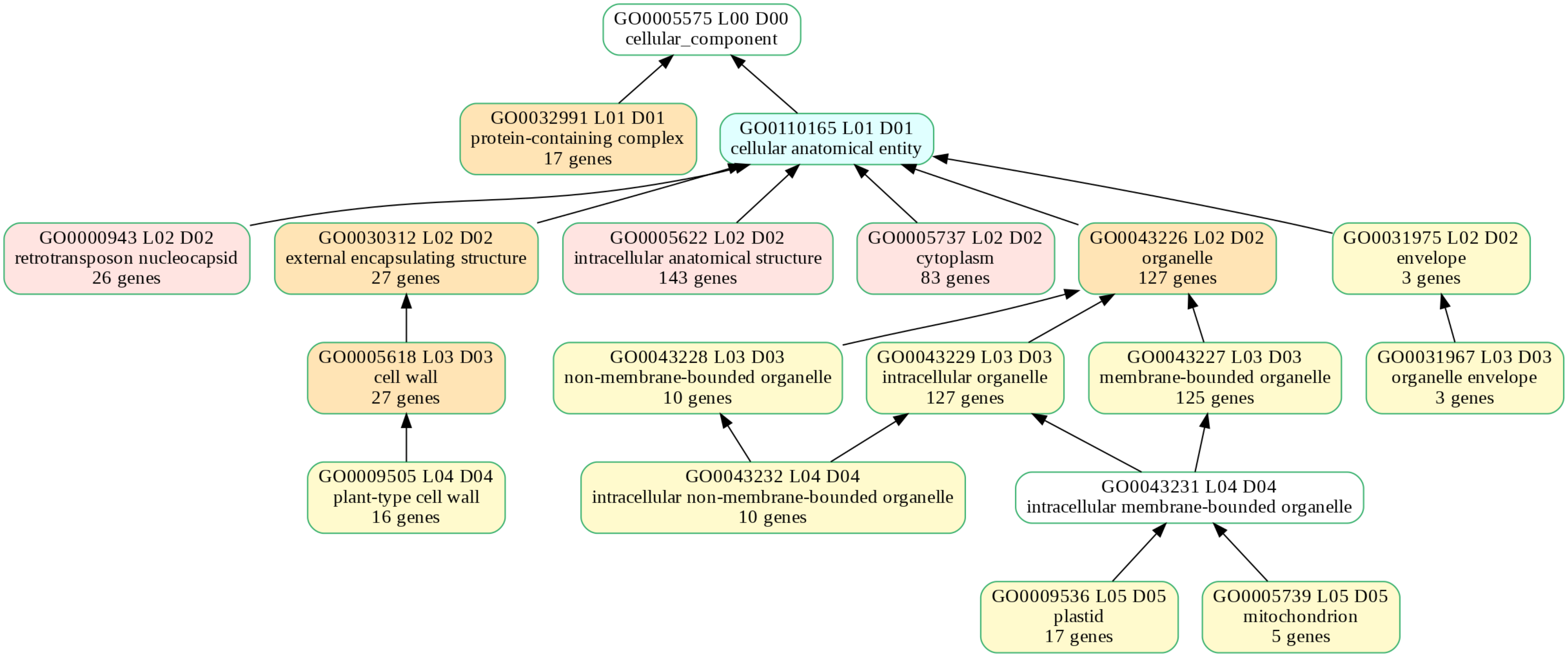


Primary Flower

Tissue: **Flower**, Assembly: **Primary**, GO: **Molecular Function**

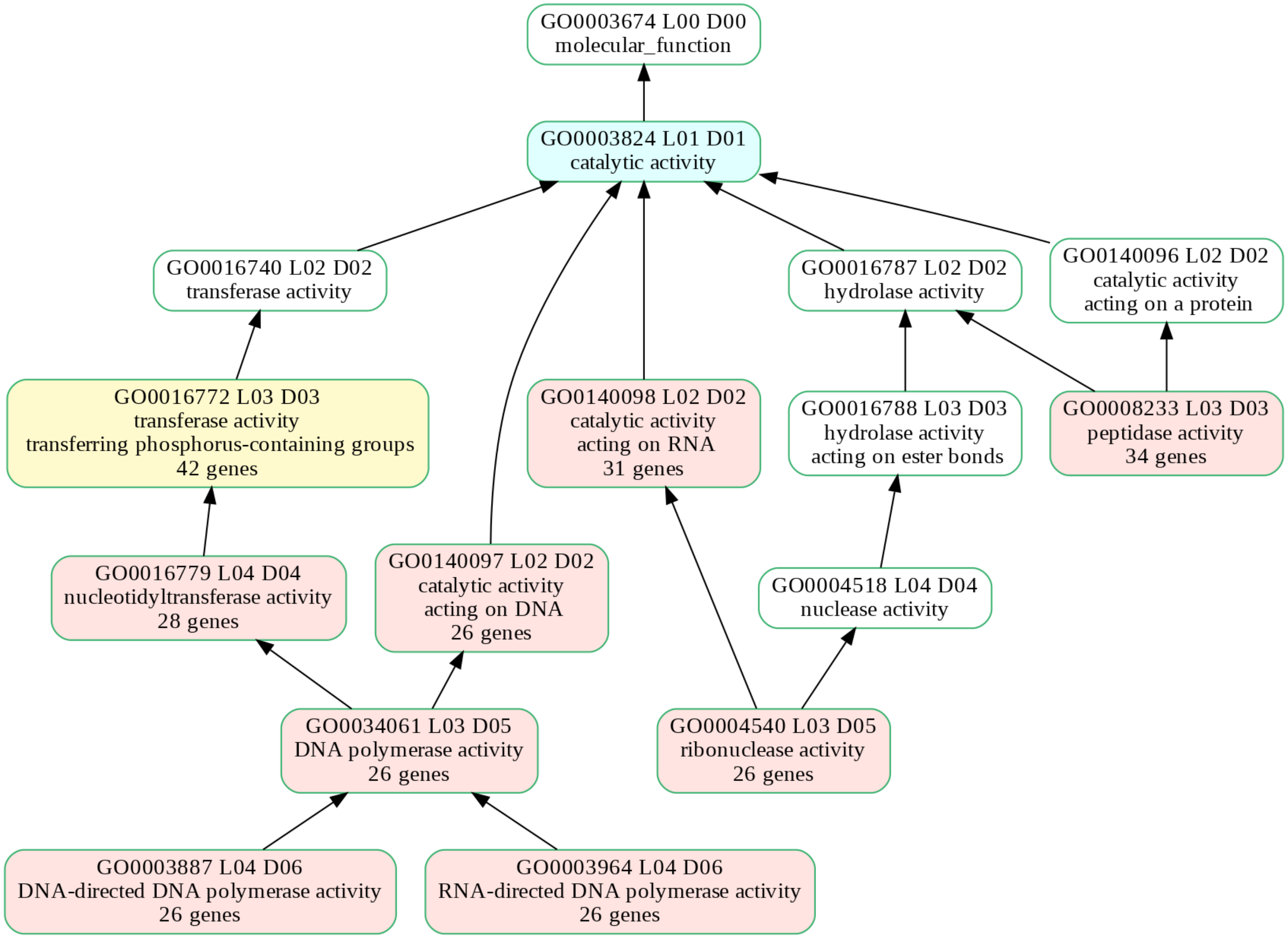

Primary Flower

Tissue: **Root**, Assembly: **Alternative**, GO: **Biological process**

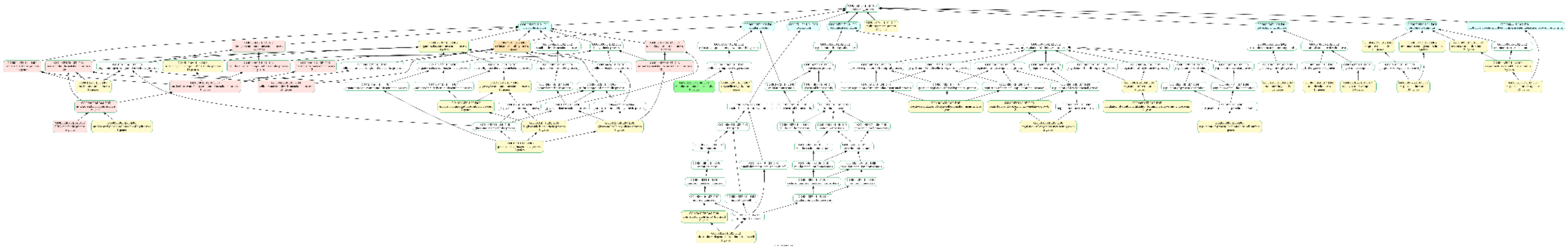

Tissue: **Root**, Assembly: **Alternative**, GO: **Cellular Component**

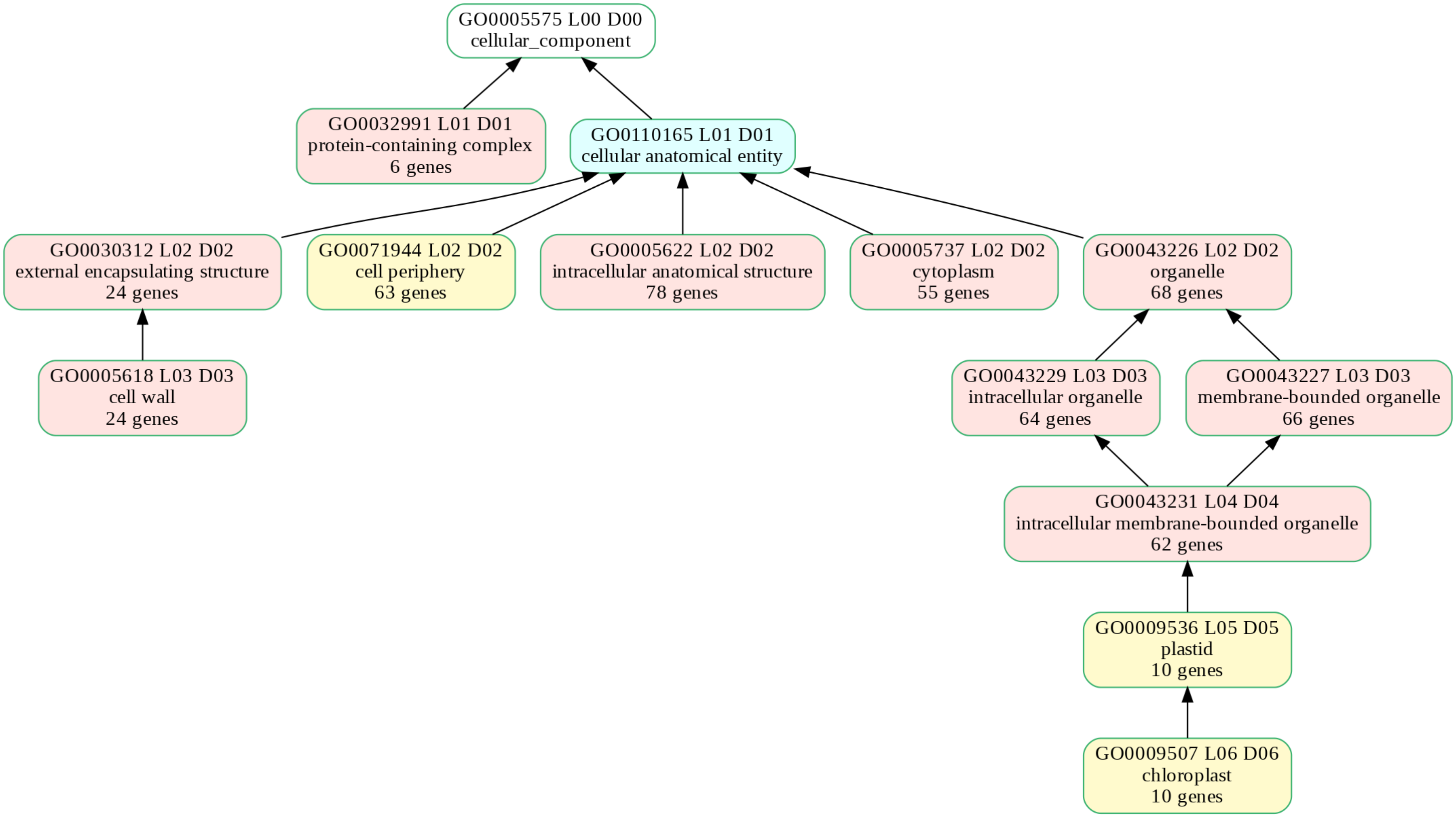


Alternative Root

Tissue: **Root**, Assembly: **Alternative**, GO: **Molecular Function**

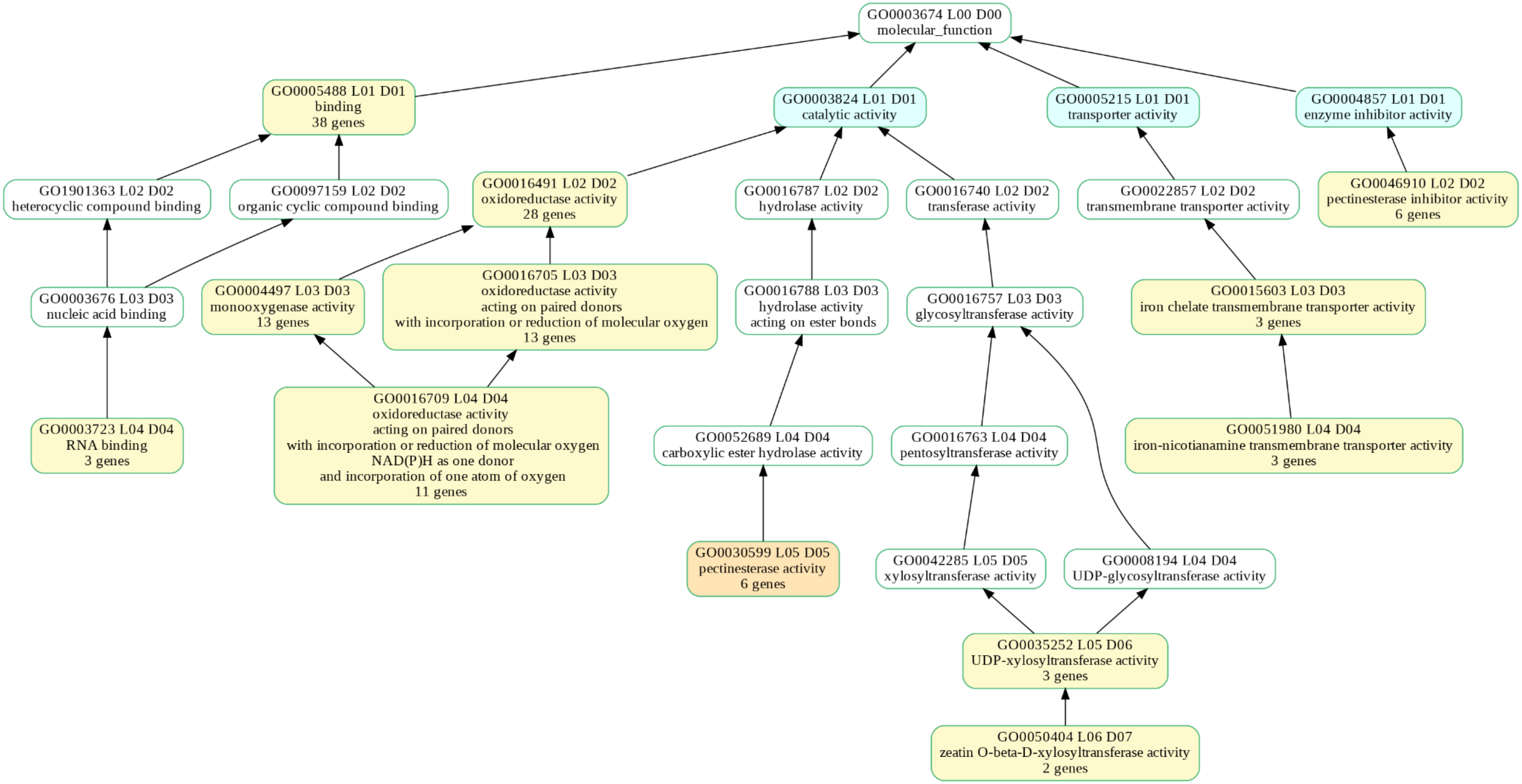


Alternative Root

Tissue: **Fruit**, Assembly: **Alternative**, GO: **Biological process**

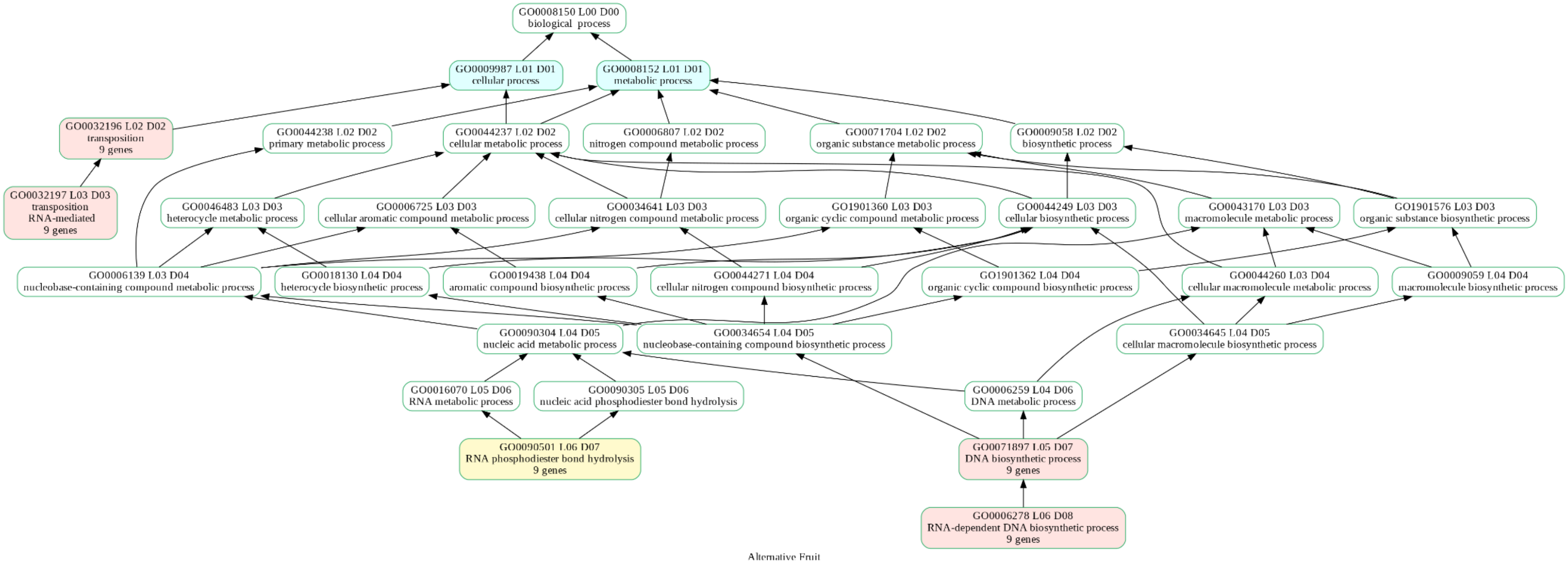

Tissue: **Fruit**, Assembly: **Alternative**, GO: **Cellular Component**

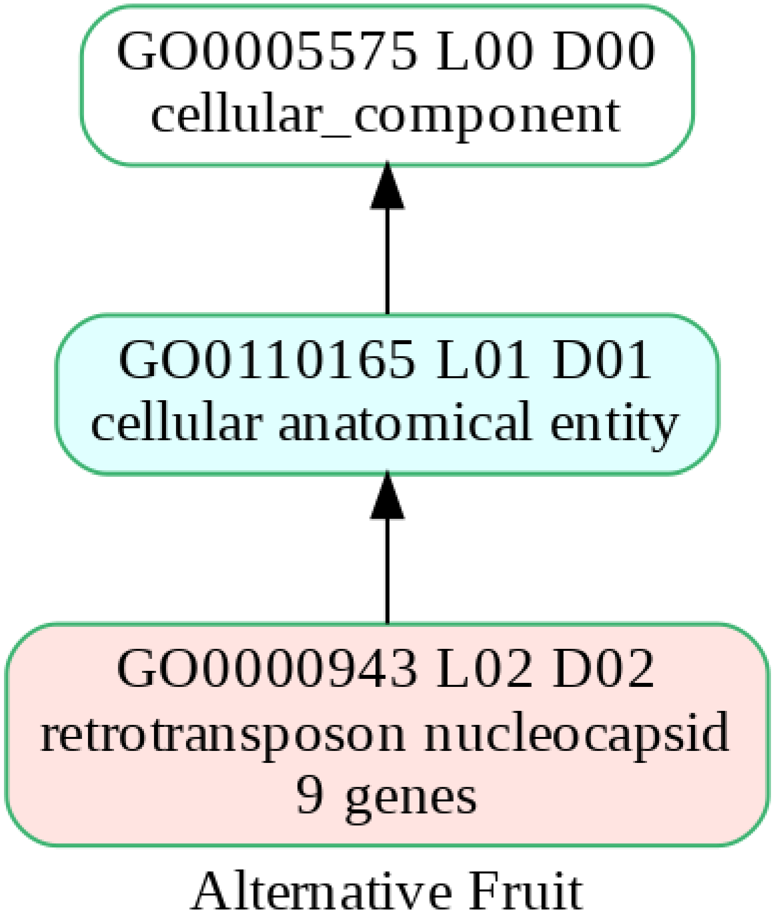

Tissue: **Fruit**, Assembly: **Alternative**, GO: **Molecular Function**

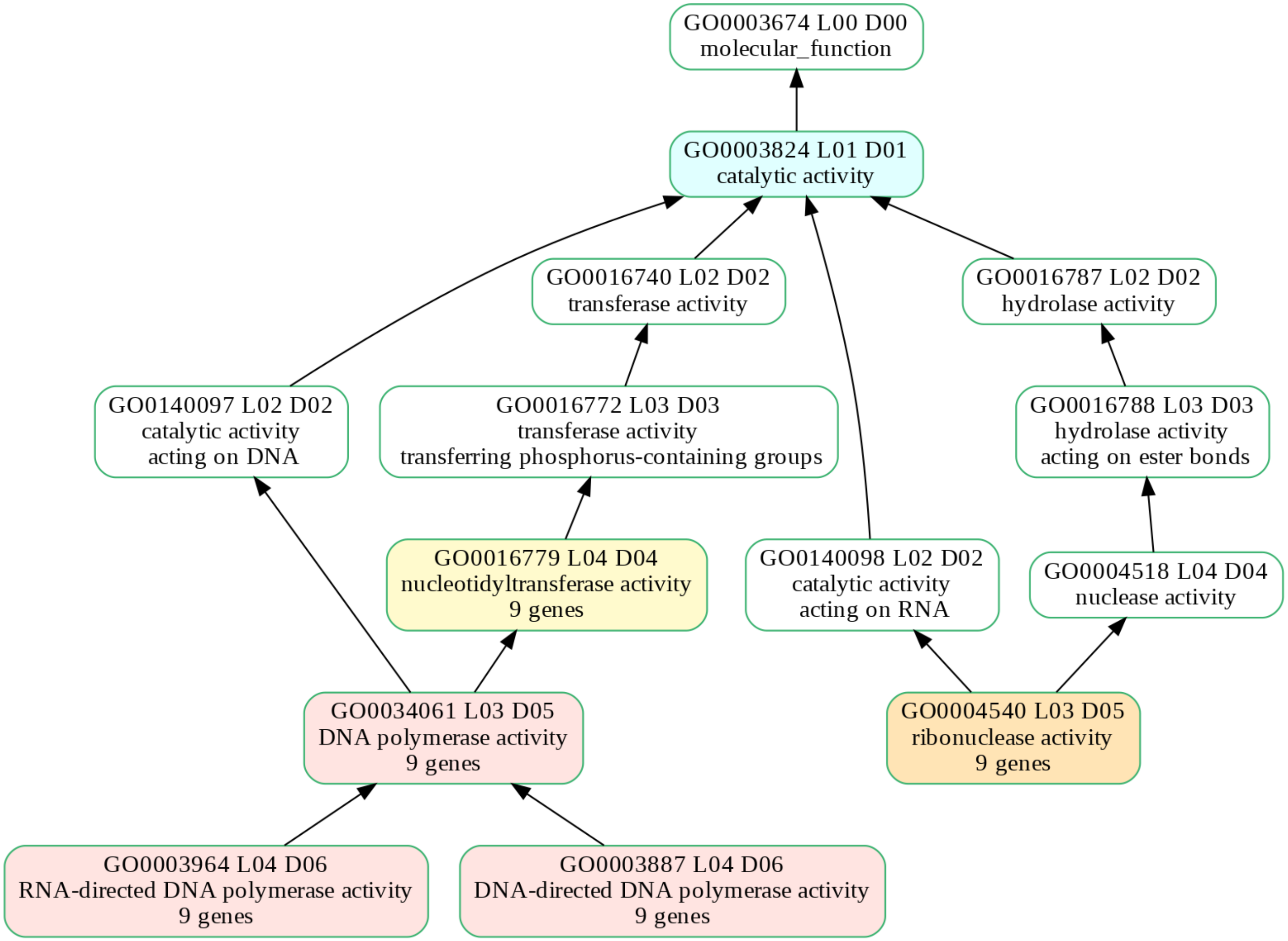


Alternative Fruit

Tissue: **Flower**, Assembly: **Alternative**, GO: **Biological process**

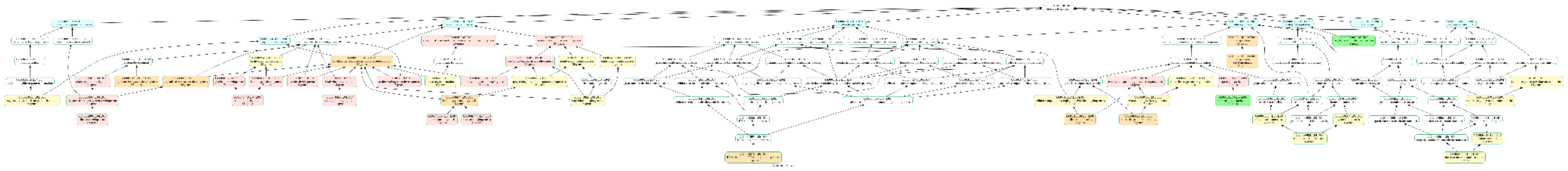

Tissue: **Flower**, Assembly: **Alternative**, GO: **Cellular Component**

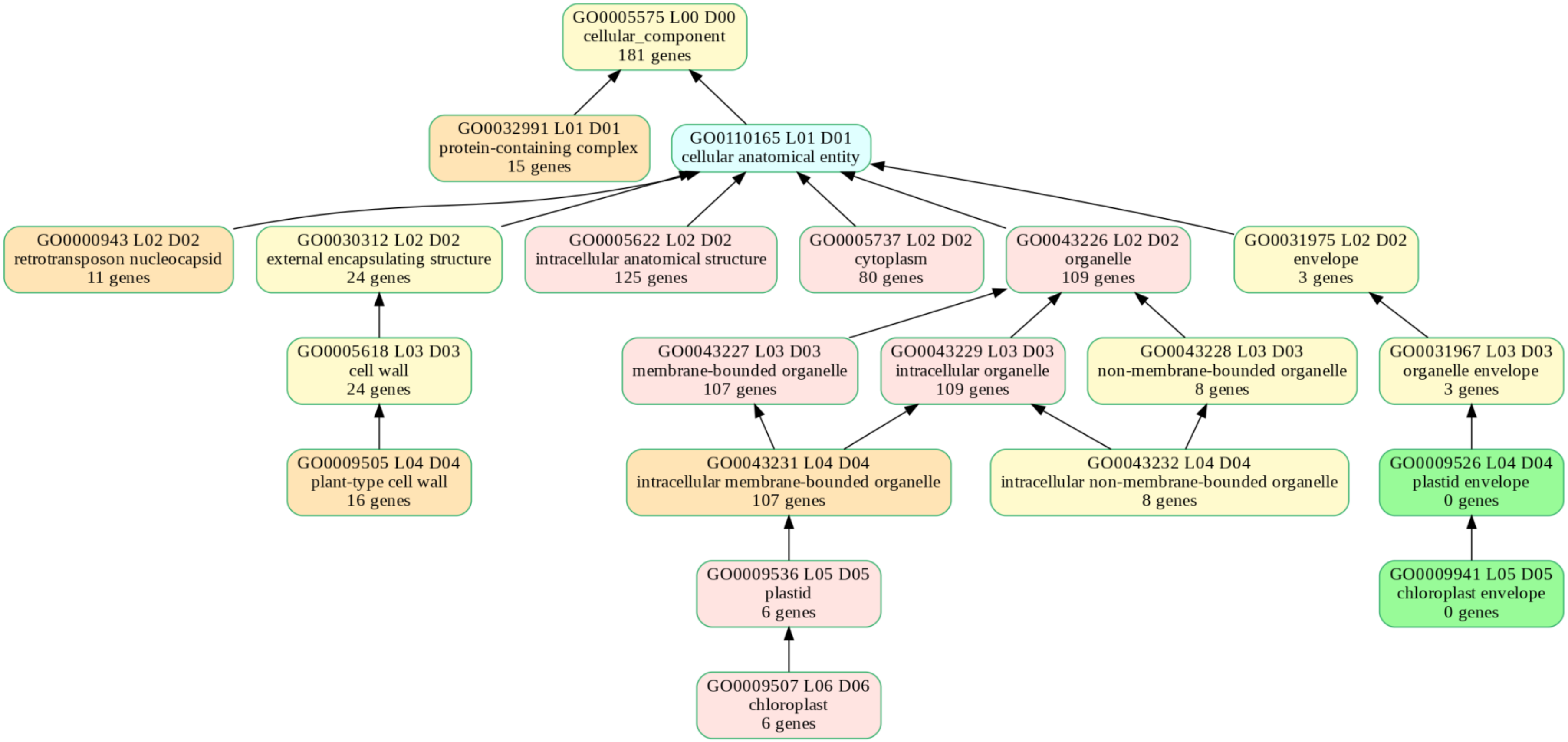


Alternative Flower

Tissue: **Flower**, Assembly: **Alternative**, GO: **Molecular Function**

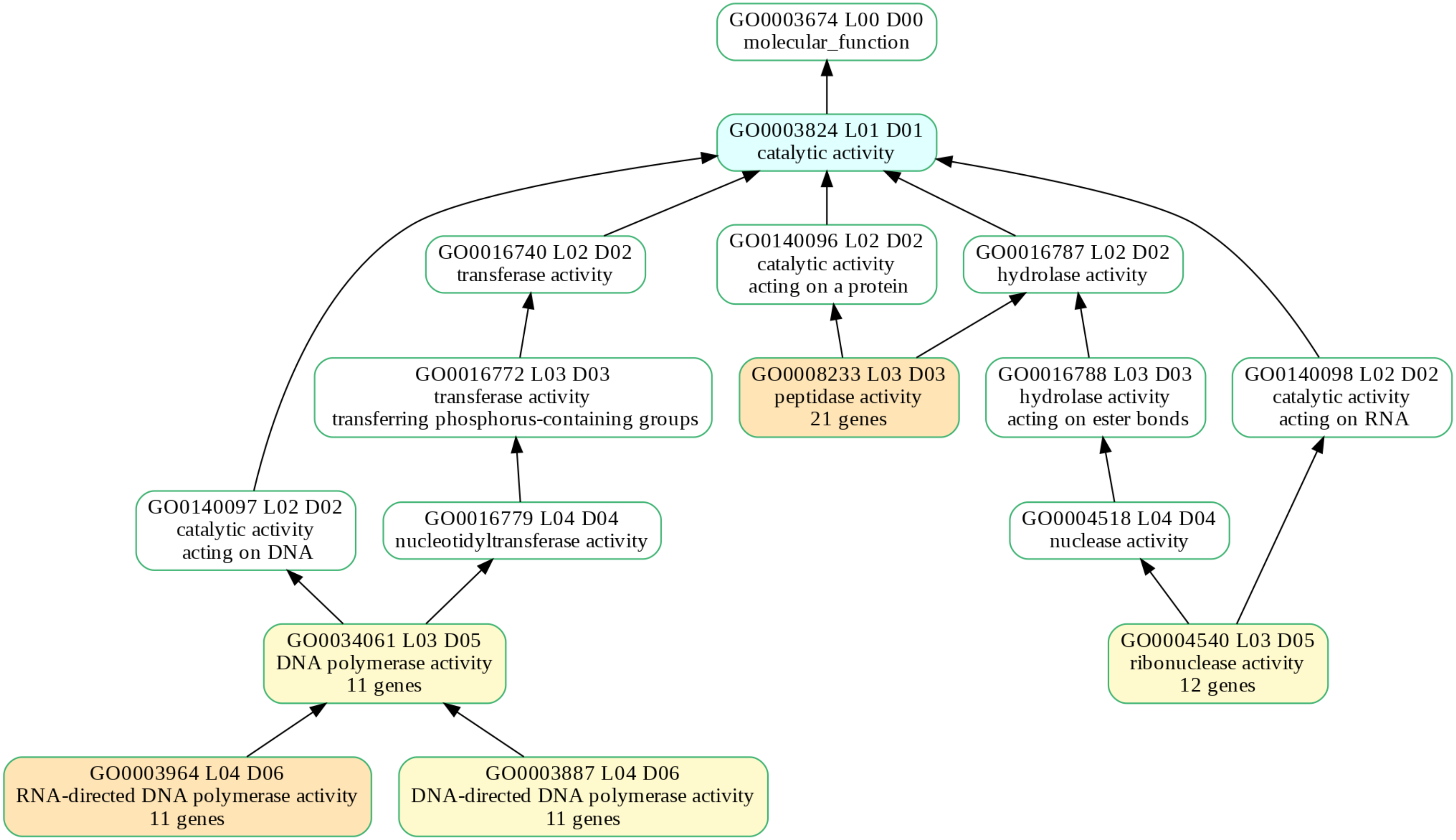


Alternative Flower
