## Supplementary material for "The haplotype-resolved reference genome of lemon (*Citrus limon* L. Burm f.)": all suplementary files: supplementary_tables.pdf

**Supplementary table 1: Coordinates of the miRNA that were found with high confidence level**

| Primary | | | | | | Alternative | | | | | |
|---|---|---|---|---|---|---|---|---|---|---|---|
| **miRNA mature** | **Chromosome** | **5p coordinates (start-end)** | **3p coordinates (start-end)** | **Hairpin coordinates** | **Strand** | **miRNA mature** | **Chromosome** | **5p coordinates (start-end)** | **3p coordinates (start-end)** | **Hairpin coordinates** | **Strand** |
| miR12105 | Chr06 | 14770633-14770655 | 14770520-14770541 | 14770665-14770464 | (-) | miR12105 | Chr06 | 16818362-16818384 | 16818249-16818270 | 16818394-16818193 | (-) |
| miR12106 | Chr02 | 47586890-47586911 | 47586830-47586850 | 47586921-47586822 | (-) | miR12106 | Chr02 | 45583012-45583033 | 45582952-45582972 | 45583043-45582944 | (-) |
| miR12109 | Chr07 | 10387966-10387989 | 10388003-10388026 | 10387936-10388056 | (+) | | | | | | |
| miR12110 | Chr09 | 29247066-29247086 | 29247123-29247144 | 29247053-29247156 | (+) | miR12110 | Chr09 | 32140712-32140732 | 32140769-32140790 | 32140699-32140802 | (+) |
| miR12111 | Chr05 | 8622192-8622215 | 8622101-8622124 | 8622237-8622088 | (-) | miR12111 | Chr05 | 9071905-9071928 | 9071814-9071837 | 9071950-9071801 | (-) |
| miR1515b | Chr01 | 4817119-4817140 | 4817247-4817267 | 4817109-4817283 | (+) | miR1515b | Chr01 | 4080686-4080707 | 4080814-4080834 | 4080676-4080850 | (+) |
| miR156a | Chr04 | 26706114-26706133 | 26706053-26706074 | 26706145-26706043 | (-) | | | | | | |
| miR156b | Chr04 | 1158594-1158613 | 1158530-1158552 | 1158643-1158501 | (-) | miR156b | Chr04 | 1201991-1202010 | 1201927-1201949 | 1202040-1201898 | (-) |
| miR156c | Chr04 | 21390625-21390644 | 21390556-21390576 | 21390674-21390525 | (-) | miR156c | Chr04 | 22411008-22411027 | 22410939-22410959 | 22411057-22410909 | (-) |
| miR156e | Chr04 | 13135928-13135948 | 13135993-13136014 | 13135900-13136044 | (+) | miR156e | Chr04 | 12797852-12797872 | 12797917-12797938 | 12797824-12797968 | (+) |
| miR156f | Chr04 | 16155151-16155171 | 16155215-16155236 | 16155121-16155266 | (+) | miR156f | Chr04 | 17213488-17213508 | 17213552-17213573 | 17213458-17213603 | (+) |
| miR156g | Chr06 | 16920503-16920523 | 16920564-16920584 | 16920473-16920614 | (+) | miR156g | Chr06 | 18961853-18961873 | 18961914-18961934 | 18961823-18961964 | (+) |
| | | | | | | miR156h | 001921F_alternative | 38948-38968 | | 38936-39036 | (+) |
| miR156j | Chr02 | 38391960-38391980 | 38392023-38392043 | 38391943-38392054 | (+) | miR156j | Chr02 | 36914697-36914717 | 36914760-36914780 | 36914680-36914791 | (+) |
| miR159a | Chr05 | 36825474-36825494 | 36825305-36825325 | 36825507-36825296 | (-) | miR159a | Chr05 | 39327063-39327083 | 39326894-39326914 | 39327096-39326885 | (-) |
| miR159b | Chr01 | 22449995-22450015 | | 22450048-22449899 | (-) | miR159b | Chr01 | 23672648-23672668 | | 23672701-23672551 | (-) |
| miR159c | Chr05 | 1739136-1739155 | 1738989-1739009 | 1739165-1738977 | (-) | miR159c | Chr05 | 1038011-1038030 | 1037864-1037884 | 1038040-1037852 | (-) |
| miR159d | Chr04 | 5043038-5043058 | | 5042949-5043071 | (+) | miR159d | Chr04 | 5235961-5235981 | | 5235872-5235994 | (+) |
| miR160a | Chr07 | 4135473-4135493 | 4135413-4135433 | 4135526-4135384 | (-) | miR160a | Chr07 | 5601541-5601561 | 5601481-5601501 | 5601593-5601452 | (-) |
| miR160b | Chr02 | 48982933-48982953 | 48982995-48983015 | 48982903-48983045 | (+) | miR160b | Chr02 | 47001148-47001168 | 47001210-47001230 | 47001118-47001260 | (+) |

| | | | | | | | | | | | |
|---|---|---|---|---|---|---|---|---|---|---|---|
| miR160c | Chr06 | 3956646-3956666 | 3956707-3956727 | 3956634-3956737 | (+) | miR160c | Chr06 | 4553905-4553925 | 4553966-4553986 | 4553893-4553996 | (+) |
| miR162 | Chr04 | 18261814-18261835 | 18261744-18261764 | 18261924-18261724 | (-) | miR162 | 001535F_alternative | 85217-85238 | 85147-85167 | 85327-85127 | (-) |
| miR164a | Chr06 | 12868956-12868976 | 12869105-12869125 | 12868943-12869181 | (+) | miR164a | Chr06 | 14939436-14939456 | 14939585-14939605 | 14939423-14939661 | (+) |
| miR164b | Chr05 | 40006690-40006710 | 40006620-40006640 | 40006740-40006590 | (-) | miR164b | Chr05 | 42797010-42797030 | 42796940-42796960 | 42797060-42796910 | (-) |
| miR164c | Chr02 | 40326920-40326940 | 40326859-40326879 | 40326970-40326829 | (-) | miR164c | Chr02 | 38768703-38768723 | 38768642-38768662 | 38768753-38768612 | (-) |
| miR164d | Chr05 | 40006690-40006710 | 40006620-40006640 | 40006740-40006590 | (-) | miR164d | Chr05 | 42797010-42797030 | 42796940-42796960 | 42797060-42796910 | (-) |
| miR166b | Chr03 | 7800495-7800516 | 7800390-7800410 | 7800526-7800380 | (-) | miR166b | 000909F_alternative | 27102-27123 | 26997-27017 | 27133-26987 | (-) |
| miR166c | Chr05 | 37646940-37646960 | 37647009-37647028 | 37646910-37647047 | (+) | | | | | | |
| | | | | | | miR166d | Chr05 | 20076385-20076402 | 20076463-20076483 | 20076368-20076495 | (+) |
| miR166e | Chr05 | 41933710-41933730 | 41933593-41933613 | 41933751-41933575 | (-) | | | | | | |
| miR166f | Chr03 | 7800637-7800657 | 7800565-7800585 | 7800687-7800535 | (-) | miR166f | 000909F_alternative | 27244-27264 | 27172-27192 | 27294-27142 | (-) |
| miR166g | Chr09 | 5636101-5636121 | 5636046-5636066 | 5636147-5636016 | (-) | miR166g | Chr09 | 5265086-5265106 | 5265031-5265051 | 5265134-5265001 | (-) |
| miR167a | Chr03 | 15160574-15160594 | 15160519-15160539 | 15160608-15160506 | (-) | miR167a | Chr03 | 16208351-16208371 | 16208296-16208316 | 16208385-16208283 | (-) |
| miR167b | Chr09 | 4927616-4927636 | 4927667-4927687 | 4927597-4927719 | (+) | | | | | | |
| miR167c | Chr02 | 49413443-49413463 | 49413741-49413761 | 49413433-49413769 | (+) | miR167c | Chr02 | 47653245-47653265 | 47653541-47653561 | 47653235-47653569 | (+) |
| miR167d | Chr05 | 38031842-38031862 | 38031794-38031817 | 38031872-38031786 | (-) | miR167d | Chr05 | 40778019-40778039 | 40777971-40777994 | 40778049-40777963 | (-) |
| miR167e | Chr05 | 38035167-38035187 | 38035081-38035103 | 38035217-38035051 | (-) | miR167e | Chr05 | 40781470-40781490 | 40781384-40781406 | 40781520-40781354 | (-) |
| miR168 | Chr03 | 14707579-14707599 | 14707457-14707477 | 14707618-14707442 | (-) | | | | | | |
| miR169c | Chr02 | 8818042-8818061 | 8817967-8817987 | 8818091-8817937 | (-) | miR169c | Chr02 | 8768589-8768608 | 8768514-8768534 | 8768636-8768484 | (-) |
| miR169e | Chr02 | 8789744-8789763 | 8789665-8789685 | 8789793-8789635 | (-) | | | | | | |
| miR169f | Chr08 | 10235387-10235406 | | 10235357-10235529 | (+) | miR169f | Chr08 | 11524902-11524921 | | 11524872-11525044 | (+) |
| miR169g | Chr07 | 15140223-15140243 | | 15140273-15140068 | (-) | miR169g | Chr07 | 16485154-16485174 | | 16485204-16484999 | (-) |
| miR169h | Chr02 | 8789744-8789763 | | 8789793-8789633 | (-) | | | | | | |
| miR169j | Chr02 | 6294401-6294420 | | 6294450-6294298 | (-) | miR169j | Chr02 | 5934482-5934501 | | 5934531-5934379 | (-) |

| | | | | | | | | | | | |
|---|---|---|---|---|---|---|---|---|---|---|---|
| miR169k | Chr06 | 21707965-21707985 | | 21708018-21707837 | (-) | miR169k | Chr06 | 23321483-23321503 | | 23321536-23321355 | (-) |
| | | | | | | miR169l | Chr02 | 8792126-8792145 | 8792051-8792070 | 8792175-8792021 | (-) |
| miR169m | Chr02 | 96354-96374 | | 96404-96228 | (-) | miR169m | Chr02 | 150681-150701 | | 150731-150555 | (-) |
| miR169n | Chr04 | 285284-285304 | 285331-285351 | 285280-285358 | (+) | miR169n | Chr04 | 222992-223012 | 223039-223059 | 222988-223066 | (+) |
| miR169o | Chr04 | 285068-285088 | | 285065-285160 | (+) | miR169o | Chr04 | 222775-222795 | | 222772-222867 | (+) |
| miR169p | Chr02 | 8819885-8819904 | | 8819911-8819795 | (-) | miR169p | Chr02 | 8770434-8770453 | | 8770460-8770342 | (-) |
| miR169q | Chr08 | 10251608-10251627 | | 10251602-10251728 | (+) | miR169q | Chr08 | 11542396-11542415 | | 11542390-11542516 | (+) |
| miR169r | Chr04 | 285068-285088 | | 285065-285160 | (+) | miR169r | Chr04 | 222775-222795 | | 222772-222867 | (+) |
| miR171a | Chr02 | 19156042-19156061 | | 19155978-19156069 | (+) | miR171a | Chr02 | 18565071-18565090 | | 18565007-18565098 | (+) |
| miR171c | Chr07 | 9595649-9595669 | 9595706-9595726 | 9595619-9595756 | (+) | miR171c | Chr07 | 11261920-11261940 | 11261977-11261997 | 11261890-11262027 | (+) |
| miR171d | Chr03 | | 26490318-26490338 | 26490422-26490294 | (-) | miR171d | Chr03 | | 26659400-26659420 | 26659508-26659370 | (-) |
| miR171e | Chr02 | | 19049709-19049729 | 19049616-19049759 | (+) | miR171e | Chr02 | | 18443552-18443572 | 18443459-18443602 | (+) |
| miR171f | Chr01 | 543595-543615 | 543519-543539 | 543645-543489 | (-) | miR171f | Chr01 | 563297-563317 | 563221-563241 | 563347-563191 | (-) |
| miR171g | Chr02 | | 19049709-19049729 | 19049616-19049759 | (+) | miR171g | Chr02 | | 18443552-18443572 | 18443459-18443602 | (+) |
| miR171h | Chr02 | | 9784735-9784755 | 9784823-9784725 | (-) | miR171h | Chr02 | | 9543053-9543073 | 9543141-9543043 | (-) |
| miR171i | Chr02 | | 43676433-43676453 | 43676363-43676465 | (+) | miR171i | Chr02 | | 42282659-42282679 | 42282589-42282691 | (+) |
| miR172a | Chr01 | 8048733-8048753 | 8048817-8048836 | 8048703-8048864 | (+) | miR172a | Chr01 | 7128280-7128300 | 7128364-7128383 | 7128250-7128411 | (+) |
| miR172b | Chr07 | 3659514-3659534 | 3659618-3659638 | 3659484-3659668 | (+) | miR172b | Chr07 | 4695356-4695376 | 4695460-4695480 | 4695326-4695510 | (+) |
| miR172c | Chr07 | 418111-418131 | 418217-418238 | 418098-418261 | (+) | miR172c | Chr07 | 420168-420188 | 420274-420295 | 420155-420318 | (+) |
| miR172d | Chr08 | 12986480-12986500 | | 12986530-12986371 | (-) | miR172d | Chr08 | 14545386-14545406 | | 14545436-14545277 | (-) |
| miR2111 | Chr06 | 7635828-7635848 | | 7635864-7635757 | (-) | miR2111 | Chr06 | 9843847-9843867 | | 9843883-9843776 | (-) |
| miR2275b | Chr09 | 11793835-11793855 | 11793897-11793918 | 11793805-11793988 | (+) | miR2275b | Chr09 | 11431711-11431731 | 11431773-11431794 | 11431681-11431864 | (+) |
| miR3627a | Chr06 | 20772009-20772030 | | 20771979-20772128 | (+) | miR3627a | Chr06 | 22499716-22499737 | | 22499686-22499835 | (+) |
| miR3627b | Chr06 | | 20768200-20768221 | 20768380-20768116 | (-) | miR3627b | Chr06 | | 22495914-22495935 | 22496094-22495830 | (-) |

| | | | | | | | | | | | |
|---|---|---|---|---|---|---|---|---|---|---|---|
| miR3627c | Chr06 | 20773815-20773836 | | 20773866-20773744 | (-) | miR3627c | Chr06 | 22501885-22501906 | | 22501936-22501814 | (-) |
| miR390a | Chr06 | 12642381-12642401 | 12642459-12642479 | 12642361-12642505 | (+) | miR390a | Chr06 | 14717939-14717959 | 14718017-14718037 | 14717919-14718074 | (+) |
| miR390b | Chr08 | 11662500-11662519 | | 11662550-11662392 | (-) | miR390b | Chr08 | 12808926-12808945 | | 12808976-12808818 | (-) |
| miR391 | Chr08 | 14347549-14347570 | 14347606-14347627 | 14347519-14347659 | (+) | miR391 | 003911F_alternative | 9445-9466 | 9388-9409 | 9496-9356 | (-) |
| miR393a | Chr02 | 4703393-4703414 | | 4703374-4703518 | (+) | miR393a | Chr02 | 4419433-4419454 | | 4419414-4419557 | (+) |
| miR393b | Chr09 | 28137600-28137621 | 28137508-28137528 | 28137651-28137478 | (-) | miR393b | Chr09 | 31140856-31140877 | 31140764-31140784 | 31140907-31140734 | (-) |
| miR3947 | Chr03 | 19455163-19455185 | | 19455044-19455205 | (+) | miR3947 | Chr03 | 20206376-20206398 | | 20206254-20206418 | (+) |
| miR3949 | Chr03 | 25014331-25014352 | | 25014444-25014321 | (-) | miR3949 | Chr03 | 24917812-24917833 | | 24917925-24917802 | (-) |
| miR394a | Chr03 | 26540935-26540954 | | 26540919-26541065 | (+) | miR394a | Chr03 | 26731055-26731074 | | 26731039-26731185 | (+) |
| miR394b | Chr03 | 8768528-8768547 | | 8768518-8768619 | (+) | miR394b | Chr03 | 8100843-8100862 | | 8100833-8100934 | (+) |
| miR3951a | Chr02 | 24537598-24537618 | 24537511-24537530 | 24537638-24537491 | (-) | | | | | | |
| miR3953 | 001180F_primary | 48601-48621 | | 48691-48591 | (-) | miR3953 | Chr05 | 13481654-13481674 | | 13481744-13481644 | (-) |
| miR3954b | Chr09 | 5742904-5742925 | 5742810-5742830 | 5743015-5742783 | (-) | miR3954b | Chr09 | 5625106-5625127 | 5625012-5625032 | 5625217-5624982 | (-) |
| miR395b | Chr07 | 15523650-15523670 | 15523610-15523630 | 15523700-15523580 | (-) | miR395b | Chr07 | 16864209-16864229 | 16864169-16864189 | 16864259-16864143 | (-) |
| | | | | | | miR396a | Chr04 | 1908849-1908869 | 1908917-1908937 | 1908829-1908981 | (+) |
| miR396c | Chr01 | 14547799-14547818 | | 14547673-14547828 | (+) | miR396c | Chr01 | 14513264-14513283 | | 14513138-14513293 | (+) |
| miR396e | Chr07 | 20969647-20969666 | 20969736-20969756 | 20969617-20969786 | (+) | miR396e | Chr07 | 22246070-22246089 | 22246159-22246179 | 22246040-22246209 | (+) |
| miR396f | Chr04 | 21269140-21269160 | 21269049-21269069 | 21269190-21269019 | (-) | miR396f | Chr04 | 22339373-22339393 | 22339284-22339304 | 22339422-22339254 | (-) |
| | | | | | | miR397 | Chr02 | 44867127-44867147 | 44867055-44867075 | 44867164-44867010 | (-) |
| miR398a | Chr07 | 14838833-14838854 | 14838931-14838951 | 14838800-14839024 | (+) | miR398a | Chr07 | 16225766-16225787 | 16225865-16225885 | 16225733-16225958 | (+) |
| miR398b | Chr02 | 4516657-4516677 | | 4516766-4516645 | (-) | miR398b | Chr02 | 4195622-4195642 | | 4195734-4195610 | (-) |
| | | | | | | miR399b | Chr05 | 7697253-7697273 | 7697152-7697172 | 7697296-7697129 | (-) |
| miR399d | Chr09 | 26871280-26871300 | 26871367-26871387 | 26871268-26871397 | (+) | miR399d | Chr09 | 29874273-29874293 | 29874360-29874380 | 29874261-29874390 | (+) |
| miR403a | Chr05 | 40819165-40819185 | 40819070-40819090 | 40819189-40819060 | (-) | | | | | | |

| | | | | | | | | | | | |
|---|---|---|---|---|---|---|---|---|---|---|---|
| miR403b | Chr09 | 781165-781185 | 781226-781246 | 781152-781256 | (+) | miR403b | Chr09 | 764756-764776 | 764817-764837 | 764743-764847 | (+) |
| miR408 | Chr05 | 41173075-41173095 | 41173015-41173035 | 41173154-41173002 | (-) | miR408 | Chr05 | 43989890-43989910 | 43989830-43989850 | 43989968-43989817 | (-) |
| miR477a | Chr07 | 11488190-11488210 | 11488255-11488275 | 11488172-11488309 | (+) | miR477a | Chr07 | 13526976-13526996 | 13527041-13527061 | 13526958-13527095 | (+) |
| miR477b | Chr07 | 20319137-20319157 | 20319210-20319231 | 20319118-20319255 | (+) | miR477b | Chr07 | 21708683-21708703 | 21708756-21708777 | 21708664-21708801 | (+) |
| miR477c | Chr07 | 20319334-20319354 | 20319396-20319416 | 20319314-20319474 | (+) | miR477c | Chr07 | 21708880-21708900 | 21708942-21708962 | 21708860-21709020 | (+) |
| miR477d | Chr05 | 1943864-1943885 | 1943932-1943952 | 1943858-1943959 | (+) | miR477d | Chr05 | 1258335-1258356 | 1258403-1258423 | 1258329-1258430 | (+) |
| miR477e | Chr05 | 1944077-1944098 | 1944134-1944154 | 1944070-1944162 | (+) | miR477e | Chr05 | 1258562-1258583 | 1258619-1258639 | 1258555-1258647 | (+) |
| miR482a | Chr02 | 51273706-51273727 | 51273789-51273810 | 51273693-51273820 | (+) | miR482a | Chr02 | 49824951-49824972 | 49825034-49825055 | 49824938-49825065 | (+) |
| miR482b | Chr02 | 51264729-51264749 | 51264793-51264814 | 51264716-51264824 | (+) | miR482b | Chr02 | 49812850-49812870 | 49812914-49812935 | 49812837-49812945 | (+) |
| miR482c | Chr02 | 51259749-51259769 | 51259805-51259826 | 51259739-51259834 | (+) | miR482c | Chr02 | 49807699-49807719 | 49807755-49807776 | 49807689-49807784 | (+) |
| miR482d | Chr05 | 10926334-10926354 | 10926425-10926446 | 10926304-10926476 | (+) | miR482d | Chr05 | 11477091-11477111 | 11477182-11477203 | 11477061-11477233 | (+) |
| miR482e | Chr02 | 51259530-51259549 | 51259611-51259632 | 51259500-51259662 | (+) | miR482e | Chr02 | 49807478-49807497 | 49807559-49807580 | 49807448-49807610 | (+) |
| miR482f | Chr02 | 51293795-51293815 | 51293845-51293865 | 51293782-51293875 | (+) | miR482f | Chr02 | 49847161-49847181 | 49847211-49847231 | 49847148-49847241 | (+) |
| miR482g | Chr02 | 51273706-51273727 | 51273789-51273810 | 51273693-51273820 | (+) | miR482g | Chr02 | 49824951-49824972 | 49825034-49825055 | 49824938-49825065 | (+) |
| miR530a | Chr08 | 15375504-15375524 | 15375419-15375439 | 15375531-15375411 | (-) | miR530a | Chr08 | 16829114-16829134 | 16829029-16829049 | 16829141-16829021 | (-) |
| miR530b | Chr08 | 15391424-15391444 | 15391539-15391559 | 15391414-15391567 | (+) | miR530b | Chr08 | 16847599-16847619 | 16847720-16847740 | 16847589-16847748 | (+) |
| miR535a | Chr02 | 34680467-34680487 | | 34680366-34680506 | (+) | miR535a | Chr02 | 33228837-33228857 | | 33228736-33228876 | (+) |
| | | | | | | miR535b | Chr02 | 33242827-33242847 | 33242887-33242907 | 33242797-33242937 | (+) |
| miR535c | Chr02 | 34680467-34680487 | 34680527-34680547 | 34680437-34680577 | (+) | | | | | | |
| miR827 | Chr02 | 22638433-22638453 | | 22638513-22638414 | (-) | miR827 | Chr02 | 22338642-22338662 | | 22338722-22338625 | (-) |
| miR857 | Chr02 | 3944810-3944833 | | 3944946-3944798 | (-) | | | | | | |
| miR858 | Chr02 | | 38538676-38538696 | 38538632-38538708 | (+) | miR858 | Chr02 | | 37068168-37068188 | 37068124-37068198 | (+) |
| miR9560 | Chr02 | 39957844-39957867 | 39957881-39957904 | 39957824-39957926 | (+) | miR9560 | Chr02 | 38421410-38421433 | 38421447-38421470 | 38421390-38421492 | (+) |

**Supplementary Table 2: Coordinates on the genome of the miRNA found with low confidence level. Putative loci.**

| Primary | | | | | | Alternative | | | | | |
|---|---|---|---|---|---|---|---|---|---|---|---|
| **miRNA mature** | **Chromosome** | **5p coordinates (start-end)** | **3p coordinates (start-end)** | **Hairpin coordinates** | **Strand** | **miRNA mature** | **Chromosome** | **5p coordinates (start-end)** | **3p coordinates (start-end)** | **Hairpin coordinates** | **Strand** |
| miR166a | Chr03 | 7800496-7800516 | 7800387-7800413 | 7800570-7800367 | (-) | miR166a | 000909F_alternative | 27103-27123 | 26994-27025 | 27177-26974 | (-) |
| miR3946 | Chr05 | 12227563-12227586 | | 12227541-12227690 | (+) | miR3946 | Chr09 | 26116762-26116785 | | 26116807-26116658 | (-) |
| miR171b | Chr05 | 18113428-18113449 | 18113352-18113372 | 18113484-18113329 | (-) | miR171b | Chr05 | 19697881-19697902 | 19697805-19697825 | 19697938-19697782 | (-) |
| miR156d | Chr02 | 51347121-51347140 | | 51347111-51347211 | (+) | miR156d | 001921F_alternative | 38946-38965 | | 38936-39036 | (+) |
| miR166h | Chr02 | | 33667850-33667871 | 33667756-33667881 | (+) | miR166h | Chr02 | | 32454580-32454601 | 32454486-32454611 | (+) |
| miR166i | Chr02 | | 33668066-33668086 | 33668001-33668098 | (+) | miR166i | Chr02 | | 32354688-32354708 | 32354621-32354720 | (+) |
| miR166j | Chr02 | | 33464271-33464291 | 33464172-33464304 | (+) | | | | | | |
| miR166k | Chr02 | | 33464484-33464504 | 33464407-33464521 | (+) | | | | | | |
| miR169b | Chr07 | | | | | miR169b | Chr07 | 16056386-16056405 | | 16056415-16056213 | (-) |
| | | | | | | miR169d | Chr02 | 20219254-20219273 | 20219181-20219201 | 20219303-20219151 | (-) |
| | | | | | | miR169i | Chr07 | 8757323-8757342 | | 8757352-8757197 | (-) |
| miR393c | Chr07 | 855039-855059 | 855101-855121 | 855009-855151 | (+) | miR393c | Chr07 | 868412-868432 | 868474-868494 | 868382-868524 | (+) |
| | | | | | | miR396d | 001821F_alternative | | 10057-10076 | 10190-10027 | (-) |
| miR399e | Chr02 | 44604832-44604852 | 44604782-44604802 | 44604882-44604752 | (-) | | | | | | |
| miR399f | Chr02 | 44988532-44988552 | 44988443-44988463 | 44988583-44988413 | (-) | | | | | | |
| miR2275a | Chr06 | 22091159-22091179 | 22091107-22091128 | 22091209-22091085 | (-) | miR2275a | Chr06 | 23608878-23608898 | 23608826-23608847 | 23608928-23608804 | (-) |
| miR3952b | Chr08 | 8751044-8751064 | 8751115-8751135 | 8751014-8751066..8751066-8751165 | (+) | miR3952b | Chr08 | 9960973-9960993 | 9961092-9961112 | 9960943-9961142 | (+) |
| miR536 | Chr07 | 16094953-16094974 | 16094891-16094911 | 14674467-14674324 | (-) | miR536 | Chr07 | 14674416-14674437 | 14674354-14674374 | 16095004-16094861 | (-) |
| miR12107 | Chr04 | 26089797-26089817 | 26089718-26089738 | 26089838-26089688 | (-) | miR12107 | Chr04 | 27062677-27062697 | 27062598-27062618 | 27062718-27062576 | (-) |
| miR12108 | Chr01 | 27308044-27308067 | 27308121-27308144 | 27308018-27308114..27308082..27308166 | (+) | miR12108 | Chr01 | 28716424-28716447 | 28716501-28716524 | 28716398-28716494..28716462-28716547 | (+) |

| | | | | | | | | | | |
|---|---|---|---|---|---|---|---|---|---|---|
| miR156i | Chr02 | 51369682-51369702 | 51369674-51369773 | (+) | miR156i | 004306F_alternative | 18760-18780 | | 18788-18691 | (-) |
| | | | | | miR3951b | Chr02 | 23503478-23503498 | 23503388-23503409 | 23503505-23503382 | (-) |

**Supplementary Table 3: Coordinate of the long trasposable elements (LTR) on the primary haplotype**

| Chromosome | Start | End | Length | Strand |
|---|---|---|---|---|
| Chr01 | 1199389 | 1204887 | 5499 | - |
| Chr01 | 1350932 | 1355994 | 5063 | + |
| Chr01 | 1513804 | 1518701 | 4898 | + |
| Chr01 | 1669501 | 1678826 | 9326 | + |
| Chr01 | 1811254 | 1827574 | 16321 | - |
| Chr01 | 2100426 | 2110084 | 9659 | - |
| Chr01 | 2134254 | 2143173 | 8920 | - |
| Chr01 | 2347913 | 2352777 | 4865 | + |
| Chr01 | 2508682 | 2513627 | 4946 | + |
| Chr01 | 2848212 | 2858801 | 10590 | + |
| Chr01 | 3531259 | 3535659 | 4401 | + |
| Chr01 | 4197759 | 4203115 | 5357 | + |
| Chr01 | 4206169 | 4214723 | 8555 | + |
| Chr01 | 5791536 | 5799564 | 8029 | - |
| Chr01 | 9899018 | 9904019 | 5002 | - |
| Chr01 | 10031417 | 10042246 | 10830 | - |
| Chr01 | 10348200 | 10353371 | 5172 | + |
| Chr01 | 10357857 | 10364664 | 6808 | - |
| Chr01 | 10377516 | 10387350 | 9835 | - |
| Chr01 | 10394690 | 10400146 | 5457 | + |
| Chr01 | 10499049 | 10504384 | 5336 | - |
| Chr01 | 10643585 | 10648673 | 5089 | + |
| Chr01 | 10669710 | 10680341 | 10632 | - |
| Chr01 | 10816612 | 10823304 | 6693 | - |
| Chr01 | 10853217 | 10859417 | 6201 | + |
| Chr01 | 11061123 | 11066550 | 5428 | + |
| Chr01 | 11363712 | 11368753 | 5042 | + |
| Chr01 | 11433804 | 11443900 | 10097 | + |
| Chr01 | 11510132 | 11515883 | 5752 | - |

| | | | | |
|---|---|---|---|---|
| Chr01 | 11776239 | 11782552 | 6314 | - |
| Chr01 | 11942055 | 11947564 | 5510 | + |
| Chr01 | 12196157 | 12218005 | 21849 | - |
| Chr01 | 12267039 | 12274084 | 7046 | + |
| Chr01 | 12293479 | 12297607 | 4129 | - |
| Chr01 | 13090568 | 13095702 | 5135 | + |
| Chr01 | 13553085 | 13561426 | 8342 | + |
| Chr01 | 13837471 | 13848725 | 11255 | + |
| Chr01 | 13977780 | 13984139 | 6360 | + |
| Chr01 | 14216592 | 14227258 | 10667 | - |
| Chr01 | 14425995 | 14446384 | 20390 | + |
| Chr01 | 14427420 | 14431536 | 4117 | - |
| Chr01 | 14432719 | 14442729 | 10011 | + |
| Chr01 | 14530373 | 14544969 | 14597 | + |
| Chr01 | 14655484 | 14660749 | 5266 | + |
| Chr01 | 14750969 | 14758010 | 7042 | - |
| Chr01 | 14784027 | 14794169 | 10143 | - |
| Chr01 | 15557133 | 15562487 | 5355 | + |
| Chr01 | 15594794 | 15601761 | 6968 | + |
| Chr01 | 16049262 | 16056774 | 7513 | + |
| Chr01 | 16212483 | 16217629 | 5147 | - |
| Chr01 | 16541372 | 16546576 | 5205 | + |
| Chr01 | 16719640 | 16730592 | 10953 | - |
| Chr01 | 16719816 | 16730174 | 10359 | - |
| Chr01 | 16720178 | 16730398 | 10221 | - |
| Chr01 | 16742135 | 16747445 | 5311 | + |
| Chr01 | 16838280 | 16846867 | 8588 | - |
| Chr01 | 16952483 | 16957878 | 5396 | + |
| Chr01 | 17003030 | 17008465 | 5436 | - |
| Chr01 | 17284613 | 17291302 | 6690 | + |
| Chr01 | 17526481 | 17531396 | 4916 | - |
| Chr01 | 17717605 | 17728970 | 11366 | - |
| Chr01 | 17760368 | 17764794 | 4427 | + |

| | | | | |
|---|---|---|---|---|
| Chr01 | 17765925 | 17783391 | 17467 | - |
| Chr01 | 17966742 | 17974400 | 7659 | + |
| Chr01 | 18367877 | 18372456 | 4580 | - |
| Chr01 | 18433405 | 18438797 | 5393 | - |
| Chr01 | 18433405 | 18444049 | 10645 | - |
| Chr01 | 18669911 | 18673514 | 3604 | + |
| Chr01 | 18669911 | 18675933 | 6023 | + |
| Chr01 | 18815227 | 18827631 | 12405 | - |
| Chr01 | 19371521 | 19380053 | 8533 | + |
| Chr01 | 19556656 | 19562018 | 5363 | - |
| Chr01 | 19675824 | 19682533 | 6710 | - |
| Chr01 | 19789689 | 19799417 | 9729 | + |
| Chr01 | 20002369 | 20009637 | 7269 | - |
| Chr01 | 20793667 | 20802139 | 8473 | + |
| Chr01 | 21034060 | 21043651 | 9592 | + |
| Chr01 | 21158751 | 21168271 | 9521 | + |
| Chr01 | 21188459 | 21193861 | 5403 | - |
| Chr01 | 21206737 | 21212316 | 5580 | + |
| Chr01 | 21286622 | 21296427 | 9806 | - |
| Chr01 | 21404159 | 21406036 | 1878 | + |
| Chr01 | 21760338 | 21763078 | 2741 | + |
| Chr01 | 21987532 | 21993030 | 5499 | + |
| Chr01 | 22174197 | 22179280 | 5084 | - |
| Chr01 | 22196158 | 22202788 | 6631 | - |
| Chr01 | 22215357 | 22225380 | 10024 | + |
| Chr01 | 22242726 | 22247637 | 4912 | + |
| Chr01 | 22701456 | 22708838 | 7383 | + |
| Chr01 | 23252717 | 23258297 | 5581 | - |
| Chr01 | 23781621 | 23787565 | 5945 | - |
| Chr01 | 23895099 | 23903587 | 8489 | - |
| Chr01 | 23939994 | 23946933 | 6940 | - |
| Chr01 | 24067570 | 24071710 | 4141 | - |
| Chr01 | 24189589 | 24194902 | 5314 | + |

| | | | | |
|---|---|---|---|---|
| Chr01 | 24494300 | 24499628 | 5329 | + |
| Chr01 | 24836975 | 24846834 | 9860 | - |
| Chr01 | 24925437 | 24930884 | 5448 | - |
| Chr01 | 25088648 | 25093807 | 5160 | + |
| Chr01 | 25247592 | 25253118 | 5527 | + |
| Chr01 | 25666850 | 25672025 | 5176 | + |
| Chr01 | 25678317 | 25685375 | 7059 | - |
| Chr01 | 25816085 | 25822109 | 6025 | - |
| Chr01 | 25979420 | 25989886 | 10467 | + |
| Chr01 | 26062216 | 26067153 | 4938 | + |
| Chr01 | 26141410 | 26146221 | 4812 | - |
| Chr01 | 26278668 | 26285593 | 6926 | - |
| Chr01 | 26519383 | 26524676 | 5294 | + |
| Chr01 | 26532346 | 26553513 | 21168 | + |
| Chr01 | 26604346 | 26611500 | 7155 | + |
| Chr01 | 27152854 | 27157575 | 4722 | + |
| Chr01 | 27543252 | 27544725 | 1474 | + |
| Chr01 | 27559733 | 27575065 | 15333 | - |
| Chr01 | 27652300 | 27654736 | 2437 | - |
| Chr01 | 28980279 | 28985351 | 5073 | - |
| Chr02 | 635071 | 637035 | 1965 | + |
| Chr02 | 785035 | 790052 | 5018 | + |
| Chr02 | 1107212 | 1112601 | 5390 | + |
| Chr02 | 1183057 | 1188303 | 5247 | + |
| Chr02 | 1222555 | 1228082 | 5528 | + |
| Chr02 | 1263655 | 1266868 | 3214 | - |
| Chr02 | 1301655 | 1307932 | 6278 | + |
| Chr02 | 1430915 | 1441415 | 10501 | - |
| Chr02 | 1464428 | 1469263 | 4836 | + |
| Chr02 | 1582569 | 1588842 | 6274 | + |
| Chr02 | 1763687 | 1787738 | 24052 | - |
| Chr02 | 1773978 | 1778478 | 4501 | + |
| Chr02 | 1796805 | 1801815 | 5011 | - |

| | | | | |
|---|---|---|---|---|
| Chr02 | 1813830 | 1837817 | 23988 | - |
| Chr02 | 1817623 | 1837817 | 20195 | - |
| Chr02 | 1826591 | 1831935 | 5345 | - |
| Chr02 | 1832836 | 1837817 | 4982 | - |
| Chr02 | 1850674 | 1855956 | 5283 | - |
| Chr02 | 1910852 | 1918804 | 7953 | + |
| Chr02 | 1968816 | 1974282 | 5467 | - |
| Chr02 | 2039606 | 2044623 | 5018 | + |
| Chr02 | 2057869 | 2064753 | 6885 | - |
| Chr02 | 2699733 | 2711917 | 12185 | - |
| Chr02 | 2706627 | 2711917 | 5291 | - |
| Chr02 | 2907990 | 2913288 | 5299 | + |
| Chr02 | 3392911 | 3399986 | 7076 | - |
| Chr02 | 3580280 | 3587933 | 7654 | + |
| Chr02 | 3603190 | 3608720 | 5531 | - |
| Chr02 | 3646559 | 3651980 | 5422 | + |
| Chr02 | 3958139 | 3967943 | 9805 | - |
| Chr02 | 3975404 | 3994648 | 19245 | + |
| Chr02 | 4003331 | 4008794 | 5464 | + |
| Chr02 | 4131822 | 4137121 | 5300 | + |
| Chr02 | 5239413 | 5251143 | 11731 | - |
| Chr02 | 5861276 | 5865273 | 3998 | + |
| Chr02 | 5861276 | 5880853 | 19578 | + |
| Chr02 | 5916192 | 5927988 | 11797 | + |
| Chr02 | 5931989 | 5940881 | 8893 | + |
| Chr02 | 6021632 | 6025138 | 3507 | - |
| Chr02 | 6252107 | 6257866 | 5760 | + |
| Chr02 | 6975649 | 6994340 | 18692 | + |
| Chr02 | 7197531 | 7202653 | 5123 | + |
| Chr02 | 7837384 | 7840071 | 2688 | + |
| Chr02 | 8254942 | 8260213 | 5272 | + |
| Chr02 | 8638893 | 8644379 | 5487 | - |
| Chr02 | 8651450 | 8656894 | 5445 | - |

| | | | | |
|---|---|---|---|---|
| Chr02 | 8833077 | 8839210 | 6134 | - |
| Chr02 | 8960691 | 8965923 | 5233 | + |
| Chr02 | 9017346 | 9039237 | 21892 | + |
| Chr02 | 9092213 | 9106196 | 13984 | - |
| Chr02 | 9114059 | 9119356 | 5298 | - |
| Chr02 | 9165159 | 9171624 | 6466 | + |
| Chr02 | 9607449 | 9612572 | 5124 | + |
| Chr02 | 9742135 | 9747431 | 5297 | + |
| Chr02 | 9931773 | 9937156 | 5384 | - |
| Chr02 | 10198219 | 10203266 | 5048 | + |
| Chr02 | 10491353 | 10504595 | 13243 | + |
| Chr02 | 10491518 | 10504578 | 13061 | - |
| Chr02 | 10854478 | 10857969 | 3492 | + |
| Chr02 | 10895233 | 10897158 | 1926 | - |
| Chr02 | 11057428 | 11060823 | 3396 | + |
| Chr02 | 11472532 | 11477941 | 5410 | - |
| Chr02 | 11559974 | 11564939 | 4966 | - |
| Chr02 | 11833901 | 11854779 | 20879 | - |
| Chr02 | 11839289 | 11847530 | 8242 | - |
| Chr02 | 12032014 | 12037745 | 5732 | - |
| Chr02 | 12281515 | 12292572 | 11058 | + |
| Chr02 | 12438774 | 12460575 | 21802 | - |
| Chr02 | 12452531 | 12459113 | 6583 | + |
| Chr02 | 12469876 | 12486893 | 17018 | - |
| Chr02 | 12471743 | 12491210 | 19468 | - |
| Chr02 | 12475995 | 12482499 | 6505 | - |
| Chr02 | 12504811 | 12510874 | 6064 | + |
| Chr02 | 12733733 | 12746862 | 13130 | - |
| Chr02 | 12733776 | 12746194 | 12419 | + |
| Chr02 | 12735001 | 12746919 | 11919 | - |
| Chr02 | 12848010 | 12858116 | 10107 | + |
| Chr02 | 12868936 | 12874517 | 5582 | + |
| Chr02 | 13071434 | 13076704 | 5271 | + |

| | | | | |
|---|---|---|---|---|
| Chr02 | 13136320 | 13142493 | 6174 | + |
| Chr02 | 13192091 | 13199194 | 7104 | - |
| Chr02 | 13347318 | 13356806 | 9489 | + |
| Chr02 | 13367308 | 13376382 | 9075 | - |
| Chr02 | 13479847 | 13484591 | 4745 | - |
| Chr02 | 13627463 | 13638830 | 11368 | + |
| Chr02 | 13667266 | 13672137 | 4872 | + |
| Chr02 | 13716656 | 13722040 | 5385 | + |
| Chr02 | 14045660 | 14055612 | 9953 | + |
| Chr02 | 14575469 | 14580432 | 4964 | - |
| Chr02 | 14824440 | 14828150 | 3711 | + |
| Chr02 | 15023567 | 15029950 | 6384 | + |
| Chr02 | 15036979 | 15054144 | 17166 | + |
| Chr02 | 15187123 | 15195418 | 8296 | - |
| Chr02 | 15827200 | 15839514 | 12315 | + |
| Chr02 | 15845562 | 15847915 | 2354 | + |
| Chr02 | 16645168 | 16653063 | 7896 | - |
| Chr02 | 16672759 | 16696371 | 23613 | - |
| Chr02 | 16740779 | 16747009 | 6231 | + |
| Chr02 | 16800939 | 16808917 | 7979 | + |
| Chr02 | 16802024 | 16808917 | 6894 | - |
| Chr02 | 16973865 | 16979284 | 5420 | + |
| Chr02 | 17103824 | 17111173 | 7350 | + |
| Chr02 | 17215770 | 17225468 | 9699 | - |
| Chr02 | 17340451 | 17345870 | 5420 | + |
| Chr02 | 17684924 | 17693261 | 8338 | + |
| Chr02 | 17968492 | 17990052 | 21561 | - |
| Chr02 | 18274323 | 18285145 | 10823 | + |
| Chr02 | 18464576 | 18479248 | 14673 | + |
| Chr02 | 18483871 | 18487397 | 3527 | + |
| Chr02 | 18489038 | 18504105 | 15068 | - |
| Chr02 | 18623354 | 18630580 | 7227 | + |
| Chr02 | 18806434 | 18811189 | 4756 | + |

| | | | | |
|---|---|---|---|---|
| Chr02 | 18991770 | 18996662 | 4893 | - |
| Chr02 | 19729305 | 19740066 | 10762 | - |
| Chr02 | 20116311 | 20123228 | 6918 | + |
| Chr02 | 20180054 | 20189921 | 9868 | + |
| Chr02 | 20315485 | 20325350 | 9866 | + |
| Chr02 | 20618346 | 20624976 | 6631 | + |
| Chr02 | 20747631 | 20756571 | 8941 | + |
| Chr02 | 20828324 | 20835291 | 6968 | - |
| Chr02 | 20872857 | 20875389 | 2533 | - |
| Chr02 | 20891998 | 20901152 | 9155 | + |
| Chr02 | 20891998 | 20901627 | 9630 | + |
| Chr02 | 21212415 | 21222230 | 9816 | - |
| Chr02 | 21272369 | 21278358 | 5990 | + |
| Chr02 | 21450402 | 21469972 | 19571 | - |
| Chr02 | 21458293 | 21469090 | 10798 | - |
| Chr02 | 21551334 | 21555878 | 4545 | + |
| Chr02 | 21841280 | 21846206 | 4927 | + |
| Chr02 | 22152352 | 22166404 | 14053 | - |
| Chr02 | 22216832 | 22222269 | 5438 | - |
| Chr02 | 22362110 | 22372672 | 10563 | - |
| Chr02 | 22389501 | 22400089 | 10589 | - |
| Chr02 | 22709914 | 22715580 | 5667 | + |
| Chr02 | 22983565 | 22988597 | 5033 | + |
| Chr02 | 23352550 | 23361426 | 8877 | - |
| Chr02 | 23369891 | 23375440 | 5550 | - |
| Chr02 | 23425890 | 23431407 | 5518 | - |
| Chr02 | 23449633 | 23465922 | 16290 | + |
| Chr02 | 23631122 | 23641894 | 10773 | + |
| Chr02 | 23749244 | 23761816 | 12573 | - |
| Chr02 | 23784638 | 23788766 | 4129 | - |
| Chr02 | 23809419 | 23812135 | 2717 | + |
| Chr02 | 23826470 | 23831564 | 5095 | + |
| Chr02 | 23871192 | 23880249 | 9058 | - |

| | | | | |
|---|---|---|---|---|
| Chr02 | 24093393 | 24098843 | 5451 | + |
| Chr02 | 24178102 | 24183456 | 5355 | - |
| Chr02 | 24251980 | 24257572 | 5593 | - |
| Chr02 | 24261335 | 24266648 | 5314 | + |
| Chr02 | 24298041 | 24308130 | 10090 | - |
| Chr02 | 24302919 | 24308146 | 5228 | - |
| Chr02 | 24364555 | 24372765 | 8211 | + |
| Chr02 | 24518009 | 24530609 | 12601 | - |
| Chr02 | 24661653 | 24667214 | 5562 | + |
| Chr02 | 24716453 | 24721574 | 5122 | + |
| Chr02 | 24910420 | 24932290 | 21871 | + |
| Chr02 | 25050192 | 25055358 | 5167 | + |
| Chr02 | 25261754 | 25269952 | 8199 | + |
| Chr02 | 25477314 | 25486680 | 9367 | - |
| Chr02 | 25660948 | 25688379 | 27432 | - |
| Chr02 | 25664676 | 25693635 | 28960 | - |
| Chr02 | 25667233 | 25684929 | 17697 | - |
| Chr02 | 25667458 | 25693635 | 26178 | - |
| Chr02 | 25671019 | 25687699 | 16681 | - |
| Chr02 | 25867492 | 25878452 | 10961 | + |
| Chr02 | 25992138 | 25997551 | 5414 | - |
| Chr02 | 26009966 | 26018210 | 8245 | + |
| Chr02 | 26084926 | 26090336 | 5411 | + |
| Chr02 | 26103390 | 26113532 | 10143 | - |
| Chr02 | 26103390 | 26113950 | 10561 | - |
| Chr02 | 26240613 | 26242505 | 1893 | - |
| Chr02 | 26304162 | 26316217 | 12056 | + |
| Chr02 | 26468625 | 26476960 | 8336 | + |
| Chr02 | 27000251 | 27002682 | 2432 | + |
| Chr02 | 27381907 | 27396435 | 14529 | + |
| Chr02 | 27401128 | 27403135 | 2008 | + |
| Chr02 | 27791644 | 27802631 | 10988 | - |
| Chr02 | 27825563 | 27832782 | 7220 | + |

| | | | | |
|---|---|---|---|---|
| Chr02 | 28036407 | 28057771 | 21365 | - |
| Chr02 | 28057784 | 28075209 | 17426 | - |
| Chr02 | 28549955 | 28554952 | 4998 | + |
| Chr02 | 28602638 | 28618729 | 16092 | + |
| Chr02 | 28692620 | 28697665 | 5046 | + |
| Chr02 | 28732305 | 28751053 | 18749 | + |
| Chr02 | 28737918 | 28743080 | 5163 | - |
| Chr02 | 28757304 | 28762645 | 5342 | + |
| Chr02 | 29345671 | 29358927 | 13257 | + |
| Chr02 | 30128976 | 30148452 | 19477 | + |
| Chr02 | 30129382 | 30139888 | 10507 | + |
| Chr02 | 30531241 | 30538063 | 6823 | + |
| Chr02 | 30581953 | 30588308 | 6356 | - |
| Chr02 | 30581953 | 30588752 | 6800 | - |
| Chr02 | 30582360 | 30588749 | 6390 | - |
| Chr02 | 30617279 | 30619269 | 1991 | - |
| Chr02 | 30624622 | 30633157 | 8536 | - |
| Chr02 | 30682837 | 30694529 | 11693 | - |
| Chr02 | 30770395 | 30782639 | 12245 | + |
| Chr02 | 30788783 | 30801393 | 12611 | - |
| Chr02 | 31327811 | 31333976 | 6166 | - |
| Chr02 | 31494979 | 31502900 | 7922 | + |
| Chr02 | 31584900 | 31586859 | 1960 | - |
| Chr02 | 31749797 | 31769756 | 19960 | + |
| Chr02 | 31908219 | 31915153 | 6935 | - |
| Chr02 | 31920263 | 31926103 | 5841 | + |
| Chr02 | 31932810 | 31934834 | 2025 | + |
| Chr02 | 32000764 | 32007614 | 6851 | + |
| Chr02 | 32177150 | 32190193 | 13044 | - |
| Chr02 | 32181734 | 32189239 | 7506 | - |
| Chr02 | 32265191 | 32267213 | 2023 | + |
| Chr02 | 32279042 | 32284501 | 5460 | - |
| Chr02 | 32372813 | 32378527 | 5715 | + |

| | | | | |
|---|---|---|---|---|
| Chr02 | 32761715 | 32766904 | 5190 | - |
| Chr02 | 32778149 | 32783393 | 5245 | - |
| Chr02 | 32893263 | 32895478 | 2216 | + |
| Chr02 | 33018528 | 33020508 | 1981 | + |
| Chr02 | 33337487 | 33342478 | 4992 | + |
| Chr02 | 33519859 | 33524966 | 5108 | - |
| Chr02 | 33673128 | 33690422 | 17295 | + |
| Chr02 | 33674452 | 33690422 | 15971 | + |
| Chr02 | 33766018 | 33772091 | 6074 | + |
| Chr02 | 33843868 | 33854176 | 10309 | + |
| Chr02 | 33848411 | 33853692 | 5282 | + |
| Chr02 | 33977833 | 33980273 | 2441 | - |
| Chr02 | 34086936 | 34092277 | 5342 | + |
| Chr02 | 34211434 | 34228303 | 16870 | + |
| Chr02 | 34253240 | 34260347 | 7108 | - |
| Chr02 | 34455554 | 34460694 | 5141 | + |
| Chr02 | 34525475 | 34544433 | 18959 | - |
| Chr02 | 34530965 | 34537297 | 6333 | + |
| Chr02 | 34633514 | 34639936 | 6423 | + |
| Chr02 | 34748249 | 34760505 | 12257 | + |
| Chr02 | 34753179 | 34758817 | 5639 | + |
| Chr02 | 34771529 | 34776894 | 5366 | + |
| Chr02 | 34816865 | 34827160 | 10296 | + |
| Chr02 | 34914424 | 34921904 | 7481 | - |
| Chr02 | 34967146 | 34977706 | 10561 | + |
| Chr02 | 35103471 | 35115611 | 12141 | + |
| Chr02 | 35134546 | 35137497 | 2952 | + |
| Chr02 | 35138477 | 35150627 | 12151 | + |
| Chr02 | 35260526 | 35270066 | 9541 | + |
| Chr02 | 35284239 | 35290679 | 6441 | + |
| Chr02 | 35321742 | 35327338 | 5597 | - |
| Chr02 | 35406642 | 35412859 | 6218 | - |
| Chr02 | 35503287 | 35512997 | 9711 | + |

| | | | | |
|---|---|---|---|---|
| Chr02 | 35845841 | 35850743 | 4903 | - |
| Chr02 | 36137963 | 36148544 | 10582 | + |
| Chr02 | 36301800 | 36307414 | 5615 | + |
| Chr02 | 36672122 | 36678482 | 6361 | + |
| Chr02 | 36913632 | 36923356 | 9725 | + |
| Chr02 | 36931206 | 36936747 | 5542 | + |
| Chr02 | 36937679 | 36953095 | 15417 | + |
| Chr02 | 36988271 | 37000458 | 12188 | + |
| Chr02 | 37016467 | 37042961 | 26495 | + |
| Chr02 | 37051917 | 37057465 | 5549 | + |
| Chr02 | 37202038 | 37224454 | 22417 | + |
| Chr02 | 37278182 | 37283728 | 5547 | + |
| Chr02 | 37576302 | 37581609 | 5308 | + |
| Chr02 | 37582311 | 37592081 | 9771 | - |
| Chr02 | 38043317 | 38052209 | 8893 | - |
| Chr02 | 38105216 | 38110122 | 4907 | + |
| Chr02 | 38431817 | 38436981 | 5165 | - |
| Chr02 | 39115047 | 39121377 | 6331 | + |
| Chr02 | 39391397 | 39400500 | 9104 | + |
| Chr02 | 40017046 | 40019265 | 2220 | - |
| Chr02 | 40605098 | 40611409 | 6312 | - |
| Chr02 | 40752498 | 40757815 | 5318 | - |
| Chr02 | 40851667 | 40856906 | 5240 | + |
| Chr02 | 41129322 | 41150405 | 21084 | + |
| Chr02 | 41158742 | 41163599 | 4858 | - |
| Chr02 | 41244694 | 41251363 | 6670 | + |
| Chr02 | 41311180 | 41316607 | 5428 | + |
| Chr02 | 41373144 | 41380757 | 7614 | + |
| Chr02 | 41481444 | 41499067 | 17624 | + |
| Chr02 | 41701856 | 41712160 | 10305 | - |
| Chr02 | 41822129 | 41827366 | 5238 | - |
| Chr02 | 41930664 | 41935857 | 5194 | + |
| Chr02 | 41966560 | 41976606 | 10047 | - |

| | | | | |
|---|---|---|---|---|
| Chr02 | 42148989 | 42157157 | 8169 | + |
| Chr02 | 42938387 | 42943341 | 4955 | + |
| Chr02 | 43962867 | 43969794 | 6928 | + |
| Chr02 | 44589566 | 44595158 | 5593 | - |
| Chr02 | 44670559 | 44677658 | 7100 | - |
| Chr02 | 46851184 | 46873238 | 22055 | + |
| Chr02 | 46852437 | 46875227 | 22791 | + |
| Chr02 | 46873253 | 46874912 | 1660 | - |
| Chr02 | 46875231 | 46880118 | 4888 | - |
| Chr02 | 47343508 | 47350557 | 7050 | + |
| Chr02 | 47361443 | 47369172 | 7730 | + |
| Chr02 | 47389621 | 47395896 | 6276 | - |
| Chr02 | 47958927 | 47960492 | 1566 | + |
| Chr02 | 48630471 | 48657813 | 27343 | - |
| Chr02 | 48645506 | 48656230 | 10725 | - |
| Chr02 | 48651010 | 48656230 | 5221 | - |
| Chr02 | 48661670 | 48666773 | 5104 | + |
| Chr02 | 49458521 | 49463573 | 5053 | + |
| Chr02 | 49476111 | 49479973 | 3863 | - |
| Chr02 | 49543953 | 49547808 | 3856 | - |
| Chr02 | 49705779 | 49712067 | 6289 | + |
| Chr02 | 49764594 | 49770006 | 5413 | - |
| Chr02 | 49798739 | 49801936 | 3198 | - |
| Chr02 | 50301722 | 50306971 | 5250 | + |
| Chr02 | 50308080 | 50317747 | 9668 | + |
| Chr02 | 50989485 | 50994766 | 5282 | + |
| Chr02 | 51331087 | 51336543 | 5457 | - |
| Chr02 | 51378591 | 51386006 | 7416 | - |
| Chr03 | 232373 | 242331 | 9959 | - |
| Chr03 | 324799 | 329081 | 4283 | - |
| Chr03 | 385707 | 387675 | 1969 | - |
| Chr03 | 464860 | 469699 | 4840 | + |
| Chr03 | 558405 | 565976 | 7572 | + |

| | | | | |
|---|---|---|---|---|
| Chr03 | 679259 | 682558 | 3300 | + |
| Chr03 | 885628 | 900780 | 15153 | + |
| Chr03 | 940983 | 963389 | 22407 | + |
| Chr03 | 989247 | 1000220 | 10974 | - |
| Chr03 | 1029282 | 1040020 | 10739 | + |
| Chr03 | 1491507 | 1507855 | 16349 | - |
| Chr03 | 1562604 | 1569775 | 7172 | + |
| Chr03 | 1623383 | 1629465 | 6083 | - |
| Chr03 | 1701651 | 1707007 | 5357 | - |
| Chr03 | 1770083 | 1782365 | 12283 | + |
| Chr03 | 2261002 | 2269098 | 8097 | + |
| Chr03 | 2383336 | 2388943 | 5608 | + |
| Chr03 | 2415873 | 2442457 | 26585 | + |
| Chr03 | 2420572 | 2443851 | 23280 | + |
| Chr03 | 2432399 | 2437695 | 5297 | + |
| Chr03 | 2451616 | 2473472 | 21857 | - |
| Chr03 | 2604695 | 2611339 | 6645 | - |
| Chr03 | 2812734 | 2817921 | 5188 | + |
| Chr03 | 2988627 | 2995933 | 7307 | + |
| Chr03 | 3155290 | 3160529 | 5240 | - |
| Chr03 | 3434910 | 3441179 | 6270 | + |
| Chr03 | 3512686 | 3518073 | 5388 | - |
| Chr03 | 3584613 | 3589976 | 5364 | + |
| Chr03 | 3701110 | 3706175 | 5066 | + |
| Chr03 | 3718431 | 3723744 | 5314 | + |
| Chr03 | 3879622 | 3885695 | 6074 | + |
| Chr03 | 3923310 | 3930427 | 7118 | - |
| Chr03 | 3970056 | 3982399 | 12344 | + |
| Chr03 | 4002129 | 4008460 | 6332 | + |
| Chr03 | 4123598 | 4141951 | 18354 | - |
| Chr03 | 4125984 | 4141951 | 15968 | - |
| Chr03 | 4481698 | 4490168 | 8471 | + |
| Chr03 | 4674221 | 4684135 | 9915 | + |

| | | | | |
|---|---|---|---|---|
| Chr03 | 4761039 | 4767519 | 6481 | - |
| Chr03 | 5042797 | 5047791 | 4995 | + |
| Chr03 | 5048631 | 5053950 | 5320 | - |
| Chr03 | 5091746 | 5097352 | 5607 | - |
| Chr03 | 5235670 | 5244309 | 8640 | - |
| Chr03 | 5286470 | 5293069 | 6600 | + |
| Chr03 | 5294191 | 5300866 | 6676 | + |
| Chr03 | 5402688 | 5410175 | 7488 | + |
| Chr03 | 5526542 | 5534010 | 7469 | + |
| Chr03 | 5647249 | 5651574 | 4326 | + |
| Chr03 | 5936009 | 5942228 | 6220 | - |
| Chr03 | 6210460 | 6215904 | 5445 | - |
| Chr03 | 6394025 | 6406711 | 12687 | - |
| Chr03 | 6394051 | 6406308 | 12258 | + |
| Chr03 | 6445881 | 6451261 | 5381 | + |
| Chr03 | 6548749 | 6555436 | 6688 | - |
| Chr03 | 6763607 | 6768542 | 4936 | - |
| Chr03 | 6902779 | 6908113 | 5335 | - |
| Chr03 | 7058645 | 7062773 | 4129 | + |
| Chr03 | 7129354 | 7139170 | 9817 | + |
| Chr03 | 7296158 | 7301313 | 5156 | - |
| Chr03 | 7367026 | 7372081 | 5056 | + |
| Chr03 | 7439156 | 7441219 | 2064 | + |
| Chr03 | 7573944 | 7577296 | 3353 | + |
| Chr03 | 7632063 | 7639870 | 7808 | + |
| Chr03 | 7736859 | 7742405 | 5547 | + |
| Chr03 | 7967686 | 7972301 | 4616 | + |
| Chr03 | 8012412 | 8021365 | 8954 | + |
| Chr03 | 8037984 | 8043156 | 5173 | + |
| Chr03 | 8126733 | 8132243 | 5511 | + |
| Chr03 | 8294944 | 8300330 | 5387 | + |
| Chr03 | 8307437 | 8313044 | 5608 | - |
| Chr03 | 8473167 | 8477105 | 3939 | + |

| | | | | |
|---|---|---|---|---|
| Chr03 | 8808885 | 8814082 | 5198 | - |
| Chr03 | 8808885 | 8814082 | 5198 | - |
| Chr03 | 8832012 | 8835663 | 3652 | - |
| Chr03 | 9339573 | 9345060 | 5488 | - |
| Chr03 | 9681182 | 9685976 | 4795 | - |
| Chr03 | 9877732 | 9890511 | 12780 | - |
| Chr03 | 10057110 | 10065170 | 8061 | + |
| Chr03 | 10076188 | 10088843 | 12656 | - |
| Chr03 | 10860698 | 10873860 | 13163 | + |
| Chr03 | 10938762 | 10943717 | 4956 | + |
| Chr03 | 11455514 | 11460806 | 5293 | - |
| Chr03 | 11638391 | 11644004 | 5614 | + |
| Chr03 | 11656750 | 11666824 | 10075 | - |
| Chr03 | 11663391 | 11666391 | 3001 | - |
| Chr03 | 11669057 | 11674508 | 5452 | + |
| Chr03 | 11970836 | 11977429 | 6594 | + |
| Chr03 | 12546132 | 12551173 | 5042 | + |
| Chr03 | 12642168 | 12647593 | 5426 | - |
| Chr03 | 12708832 | 12714256 | 5425 | + |
| Chr03 | 12735194 | 12740179 | 4986 | + |
| Chr03 | 12895479 | 12906399 | 10921 | - |
| Chr03 | 12950285 | 12955745 | 5461 | - |
| Chr03 | 13383726 | 13392566 | 8841 | + |
| Chr03 | 13502508 | 13507452 | 4945 | + |
| Chr03 | 13848238 | 13853852 | 5615 | - |
| Chr03 | 13960312 | 13965174 | 4863 | + |
| Chr03 | 14149700 | 14155160 | 5461 | - |
| Chr03 | 14360593 | 14365519 | 4927 | + |
| Chr03 | 14415683 | 14420707 | 5025 | + |
| Chr03 | 14518212 | 14539546 | 21335 | - |
| Chr03 | 14829481 | 14835121 | 5641 | - |
| Chr03 | 15104892 | 15110069 | 5178 | + |
| Chr03 | 15539562 | 15548897 | 9336 | - |

| | | | | |
|---|---|---|---|---|
| Chr03 | 15674173 | 15680609 | 6437 | + |
| Chr03 | 15713374 | 15720328 | 6955 | + |
| Chr03 | 16007457 | 16022474 | 15018 | - |
| Chr03 | 16031080 | 16043293 | 12214 | + |
| Chr03 | 16047128 | 16054713 | 7586 | + |
| Chr03 | 16129960 | 16149246 | 19287 | + |
| Chr03 | 16323299 | 16337613 | 14315 | - |
| Chr03 | 16363161 | 16368477 | 5317 | - |
| Chr03 | 16373613 | 16382905 | 9293 | - |
| Chr03 | 16400811 | 16406141 | 5331 | - |
| Chr03 | 16429676 | 16436870 | 7195 | + |
| Chr03 | 16823295 | 16828372 | 5078 | + |
| Chr03 | 16857843 | 16863218 | 5376 | + |
| Chr03 | 16956369 | 16961102 | 4734 | - |
| Chr03 | 17151664 | 17161385 | 9722 | - |
| Chr03 | 17193023 | 17198446 | 5424 | - |
| Chr03 | 17286432 | 17291742 | 5311 | - |
| Chr03 | 17320186 | 17323234 | 3049 | - |
| Chr03 | 17376797 | 17379850 | 3054 | - |
| Chr03 | 17717793 | 17724502 | 6710 | + |
| Chr03 | 18146421 | 18156770 | 10350 | + |
| Chr03 | 18732142 | 18738277 | 6136 | - |
| Chr03 | 19637750 | 19642786 | 5037 | + |
| Chr03 | 20833925 | 20837487 | 3563 | + |
| Chr03 | 20836001 | 20839615 | 3615 | + |
| Chr03 | 20836529 | 20840844 | 4316 | + |
| Chr03 | 20837585 | 20839615 | 2031 | + |
| Chr03 | 21005664 | 21011075 | 5412 | + |
| Chr03 | 21789382 | 21794817 | 5436 | + |
| Chr03 | 22959926 | 22962366 | 2441 | - |
| Chr03 | 24438105 | 24452017 | 13913 | + |
| Chr03 | 25175871 | 25184340 | 8470 | + |
| Chr03 | 25757735 | 25763384 | 5650 | + |

| | | | | |
|---|---|---|---|---|
| Chr03 | 26064510 | 26075593 | 11084 | + |
| Chr03 | 26554262 | 26559507 | 5246 | - |
| Chr04 | 400303 | 410991 | 10689 | + |
| Chr04 | 842202 | 854364 | 12163 | + |
| Chr04 | 844090 | 850936 | 6847 | - |
| Chr04 | 2237229 | 2243598 | 6370 | + |
| Chr04 | 2245902 | 2252093 | 6192 | + |
| Chr04 | 2468892 | 2473644 | 4753 | + |
| Chr04 | 2515815 | 2520779 | 4965 | - |
| Chr04 | 2536938 | 2541973 | 5036 | + |
| Chr04 | 3322673 | 3327833 | 5161 | + |
| Chr04 | 4034630 | 4040071 | 5442 | - |
| Chr04 | 4058165 | 4063592 | 5428 | - |
| Chr04 | 4121871 | 4127367 | 5497 | - |
| Chr04 | 4205806 | 4211102 | 5297 | - |
| Chr04 | 4277663 | 4292176 | 14514 | + |
| Chr04 | 4897352 | 4903996 | 6645 | + |
| Chr04 | 4999248 | 5004782 | 5535 | + |
| Chr04 | 5335679 | 5340501 | 4823 | - |
| Chr04 | 6295518 | 6303294 | 7777 | + |
| Chr04 | 6324041 | 6332820 | 8780 | + |
| Chr04 | 6341577 | 6347118 | 5542 | + |
| Chr04 | 6556700 | 6565424 | 8725 | + |
| Chr04 | 6663228 | 6668638 | 5411 | - |
| Chr04 | 6675327 | 6680822 | 5496 | + |
| Chr04 | 6781228 | 6786619 | 5392 | - |
| Chr04 | 6935892 | 6943341 | 7450 | - |
| Chr04 | 7042864 | 7048103 | 5240 | + |
| Chr04 | 7164496 | 7175347 | 10852 | + |
| Chr04 | 7242365 | 7247556 | 5192 | - |
| Chr04 | 7305385 | 7310528 | 5144 | - |
| Chr04 | 7433671 | 7439211 | 5541 | - |
| Chr04 | 7536917 | 7541518 | 4602 | + |

| | | | | |
|---|---|---|---|---|
| Chr04 | 7551765 | 7571996 | 20232 | - |
| Chr04 | 7552119 | 7562332 | 10214 | - |
| Chr04 | 7556458 | 7562094 | 5637 | - |
| Chr04 | 8022167 | 8033204 | 11038 | - |
| Chr04 | 8112776 | 8116726 | 3951 | + |
| Chr04 | 8147373 | 8157057 | 9685 | - |
| Chr04 | 8189867 | 8209420 | 19554 | + |
| Chr04 | 8338937 | 8344111 | 5175 | - |
| Chr04 | 8356810 | 8361983 | 5174 | + |
| Chr04 | 8356810 | 8378840 | 22031 | + |
| Chr04 | 8373636 | 8378840 | 5205 | + |
| Chr04 | 8373636 | 8395318 | 21683 | + |
| Chr04 | 8390107 | 8395318 | 5212 | + |
| Chr04 | 8453650 | 8460203 | 6554 | - |
| Chr04 | 8586527 | 8591727 | 5201 | + |
| Chr04 | 8620985 | 8626437 | 5453 | + |
| Chr04 | 8778975 | 8789922 | 10948 | - |
| Chr04 | 8833168 | 8838096 | 4929 | + |
| Chr04 | 8863112 | 8872854 | 9743 | - |
| Chr04 | 8940609 | 8947385 | 6777 | - |
| Chr04 | 9039546 | 9044631 | 5086 | - |
| Chr04 | 9051767 | 9071547 | 19781 | - |
| Chr04 | 9078325 | 9095107 | 16783 | + |
| Chr04 | 9081607 | 9092130 | 10524 | + |
| Chr04 | 9160297 | 9165597 | 5301 | - |
| Chr04 | 9252313 | 9261767 | 9455 | + |
| Chr04 | 9418860 | 9423568 | 4709 | + |
| Chr04 | 9520753 | 9528258 | 7506 | + |
| Chr04 | 9582853 | 9588031 | 5179 | - |
| Chr04 | 9773949 | 9791381 | 17433 | - |
| Chr04 | 9969573 | 9983082 | 13510 | - |
| Chr04 | 10064320 | 10071683 | 7364 | - |
| Chr04 | 10072072 | 10075451 | 3380 | + |

| | | | | |
|---|---|---|---|---|
| Chr04 | 10072093 | 10074452 | 2360 | + |
| Chr04 | 10072093 | 10074729 | 2637 | + |
| Chr04 | 10072093 | 10075005 | 2913 | + |
| Chr04 | 10072104 | 10074185 | 2082 | + |
| Chr04 | 10072104 | 10075289 | 3186 | + |
| Chr04 | 10477909 | 10482808 | 4900 | + |
| Chr04 | 10512205 | 10517267 | 5063 | - |
| Chr04 | 10953805 | 10959002 | 5198 | + |
| Chr04 | 10973332 | 10978738 | 5407 | - |
| Chr04 | 11086371 | 11091679 | 5309 | - |
| Chr04 | 11093478 | 11098461 | 4984 | + |
| Chr04 | 11669923 | 11680750 | 10828 | - |
| Chr04 | 11814178 | 11823765 | 9588 | - |
| Chr04 | 11861530 | 11872152 | 10623 | - |
| Chr04 | 11897067 | 11903132 | 6066 | - |
| Chr04 | 12093590 | 12102301 | 8712 | - |
| Chr04 | 13153054 | 13162806 | 9753 | - |
| Chr04 | 13192892 | 13213073 | 20182 | + |
| Chr04 | 13510059 | 13515456 | 5398 | + |
| Chr04 | 13821392 | 13826135 | 4744 | + |
| Chr04 | 14071352 | 14076859 | 5508 | + |
| Chr04 | 14228364 | 14233839 | 5476 | + |
| Chr04 | 14228364 | 14233839 | 5476 | + |
| Chr04 | 14625559 | 14632512 | 6954 | - |
| Chr04 | 14678552 | 14683977 | 5426 | - |
| Chr04 | 14696216 | 14703049 | 6834 | + |
| Chr04 | 14988439 | 14998112 | 9674 | + |
| Chr04 | 15024765 | 15029112 | 4348 | + |
| Chr04 | 15050101 | 15055110 | 5010 | + |
| Chr04 | 15207902 | 15217870 | 9969 | + |
| Chr04 | 15483934 | 15490731 | 6798 | + |
| Chr04 | 15716694 | 15722353 | 5660 | - |
| Chr04 | 15961805 | 15967212 | 5408 | + |

| | | | | |
|---|---|---|---|---|
| Chr04 | 16022602 | 16027548 | 4947 | + |
| Chr04 | 16023160 | 16027429 | 4270 | - |
| Chr04 | 16067252 | 16080515 | 13264 | - |
| Chr04 | 16748813 | 16760964 | 12152 | + |
| Chr04 | 16835079 | 16843870 | 8792 | - |
| Chr04 | 17100936 | 17107935 | 7000 | + |
| Chr04 | 17409248 | 17412855 | 3608 | - |
| Chr04 | 17470558 | 17480651 | 10094 | + |
| Chr04 | 17542091 | 17544598 | 2508 | - |
| Chr04 | 17648763 | 17654847 | 6085 | - |
| Chr04 | 17826651 | 17834194 | 7544 | + |
| Chr04 | 18122457 | 18131174 | 8718 | + |
| Chr04 | 18302488 | 18306541 | 4054 | + |
| Chr04 | 18502834 | 18509468 | 6635 | + |
| Chr04 | 18739622 | 18744873 | 5252 | + |
| Chr04 | 19118115 | 19120438 | 2324 | + |
| Chr04 | 19336721 | 19358221 | 21501 | + |
| Chr04 | 19337504 | 19352475 | 14972 | + |
| Chr04 | 19466994 | 19482724 | 15731 | - |
| Chr04 | 19468405 | 19482724 | 14320 | - |
| Chr04 | 19643175 | 19652649 | 9475 | + |
| Chr04 | 19657352 | 19665484 | 8133 | + |
| Chr04 | 19703182 | 19708455 | 5274 | + |
| Chr04 | 19717033 | 19722760 | 5728 | + |
| Chr04 | 19735882 | 19749659 | 13778 | - |
| Chr04 | 19884318 | 19888283 | 3966 | - |
| Chr04 | 19902990 | 19908318 | 5329 | + |
| Chr04 | 20070067 | 20077830 | 7764 | + |
| Chr04 | 20117619 | 20123135 | 5517 | + |
| Chr04 | 20553601 | 20558778 | 5178 | + |
| Chr04 | 20803914 | 20820346 | 16433 | + |
| Chr04 | 21134766 | 21139458 | 4693 | + |
| Chr04 | 23299701 | 23307620 | 7920 | + |

| | | | | |
|---|---|---|---|---|
| Chr04 | 23373237 | 23378534 | 5298 | - |
| Chr04 | 23448327 | 23457840 | 9514 | - |
| Chr04 | 23469447 | 23472453 | 3007 | - |
| Chr04 | 23895288 | 23900669 | 5382 | + |
| Chr04 | 23989674 | 23995220 | 5547 | + |
| Chr04 | 24398255 | 24403399 | 5145 | + |
| Chr04 | 24421216 | 24424360 | 3145 | - |
| Chr04 | 25502170 | 25507419 | 5250 | + |
| Chr04 | 26401279 | 26411111 | 9833 | - |
| Chr04 | 26623873 | 26634254 | 10382 | + |
| Chr04 | 26766004 | 26770095 | 4092 | - |
| Chr05 | 61546 | 68173 | 6628 | - |
| Chr05 | 144017 | 148965 | 4949 | + |
| Chr05 | 371319 | 377747 | 6429 | + |
| Chr05 | 1326170 | 1336341 | 10172 | + |
| Chr05 | 1987957 | 1993596 | 5640 | - |
| Chr05 | 4314153 | 4326086 | 11934 | + |
| Chr05 | 5196877 | 5220050 | 23174 | - |
| Chr05 | 5236399 | 5266738 | 30340 | - |
| Chr05 | 5666628 | 5685241 | 18614 | - |
| Chr05 | 6403372 | 6414120 | 10749 | - |
| Chr05 | 6753682 | 6758476 | 4795 | - |
| Chr05 | 7254158 | 7258898 | 4741 | - |
| Chr05 | 8585346 | 8587430 | 2085 | + |
| Chr05 | 8642375 | 8647800 | 5426 | - |
| Chr05 | 8663040 | 8668639 | 5600 | - |
| Chr05 | 8783333 | 8789054 | 5722 | + |
| Chr05 | 8813714 | 8818824 | 5111 | + |
| Chr05 | 9780823 | 9786039 | 5217 | + |
| Chr05 | 9890034 | 9894086 | 4053 | - |
| Chr05 | 9927076 | 9935590 | 8515 | + |
| Chr05 | 9970867 | 9975969 | 5103 | - |
| Chr05 | 9995696 | 10001824 | 6129 | + |

| | | | | |
|---|---|---|---|---|
| Chr05 | 10015047 | 10023859 | 8813 | + |
| Chr05 | 10439002 | 10454081 | 15080 | - |
| Chr05 | 10444483 | 10452634 | 8152 | + |
| Chr05 | 10534309 | 10543485 | 9177 | + |
| Chr05 | 10976658 | 10990254 | 13597 | - |
| Chr05 | 11080588 | 11092756 | 12169 | - |
| Chr05 | 11182926 | 11188025 | 5100 | + |
| Chr05 | 11274650 | 11279776 | 5127 | + |
| Chr05 | 11298200 | 11303121 | 4922 | + |
| Chr05 | 11332131 | 11345247 | 13117 | - |
| Chr05 | 11494378 | 11499868 | 5491 | + |
| Chr05 | 11507984 | 11517664 | 9681 | + |
| Chr05 | 11523850 | 11529013 | 5164 | + |
| Chr05 | 11553170 | 11560410 | 7241 | + |
| Chr05 | 11646990 | 11654357 | 7368 | - |
| Chr05 | 11675351 | 11680357 | 5007 | - |
| Chr05 | 12169967 | 12178318 | 8352 | + |
| Chr05 | 12344730 | 12361012 | 16283 | - |
| Chr05 | 12353135 | 12359275 | 6141 | + |
| Chr05 | 12366678 | 12375230 | 8553 | + |
| Chr05 | 12536452 | 12546334 | 9883 | + |
| Chr05 | 12684747 | 12690139 | 5393 | - |
| Chr05 | 12755738 | 12765928 | 10191 | - |
| Chr05 | 13011805 | 13018795 | 6991 | - |
| Chr05 | 13052387 | 13063021 | 10635 | + |
| Chr05 | 13052387 | 13065431 | 13045 | + |
| Chr05 | 13052387 | 13069053 | 16667 | + |
| Chr05 | 13080762 | 13090470 | 9709 | - |
| Chr05 | 13399900 | 13405335 | 5436 | - |
| Chr05 | 13584286 | 13594029 | 9744 | - |
| Chr05 | 13670396 | 13676892 | 6497 | + |
| Chr05 | 13712336 | 13713740 | 1405 | - |
| Chr05 | 14110793 | 14120650 | 9858 | - |

| | | | | |
|---|---|---|---|---|
| Chr05 | 14217259 | 14236219 | 18961 | + |
| Chr05 | 14361897 | 14367224 | 5328 | - |
| Chr05 | 15123794 | 15130398 | 6605 | + |
| Chr05 | 15296745 | 15314993 | 18249 | + |
| Chr05 | 15469996 | 15477707 | 7712 | - |
| Chr05 | 16457338 | 16462758 | 5421 | + |
| Chr05 | 16540706 | 16547303 | 6598 | - |
| Chr05 | 16665952 | 16667992 | 2041 | - |
| Chr05 | 16992511 | 16999317 | 6807 | + |
| Chr05 | 17064645 | 17069748 | 5104 | + |
| Chr05 | 17256251 | 17266363 | 10113 | - |
| Chr05 | 17715377 | 17721795 | 6419 | + |
| Chr05 | 17789258 | 17810422 | 21165 | + |
| Chr05 | 17919223 | 17924407 | 5185 | + |
| Chr05 | 18508082 | 18529888 | 21807 | - |
| Chr05 | 18568267 | 18581814 | 13548 | + |
| Chr05 | 18618371 | 18628250 | 9880 | - |
| Chr05 | 18663283 | 18671902 | 8620 | + |
| Chr05 | 18663313 | 18675447 | 12135 | - |
| Chr05 | 18670391 | 18675447 | 5057 | - |
| Chr05 | 18759121 | 18764434 | 5314 | + |
| Chr05 | 18808531 | 18810812 | 2282 | + |
| Chr05 | 18920274 | 18927260 | 6987 | + |
| Chr05 | 19040831 | 19048283 | 7453 | - |
| Chr05 | 19049738 | 19063724 | 13987 | - |
| Chr05 | 19057841 | 19063234 | 5394 | - |
| Chr05 | 19153499 | 19158493 | 4995 | + |
| Chr05 | 19255370 | 19266266 | 10897 | - |
| Chr05 | 19268864 | 19286865 | 18002 | - |
| Chr05 | 19268864 | 19286886 | 18023 | - |
| Chr05 | 19269628 | 19294042 | 24415 | + |
| Chr05 | 19271573 | 19292886 | 21314 | + |
| Chr05 | 19278527 | 19285631 | 7105 | - |

| | | | | |
|---|---|---|---|---|
| Chr05 | 19285464 | 19295989 | 10526 | + |
| Chr05 | 19375088 | 19381863 | 6776 | - |
| Chr05 | 19628678 | 19638952 | 10275 | + |
| Chr05 | 19707846 | 19729002 | 21157 | + |
| Chr05 | 19707849 | 19729439 | 21591 | - |
| Chr05 | 19708227 | 19729475 | 21249 | + |
| Chr05 | 19824044 | 19831014 | 6971 | - |
| Chr05 | 20267555 | 20279650 | 12096 | + |
| Chr05 | 20416295 | 20427904 | 11610 | + |
| Chr05 | 20942991 | 20947888 | 4898 | + |
| Chr05 | 21134535 | 21160139 | 25605 | + |
| Chr05 | 21227999 | 21237878 | 9880 | - |
| Chr05 | 21733294 | 21736284 | 2991 | - |
| Chr05 | 22031397 | 22036106 | 4710 | + |
| Chr05 | 22136731 | 22146672 | 9942 | - |
| Chr05 | 22221496 | 22246968 | 25473 | - |
| Chr05 | 22225523 | 22242059 | 16537 | - |
| Chr05 | 22225809 | 22248543 | 22735 | - |
| Chr05 | 22322005 | 22327274 | 5270 | - |
| Chr05 | 22522926 | 22527016 | 4091 | - |
| Chr05 | 22652794 | 22657581 | 4788 | - |
| Chr05 | 22714435 | 22723686 | 9252 | + |
| Chr05 | 22957428 | 22959119 | 1692 | - |
| Chr05 | 22957630 | 22959119 | 1490 | - |
| Chr05 | 23144342 | 23149576 | 5235 | + |
| Chr05 | 23167622 | 23177709 | 10088 | - |
| Chr05 | 23772694 | 23784190 | 11497 | + |
| Chr05 | 23787575 | 23792963 | 5389 | - |
| Chr05 | 23797564 | 23808380 | 10817 | + |
| Chr05 | 23824750 | 23830193 | 5444 | + |
| Chr05 | 23973026 | 23977421 | 4396 | + |
| Chr05 | 23983854 | 23994147 | 10294 | + |
| Chr05 | 24052948 | 24058299 | 5352 | - |

| | | | | |
|---|---|---|---|---|
| Chr05 | 24115876 | 24121249 | 5374 | - |
| Chr05 | 24524950 | 24530387 | 5438 | + |
| Chr05 | 24658923 | 24663789 | 4867 | + |
| Chr05 | 25266876 | 25285052 | 18177 | + |
| Chr05 | 25289555 | 25294553 | 4999 | - |
| Chr05 | 25377535 | 25382803 | 5269 | + |
| Chr05 | 25541105 | 25546649 | 5545 | + |
| Chr05 | 25705220 | 25710269 | 5050 | + |
| Chr05 | 25793033 | 25799662 | 6630 | - |
| Chr05 | 25893962 | 25916680 | 22719 | + |
| Chr05 | 25919036 | 25931024 | 11989 | - |
| Chr05 | 25935435 | 25944305 | 8871 | - |
| Chr05 | 26036369 | 26041277 | 4909 | - |
| Chr05 | 26342515 | 26347961 | 5447 | + |
| Chr05 | 26422791 | 26428034 | 5244 | + |
| Chr05 | 26428290 | 26432573 | 4284 | + |
| Chr05 | 26501621 | 26507026 | 5406 | + |
| Chr05 | 26509999 | 26515080 | 5082 | + |
| Chr05 | 26556710 | 26561793 | 5084 | + |
| Chr05 | 26658173 | 26660052 | 1880 | + |
| Chr05 | 26783047 | 26788671 | 5625 | + |
| Chr05 | 26892004 | 26897780 | 5777 | + |
| Chr05 | 27039510 | 27044924 | 5415 | - |
| Chr05 | 27046172 | 27052547 | 6376 | - |
| Chr05 | 27145558 | 27154652 | 9095 | - |
| Chr05 | 27196063 | 27210820 | 14758 | - |
| Chr05 | 27286837 | 27297125 | 10289 | - |
| Chr05 | 27488689 | 27495293 | 6605 | - |
| Chr05 | 27525771 | 27538184 | 12414 | + |
| Chr05 | 27793669 | 27798603 | 4935 | + |
| Chr05 | 28288430 | 28290804 | 2375 | - |
| Chr05 | 28476454 | 28485827 | 9374 | + |
| Chr05 | 28559799 | 28571758 | 11960 | + |

| | | | | |
|---|---|---|---|---|
| Chr05 | 28628145 | 28637498 | 9354 | - |
| Chr05 | 29099186 | 29108000 | 8815 | + |
| Chr05 | 29100034 | 29103255 | 3222 | + |
| Chr05 | 29102882 | 29109101 | 6220 | - |
| Chr05 | 29118350 | 29128412 | 10063 | + |
| Chr05 | 29259776 | 29264855 | 5080 | - |
| Chr05 | 29399142 | 29404631 | 5490 | + |
| Chr05 | 29419358 | 29428153 | 8796 | + |
| Chr05 | 29499726 | 29505618 | 5893 | + |
| Chr05 | 29534772 | 29540231 | 5460 | - |
| Chr05 | 29638708 | 29645219 | 6512 | + |
| Chr05 | 29810076 | 29820940 | 10865 | - |
| Chr05 | 29965081 | 29975665 | 10585 | - |
| Chr05 | 30203580 | 30208162 | 4583 | - |
| Chr05 | 30714231 | 30717503 | 3273 | - |
| Chr05 | 30952548 | 30962336 | 9789 | - |
| Chr05 | 30952952 | 30962336 | 9385 | - |
| Chr05 | 31072384 | 31078016 | 5633 | + |
| Chr05 | 31130241 | 31131854 | 1614 | + |
| Chr05 | 31255749 | 31260511 | 4763 | - |
| Chr05 | 31777506 | 31789162 | 11657 | - |
| Chr05 | 31953006 | 31958243 | 5238 | - |
| Chr05 | 32406831 | 32412235 | 5405 | - |
| Chr05 | 33209698 | 33221350 | 11653 | - |
| Chr05 | 33303941 | 33310852 | 6912 | + |
| Chr05 | 33705175 | 33728734 | 23560 | - |
| Chr05 | 33960907 | 33966139 | 5233 | - |
| Chr05 | 33977850 | 33985460 | 7611 | + |
| Chr05 | 34365604 | 34378354 | 12751 | - |
| Chr05 | 34422737 | 34433501 | 10765 | - |
| Chr05 | 34622332 | 34632787 | 10456 | + |
| Chr05 | 34838168 | 34843410 | 5243 | - |
| Chr05 | 34868601 | 34880033 | 11433 | + |

| | | | | |
|---|---|---|---|---|
| Chr05 | 35450638 | 35455971 | 5334 | - |
| Chr05 | 36435483 | 36440949 | 5467 | + |
| Chr05 | 40326826 | 40331706 | 4881 | - |
| Chr05 | 40362677 | 40367492 | 4816 | + |
| Chr05 | 40464130 | 40470273 | 6144 | + |
| Chr05 | 42447834 | 42455432 | 7599 | - |
| Chr05 | 42448149 | 42455830 | 7682 | - |
| Chr05 | 43766641 | 43771608 | 4968 | - |
| Chr05 | 44344155 | 44354116 | 9962 | + |
| Chr05 | 44854213 | 44860517 | 6305 | + |
| Chr05 | 45355987 | 45362596 | 6610 | - |
| Chr06 | 160057 | 167032 | 6976 | - |
| Chr06 | 428895 | 434216 | 5322 | - |
| Chr06 | 699508 | 717028 | 17521 | + |
| Chr06 | 1006327 | 1013264 | 6938 | + |
| Chr06 | 1081432 | 1089024 | 7593 | + |
| Chr06 | 1152794 | 1157843 | 5050 | + |
| Chr06 | 1193829 | 1199369 | 5541 | - |
| Chr06 | 1214566 | 1223374 | 8809 | + |
| Chr06 | 1250818 | 1273330 | 22513 | + |
| Chr06 | 1252610 | 1262740 | 10131 | - |
| Chr06 | 1398571 | 1404477 | 5907 | + |
| Chr06 | 1563509 | 1571912 | 8404 | + |
| Chr06 | 1563509 | 1576772 | 13264 | + |
| Chr06 | 1670389 | 1675713 | 5325 | - |
| Chr06 | 1749934 | 1761015 | 11082 | - |
| Chr06 | 2157479 | 2162699 | 5221 | + |
| Chr06 | 2166291 | 2175288 | 8998 | - |
| Chr06 | 2263919 | 2273711 | 9793 | - |
| Chr06 | 2470018 | 2482826 | 12809 | + |
| Chr06 | 2653999 | 2658592 | 4594 | + |
| Chr06 | 2670660 | 2676284 | 5625 | + |
| Chr06 | 2821666 | 2827463 | 5798 | + |

| | | | | |
|---|---|---|---|---|
| Chr06 | 2992187 | 2994617 | 2431 | - |
| Chr06 | 3078049 | 3096881 | 18833 | + |
| Chr06 | 3098551 | 3103540 | 4990 | - |
| Chr06 | 3182304 | 3187370 | 5067 | - |
| Chr06 | 3363379 | 3369480 | 6102 | + |
| Chr06 | 3486061 | 3497175 | 11115 | + |
| Chr06 | 3487712 | 3496273 | 8562 | + |
| Chr06 | 3487712 | 3497175 | 9464 | + |
| Chr06 | 3487712 | 3497175 | 9464 | + |
| Chr06 | 3488618 | 3496273 | 7656 | + |
| Chr06 | 3488618 | 3496273 | 7656 | + |
| Chr06 | 3488618 | 3496273 | 7656 | + |
| Chr06 | 3996501 | 4000804 | 4304 | + |
| Chr06 | 3996501 | 4011977 | 15477 | + |
| Chr06 | 4083630 | 4089172 | 5543 | + |
| Chr06 | 4359700 | 4364664 | 4965 | + |
| Chr06 | 4617306 | 4621805 | 4500 | - |
| Chr06 | 4628601 | 4632725 | 4125 | + |
| Chr06 | 4645087 | 4650469 | 5383 | + |
| Chr06 | 5216710 | 5226476 | 9767 | + |
| Chr06 | 5271973 | 5278889 | 6917 | - |
| Chr06 | 6401080 | 6407743 | 6664 | - |
| Chr06 | 6442821 | 6445864 | 3044 | - |
| Chr06 | 6788647 | 6799699 | 11053 | + |
| Chr06 | 7123043 | 7134859 | 11817 | - |
| Chr06 | 7182660 | 7185866 | 3207 | - |
| Chr06 | 7412520 | 7417533 | 5014 | - |
| Chr06 | 7412520 | 7417704 | 5185 | - |
| Chr06 | 7532204 | 7537338 | 5135 | + |
| Chr06 | 7804325 | 7806720 | 2396 | + |
| Chr06 | 7817047 | 7822726 | 5680 | + |
| Chr06 | 7872588 | 7877956 | 5369 | - |
| Chr06 | 8040248 | 8043954 | 3707 | + |

| | | | | |
|---|---|---|---|---|
| Chr06 | 8234444 | 8245294 | 10851 | - |
| Chr06 | 8362026 | 8364633 | 2608 | + |
| Chr06 | 8362026 | 8374427 | 12402 | + |
| Chr06 | 8630699 | 8634723 | 4025 | + |
| Chr06 | 8908517 | 8912879 | 4363 | + |
| Chr06 | 9334554 | 9347077 | 12524 | + |
| Chr06 | 9522551 | 9526473 | 3923 | + |
| Chr06 | 9532085 | 9537261 | 5177 | + |
| Chr06 | 9609364 | 9619647 | 10284 | + |
| Chr06 | 9610733 | 9615883 | 5151 | - |
| Chr06 | 9610733 | 9634054 | 23322 | - |
| Chr06 | 9627527 | 9637818 | 10292 | + |
| Chr06 | 9628898 | 9634054 | 5157 | - |
| Chr06 | 9712521 | 9717723 | 5203 | - |
| Chr06 | 9770752 | 9776098 | 5347 | + |
| Chr06 | 9928123 | 9933550 | 5428 | - |
| Chr06 | 10154547 | 10161786 | 7240 | + |
| Chr06 | 10199092 | 10212276 | 13185 | + |
| Chr06 | 10751969 | 10766714 | 14746 | - |
| Chr06 | 10810385 | 10815723 | 5339 | - |
| Chr06 | 11543961 | 11558247 | 14287 | - |
| Chr06 | 11643998 | 11648787 | 4790 | - |
| Chr06 | 11674152 | 11679373 | 5222 | + |
| Chr06 | 11898110 | 11902712 | 4603 | - |
| Chr06 | 12353783 | 12362805 | 9023 | + |
| Chr06 | 12684847 | 12692422 | 7576 | + |
| Chr06 | 14249651 | 14256734 | 7084 | + |
| Chr06 | 14309206 | 14314707 | 5502 | + |
| Chr06 | 14470699 | 14482645 | 11947 | + |
| Chr06 | 14471612 | 14478286 | 6675 | - |
| Chr06 | 14718079 | 14721098 | 3020 | + |
| Chr06 | 15342504 | 15347421 | 4918 | + |
| Chr06 | 15449087 | 15460815 | 11729 | - |

| | | | | |
|---|---|---|---|---|
| Chr06 | 15586553 | 15597107 | 10555 | + |
| Chr06 | 15586553 | 15599215 | 12663 | + |
| Chr06 | 15586553 | 15599391 | 12839 | + |
| Chr06 | 16898389 | 16907265 | 8877 | + |
| Chr06 | 17283881 | 17300758 | 16878 | + |
| Chr06 | 17307021 | 17324089 | 17069 | + |
| Chr06 | 17867584 | 17874385 | 6802 | + |
| Chr06 | 19282666 | 19288228 | 5563 | - |
| Chr06 | 20018350 | 20028541 | 10192 | + |
| Chr06 | 20105239 | 20112190 | 6952 | + |
| Chr06 | 20114874 | 20122787 | 7914 | - |
| Chr06 | 20155876 | 20161960 | 6085 | - |
| Chr06 | 20233131 | 20239719 | 6589 | - |
| Chr06 | 20303922 | 20311394 | 7473 | + |
| Chr06 | 20509223 | 20522838 | 13616 | + |
| Chr06 | 20544542 | 20552127 | 7586 | - |
| Chr06 | 21018056 | 21037605 | 19550 | + |
| Chr06 | 21032606 | 21037605 | 5000 | + |
| Chr06 | 21806007 | 21817353 | 11347 | - |
| Chr07 | 1072998 | 1083839 | 10842 | - |
| Chr07 | 1575745 | 1595742 | 19998 | + |
| Chr07 | 2473116 | 2478421 | 5306 | + |
| Chr07 | 2556690 | 2562302 | 5613 | - |
| Chr07 | 2563994 | 2566065 | 2072 | + |
| Chr07 | 2604233 | 2612061 | 7829 | + |
| Chr07 | 2777381 | 2784671 | 7291 | + |
| Chr07 | 2862367 | 2867331 | 4965 | + |
| Chr07 | 2951875 | 2962377 | 10503 | - |
| Chr07 | 3103048 | 3107939 | 4892 | + |
| Chr07 | 3522770 | 3528110 | 5341 | - |
| Chr07 | 3529631 | 3540180 | 10550 | - |
| Chr07 | 4773898 | 4777732 | 3835 | - |
| Chr07 | 4924970 | 4928822 | 3853 | + |

| | | | | |
|---|---|---|---|---|
| Chr07 | 5136442 | 5141501 | 5060 | + |
| Chr07 | 5579325 | 5586331 | 7007 | + |
| Chr07 | 5750271 | 5755882 | 5612 | - |
| Chr07 | 6198502 | 6201789 | 3288 | + |
| Chr07 | 6355112 | 6361935 | 6824 | + |
| Chr07 | 6505583 | 6510590 | 5008 | + |
| Chr07 | 7802942 | 7804896 | 1955 | + |
| Chr07 | 7813727 | 7822373 | 8647 | + |
| Chr07 | 7833490 | 7838465 | 4976 | - |
| Chr07 | 8074906 | 8081734 | 6829 | + |
| Chr07 | 8156041 | 8159575 | 3535 | + |
| Chr07 | 8194400 | 8206462 | 12063 | + |
| Chr07 | 8239952 | 8245016 | 5065 | + |
| Chr07 | 8286252 | 8295188 | 8937 | + |
| Chr07 | 8286252 | 8307547 | 21296 | + |
| Chr07 | 8466802 | 8469569 | 2768 | - |
| Chr07 | 8646771 | 8667687 | 20917 | - |
| Chr07 | 8808802 | 8814459 | 5658 | - |
| Chr07 | 9240954 | 9251958 | 11005 | + |
| Chr07 | 9432608 | 9435681 | 3074 | - |
| Chr07 | 9935049 | 9937106 | 2058 | - |
| Chr07 | 10224834 | 10231268 | 6435 | - |
| Chr07 | 10283469 | 10288613 | 5145 | - |
| Chr07 | 10301974 | 10305599 | 3626 | + |
| Chr07 | 10761941 | 10766596 | 4656 | + |
| Chr07 | 10920840 | 10934225 | 13386 | + |
| Chr07 | 11280249 | 11294472 | 14224 | + |
| Chr07 | 11332245 | 11338776 | 6532 | - |
| Chr07 | 11430060 | 11442240 | 12181 | + |
| Chr07 | 11556036 | 11564377 | 8342 | - |
| Chr07 | 11721018 | 11733616 | 12599 | + |
| Chr07 | 11728086 | 11748422 | 20337 | + |
| Chr07 | 11731821 | 11750247 | 18427 | - |

| | | | | |
|---|---|---|---|---|
| Chr07 | 11737448 | 11746116 | 8669 | - |
| Chr07 | 11751116 | 11756290 | 5175 | - |
| Chr07 | 11879842 | 11885081 | 5240 | - |
| Chr07 | 12155857 | 12164751 | 8895 | + |
| Chr07 | 12196821 | 12202351 | 5531 | + |
| Chr07 | 12204647 | 12209238 | 4592 | - |
| Chr07 | 12352697 | 12365437 | 12741 | + |
| Chr07 | 12409803 | 12435126 | 25324 | - |
| Chr07 | 12410700 | 12435126 | 24427 | - |
| Chr07 | 12442240 | 12447261 | 5022 | - |
| Chr07 | 12956641 | 12967926 | 11286 | + |
| Chr07 | 13407983 | 13418762 | 10780 | - |
| Chr07 | 13577311 | 13588049 | 10739 | - |
| Chr07 | 14574199 | 14580302 | 6104 | - |
| Chr07 | 14973612 | 14997171 | 23560 | + |
| Chr07 | 16059460 | 16064891 | 5432 | + |
| Chr07 | 16954871 | 16958589 | 3719 | - |
| Chr07 | 17055703 | 17057927 | 2225 | - |
| Chr07 | 17729188 | 17741978 | 12791 | + |
| Chr07 | 17886691 | 17896216 | 9526 | + |
| Chr07 | 19254224 | 19256858 | 2635 | + |
| Chr07 | 20382560 | 20388158 | 5599 | - |
| Chr07 | 20398573 | 20400423 | 1851 | - |
| Chr07 | 20639844 | 20649900 | 10057 | - |
| Chr07 | 20819576 | 20823725 | 4150 | - |
| Chr07 | 21290183 | 21296899 | 6717 | + |
| Chr08 | 346742 | 351707 | 4966 | - |
| Chr08 | 398361 | 407117 | 8757 | + |
| Chr08 | 664597 | 683250 | 18654 | - |
| Chr08 | 725025 | 735921 | 10897 | + |
| Chr08 | 1351221 | 1353321 | 2101 | + |
| Chr08 | 1384353 | 1396429 | 12077 | + |
| Chr08 | 1678048 | 1690023 | 11976 | + |

| | | | | |
|---|---|---|---|---|
| Chr08 | 1725393 | 1731194 | 5802 | + |
| Chr08 | 1796586 | 1814691 | 18106 | - |
| Chr08 | 1809131 | 1822965 | 13835 | + |
| Chr08 | 1809141 | 1823276 | 14136 | + |
| Chr08 | 1813053 | 1823276 | 10224 | + |
| Chr08 | 1935210 | 1940297 | 5088 | - |
| Chr08 | 2462195 | 2464005 | 1811 | + |
| Chr08 | 2990206 | 3000574 | 10369 | + |
| Chr08 | 3546719 | 3560120 | 13402 | + |
| Chr08 | 3682077 | 3686900 | 4824 | + |
| Chr08 | 4297534 | 4302987 | 5454 | + |
| Chr08 | 4331148 | 4339554 | 8407 | - |
| Chr08 | 4371337 | 4377819 | 6483 | - |
| Chr08 | 4504033 | 4507273 | 3241 | + |
| Chr08 | 4581996 | 4588419 | 6424 | + |
| Chr08 | 5339829 | 5351192 | 11364 | - |
| Chr08 | 5493331 | 5498073 | 4743 | - |
| Chr08 | 5671917 | 5677311 | 5395 | - |
| Chr08 | 6956982 | 6971874 | 14893 | - |
| Chr08 | 7326274 | 7336250 | 9977 | - |
| Chr08 | 7439275 | 7449642 | 10368 | + |
| Chr08 | 7520538 | 7542563 | 22026 | - |
| Chr08 | 7731915 | 7736971 | 5057 | + |
| Chr08 | 7874016 | 7879589 | 5574 | + |
| Chr08 | 7912469 | 7921360 | 8892 | + |
| Chr08 | 8016537 | 8023528 | 6992 | - |
| Chr08 | 8196280 | 8201840 | 5561 | - |
| Chr08 | 8207243 | 8218957 | 11715 | - |
| Chr08 | 8279179 | 8284543 | 5365 | + |
| Chr08 | 8290035 | 8294927 | 4893 | + |
| Chr08 | 8581854 | 8583843 | 1990 | - |
| Chr08 | 8617640 | 8621929 | 4290 | + |
| Chr08 | 8685134 | 8690370 | 5237 | - |

| | | | | |
|---|---|---|---|---|
| Chr08 | 8695537 | 8708754 | 13218 | - |
| Chr08 | 8698281 | 8705188 | 6908 | - |
| Chr08 | 8867644 | 8888558 | 20915 | + |
| Chr08 | 9031636 | 9036829 | 5194 | + |
| Chr08 | 9265103 | 9273933 | 8831 | - |
| Chr08 | 9589383 | 9594392 | 5010 | + |
| Chr08 | 9647964 | 9656836 | 8873 | - |
| Chr08 | 9764172 | 9769849 | 5678 | - |
| Chr08 | 10289183 | 10294592 | 5410 | - |
| Chr08 | 10660957 | 10666069 | 5113 | - |
| Chr08 | 10700711 | 10706452 | 5742 | + |
| Chr08 | 10820432 | 10827506 | 7075 | - |
| Chr08 | 10920726 | 10925979 | 5254 | + |
| Chr08 | 11005114 | 11010540 | 5427 | + |
| Chr08 | 11240220 | 11245257 | 5038 | + |
| Chr08 | 11252324 | 11257226 | 4903 | + |
| Chr08 | 11375244 | 11406245 | 31002 | + |
| Chr08 | 11377437 | 11406245 | 28809 | + |
| Chr08 | 11380143 | 11403968 | 23826 | + |
| Chr08 | 11381224 | 11403968 | 22745 | + |
| Chr08 | 11381945 | 11404471 | 22527 | + |
| Chr08 | 11382302 | 11404129 | 21828 | + |
| Chr08 | 11382482 | 11403968 | 21487 | + |
| Chr08 | 11382662 | 11403787 | 21126 | + |
| Chr08 | 11383383 | 11403065 | 19683 | + |
| Chr08 | 11390947 | 11421800 | 30854 | + |
| Chr08 | 11390947 | 11421947 | 31001 | + |
| Chr08 | 11390947 | 11422079 | 31133 | + |
| Chr08 | 11391308 | 11412299 | 20992 | + |
| Chr08 | 11392197 | 11411413 | 19217 | + |
| Chr08 | 11402893 | 11406220 | 3328 | + |
| Chr08 | 11402893 | 11408002 | 5110 | + |
| Chr08 | 11402893 | 11408905 | 6013 | + |

| | | | | |
|---|---|---|---|---|
| Chr08 | 11402893 | 11408905 | 6013 | + |
| Chr08 | 11402893 | 11410929 | 8037 | + |
| Chr08 | 11402893 | 11412886 | 9994 | + |
| Chr08 | 11402893 | 11415378 | 12486 | + |
| Chr08 | 11402893 | 11415736 | 12844 | + |
| Chr08 | 11402893 | 11416605 | 13713 | + |
| Chr08 | 11402893 | 11417140 | 14248 | + |
| Chr08 | 11402893 | 11417674 | 14782 | + |
| Chr08 | 11402893 | 11418564 | 15672 | + |
| Chr08 | 11402893 | 11418890 | 15998 | + |
| Chr08 | 11402893 | 11419069 | 16177 | + |
| Chr08 | 11402893 | 11419069 | 16177 | + |
| Chr08 | 11402893 | 11420262 | 17370 | + |
| Chr08 | 11402893 | 11421296 | 18404 | + |
| Chr08 | 11402893 | 11422136 | 19244 | + |
| Chr08 | 11402893 | 11422136 | 19244 | + |
| Chr08 | 11402893 | 11423166 | 20274 | + |
| Chr08 | 11402893 | 11423342 | 20450 | + |
| Chr08 | 11402893 | 11426080 | 23188 | + |
| Chr08 | 11402893 | 11426080 | 23188 | + |
| Chr08 | 11403602 | 11422160 | 18559 | + |
| Chr08 | 11403782 | 11410954 | 7173 | + |
| Chr08 | 11403963 | 11422992 | 19030 | + |
| Chr08 | 11404124 | 11423493 | 19370 | + |
| Chr08 | 11404647 | 11420285 | 15639 | + |
| Chr08 | 11404647 | 11421800 | 17154 | + |
| Chr08 | 11404647 | 11422992 | 18346 | + |
| Chr08 | 11404647 | 11424515 | 19869 | + |
| Chr08 | 11404808 | 11421947 | 17140 | + |
| Chr08 | 11404808 | 11424061 | 19254 | + |
| Chr08 | 11405201 | 11422160 | 16960 | + |
| Chr08 | 11405201 | 11424515 | 19315 | + |
| Chr08 | 11405201 | 11424899 | 19699 | + |

| | | | | |
|---|---|---|---|---|
| Chr08 | 11405563 | 11419933 | 14371 | + |
| Chr08 | 11405563 | 11420593 | 15031 | + |
| Chr08 | 11405720 | 11423493 | 17774 | + |
| Chr08 | 11405880 | 11422992 | 17113 | + |
| Chr08 | 11405880 | 11423493 | 17614 | + |
| Chr08 | 11405880 | 11424515 | 18636 | + |
| Chr08 | 11405880 | 11425914 | 20035 | + |
| Chr08 | 11406059 | 11422464 | 16406 | + |
| Chr08 | 11406059 | 11422963 | 16905 | + |
| Chr08 | 11406240 | 11422963 | 16724 | + |
| Chr08 | 11406251 | 11422992 | 16742 | + |
| Chr08 | 11406251 | 11425357 | 19107 | + |
| Chr08 | 11406400 | 11421947 | 15548 | + |
| Chr08 | 11406400 | 11422992 | 16593 | + |
| Chr08 | 11406400 | 11423464 | 17065 | + |
| Chr08 | 11406580 | 11420593 | 14014 | + |
| Chr08 | 11406580 | 11421947 | 15368 | + |
| Chr08 | 11406580 | 11424061 | 17482 | + |
| Chr08 | 11406937 | 11421800 | 14864 | + |
| Chr08 | 11406937 | 11422257 | 15321 | + |
| Chr08 | 11406937 | 11423493 | 16557 | + |
| Chr08 | 11406937 | 11424515 | 17579 | + |
| Chr08 | 11406937 | 11424899 | 17963 | + |
| Chr08 | 11406937 | 11425177 | 18241 | + |
| Chr08 | 11407299 | 11422992 | 15694 | + |
| Chr08 | 11407299 | 11423367 | 16069 | + |
| Chr08 | 11407660 | 11423012 | 15353 | + |
| Chr08 | 11408394 | 11415402 | 7009 | + |
| Chr08 | 11408564 | 11422160 | 13597 | + |
| Chr08 | 11409168 | 11418715 | 9548 | + |
| Chr08 | 11409168 | 11418715 | 9548 | + |
| Chr08 | 11409168 | 11419894 | 10727 | + |
| Chr08 | 11409168 | 11419923 | 10756 | + |

| | | | | |
|---|---|---|---|---|
| Chr08 | 11409168 | 11420593 | 11426 | + |
| Chr08 | 11409168 | 11420593 | 11426 | + |
| Chr08 | 11409168 | 11420593 | 11426 | + |
| Chr08 | 11409168 | 11421800 | 12633 | + |
| Chr08 | 11409168 | 11422992 | 13825 | + |
| Chr08 | 11409168 | 11422992 | 13825 | + |
| Chr08 | 11409168 | 11422992 | 13825 | + |
| Chr08 | 11409168 | 11423464 | 14297 | + |
| Chr08 | 11409168 | 11423464 | 14297 | + |
| Chr08 | 11409168 | 11423493 | 14326 | + |
| Chr08 | 11409168 | 11423493 | 14326 | + |
| Chr08 | 11409168 | 11423493 | 14326 | + |
| Chr08 | 11409168 | 11424486 | 15319 | + |
| Chr08 | 11409168 | 11424486 | 15319 | + |
| Chr08 | 11409168 | 11424486 | 15319 | + |
| Chr08 | 11409168 | 11424515 | 15348 | + |
| Chr08 | 11409168 | 11424515 | 15348 | + |
| Chr08 | 11409168 | 11424515 | 15348 | + |
| Chr08 | 11409168 | 11424515 | 15348 | + |
| Chr08 | 11409168 | 11425386 | 16219 | + |
| Chr08 | 11409696 | 11422992 | 13297 | + |
| Chr08 | 11409696 | 11424515 | 14820 | + |
| Chr08 | 11410225 | 11423493 | 13269 | + |
| Chr08 | 11410225 | 11424486 | 14262 | + |
| Chr08 | 11410587 | 11421947 | 11361 | + |
| Chr08 | 11410587 | 11424515 | 13929 | + |
| Chr08 | 11410768 | 11420593 | 9826 | + |
| Chr08 | 11410768 | 11421771 | 11004 | + |
| Chr08 | 11410768 | 11421800 | 11033 | + |
| Chr08 | 11410768 | 11422435 | 11668 | + |
| Chr08 | 11410768 | 11422963 | 12196 | + |
| Chr08 | 11410768 | 11422992 | 12225 | + |
| Chr08 | 11410768 | 11423493 | 12726 | + |

| | | | | |
|---|---|---|---|---|
| Chr08 | 11411310 | 11421947 | 10638 | + |
| Chr08 | 11411490 | 11423493 | 12004 | + |
| Chr08 | 11411490 | 11424486 | 12997 | + |
| Chr08 | 11412196 | 11423464 | 11269 | + |
| Chr08 | 11414221 | 11447724 | 33504 | + |
| Chr08 | 11414513 | 11420593 | 6081 | + |
| Chr08 | 11414513 | 11421810 | 7298 | + |
| Chr08 | 11414513 | 11422478 | 7966 | + |
| Chr08 | 11414513 | 11422963 | 8451 | + |
| Chr08 | 11414513 | 11422992 | 8480 | + |
| Chr08 | 11414513 | 11422992 | 8480 | + |
| Chr08 | 11414513 | 11423464 | 8952 | + |
| Chr08 | 11414513 | 11423493 | 8981 | + |
| Chr08 | 11414513 | 11423493 | 8981 | + |
| Chr08 | 11414513 | 11424187 | 9675 | + |
| Chr08 | 11414513 | 11424486 | 9974 | + |
| Chr08 | 11414513 | 11424486 | 9974 | + |
| Chr08 | 11414513 | 11424515 | 10003 | + |
| Chr08 | 11414513 | 11424515 | 10003 | + |
| Chr08 | 11415038 | 11421771 | 6734 | + |
| Chr08 | 11415038 | 11422992 | 7955 | + |
| Chr08 | 11415038 | 11422992 | 7955 | + |
| Chr08 | 11415038 | 11423285 | 8248 | + |
| Chr08 | 11415038 | 11423464 | 8427 | + |
| Chr08 | 11415038 | 11424515 | 9478 | + |
| Chr08 | 11415038 | 11426564 | 11527 | + |
| Chr08 | 11415218 | 11420593 | 5376 | + |
| Chr08 | 11415218 | 11421417 | 6200 | + |
| Chr08 | 11415218 | 11421800 | 6583 | + |
| Chr08 | 11415218 | 11421947 | 6730 | + |
| Chr08 | 11415397 | 11421800 | 6404 | + |
| Chr08 | 11415397 | 11421947 | 6551 | + |
| Chr08 | 11415397 | 11422464 | 7068 | + |

| | | | | |
|---|---|---|---|---|
| Chr08 | 11415397 | 11422963 | 7567 | + |
| Chr08 | 11415577 | 11420593 | 5017 | + |
| Chr08 | 11415577 | 11421620 | 6044 | + |
| Chr08 | 11415577 | 11421771 | 6195 | + |
| Chr08 | 11415577 | 11421800 | 6224 | + |
| Chr08 | 11415577 | 11422963 | 7387 | + |
| Chr08 | 11415577 | 11422963 | 7387 | + |
| Chr08 | 11415577 | 11422992 | 7416 | + |
| Chr08 | 11415577 | 11422992 | 7416 | + |
| Chr08 | 11415577 | 11423493 | 7917 | + |
| Chr08 | 11415577 | 11423493 | 7917 | + |
| Chr08 | 11415577 | 11423493 | 7917 | + |
| Chr08 | 11415577 | 11424486 | 8910 | + |
| Chr08 | 11415577 | 11424515 | 8939 | + |
| Chr08 | 11415755 | 11420603 | 4849 | + |
| Chr08 | 11415755 | 11421417 | 5663 | + |
| Chr08 | 11415755 | 11421771 | 6017 | + |
| Chr08 | 11415755 | 11421800 | 6046 | + |
| Chr08 | 11415755 | 11421810 | 6056 | + |
| Chr08 | 11415755 | 11421947 | 6193 | + |
| Chr08 | 11415755 | 11421947 | 6193 | + |
| Chr08 | 11415755 | 11421947 | 6193 | + |
| Chr08 | 11415755 | 11422478 | 6724 | + |
| Chr08 | 11415755 | 11422602 | 6848 | + |
| Chr08 | 11415755 | 11422992 | 7238 | + |
| Chr08 | 11415755 | 11423464 | 7710 | + |
| Chr08 | 11415755 | 11423493 | 7739 | + |
| Chr08 | 11415755 | 11424187 | 8433 | + |
| Chr08 | 11415755 | 11424486 | 8732 | + |
| Chr08 | 11415755 | 11424515 | 8761 | + |
| Chr08 | 11415755 | 11424515 | 8761 | + |
| Chr08 | 11415933 | 11421771 | 5839 | + |
| Chr08 | 11415933 | 11421800 | 5868 | + |

| | | | | |
|---|---|---|---|---|
| Chr08 | 11415933 | 11422992 | 7060 | + |
| Chr08 | 11416473 | 11422257 | 5785 | + |
| Chr08 | 11417159 | 11421771 | 4613 | + |
| Chr08 | 11417159 | 11423464 | 6306 | + |
| Chr08 | 11417159 | 11425714 | 8556 | + |
| Chr08 | 11417159 | 11427639 | 10481 | + |
| Chr08 | 11417337 | 11421771 | 4435 | + |
| Chr08 | 11417337 | 11422992 | 5656 | + |
| Chr08 | 11417337 | 11423493 | 6157 | + |
| Chr08 | 11417337 | 11424515 | 7179 | + |
| Chr08 | 11417337 | 11425026 | 7690 | + |
| Chr08 | 11417337 | 11425386 | 8050 | + |
| Chr08 | 11417515 | 11420593 | 3079 | + |
| Chr08 | 11417515 | 11422478 | 4964 | + |
| Chr08 | 11417515 | 11423464 | 5950 | + |
| Chr08 | 11417515 | 11424486 | 6972 | + |
| Chr08 | 11417515 | 11424486 | 6972 | + |
| Chr08 | 11417515 | 11424515 | 7001 | + |
| Chr08 | 11417515 | 11424515 | 7001 | + |
| Chr08 | 11417869 | 11420603 | 2735 | + |
| Chr08 | 11417869 | 11420916 | 3048 | + |
| Chr08 | 11418049 | 11423493 | 5445 | + |
| Chr08 | 11418049 | 11424486 | 6438 | + |
| Chr08 | 11418049 | 11425533 | 7485 | + |
| Chr08 | 11418229 | 11422257 | 4029 | + |
| Chr08 | 11418584 | 11420233 | 1650 | + |
| Chr08 | 11418584 | 11423493 | 4910 | + |
| Chr08 | 11418584 | 11425026 | 6443 | + |
| Chr08 | 11418720 | 11421771 | 3052 | + |
| Chr08 | 11418720 | 11422464 | 3745 | + |
| Chr08 | 11418720 | 11422992 | 4273 | + |
| Chr08 | 11418720 | 11425386 | 6667 | + |
| Chr08 | 11418720 | 11428170 | 9451 | + |

| | | | | |
|---|---|---|---|---|
| Chr08 | 11418720 | 11428199 | 9480 | + |
| Chr08 | 11418720 | 11428921 | 10202 | + |
| Chr08 | 11418720 | 11429459 | 10740 | + |
| Chr08 | 11418720 | 11435053 | 16334 | + |
| Chr08 | 11418720 | 11435592 | 16873 | + |
| Chr08 | 11418720 | 11441835 | 23116 | + |
| Chr08 | 11418720 | 11443101 | 24382 | + |
| Chr08 | 11418720 | 11445246 | 26527 | + |
| Chr08 | 11418720 | 11445759 | 27040 | + |
| Chr08 | 11418720 | 11451439 | 32720 | + |
| Chr08 | 11418720 | 11453227 | 34508 | + |
| Chr08 | 11418745 | 11447722 | 28978 | + |
| Chr08 | 11418758 | 11422801 | 4044 | + |
| Chr08 | 11419088 | 11422992 | 3905 | + |
| Chr08 | 11419088 | 11423493 | 4406 | + |
| Chr08 | 11419116 | 11428189 | 9074 | + |
| Chr08 | 11419435 | 11451439 | 32005 | + |
| Chr08 | 11419793 | 11423493 | 3701 | + |
| Chr08 | 11419793 | 11424515 | 4723 | + |
| Chr08 | 11419793 | 11425026 | 5234 | + |
| Chr08 | 11419948 | 11435592 | 15645 | + |
| Chr08 | 11419948 | 11437569 | 17622 | + |
| Chr08 | 11419948 | 11446458 | 26511 | + |
| Chr08 | 11419948 | 11447029 | 27082 | + |
| Chr08 | 11419948 | 11448084 | 28137 | + |
| Chr08 | 11419948 | 11450020 | 30073 | + |
| Chr08 | 11419948 | 11450935 | 30988 | + |
| Chr08 | 11419948 | 11451439 | 31492 | + |
| Chr08 | 11419948 | 11451648 | 31701 | + |
| Chr08 | 11419948 | 11452546 | 32599 | + |
| Chr08 | 11420100 | 11421771 | 1672 | + |
| Chr08 | 11420100 | 11422992 | 2893 | + |
| Chr08 | 11420100 | 11422992 | 2893 | + |

| | | | | |
|---|---|---|---|---|
| Chr08 | 11420100 | 11422992 | 2893 | + |
| Chr08 | 11420100 | 11423464 | 3365 | + |
| Chr08 | 11420100 | 11423493 | 3394 | + |
| Chr08 | 11420100 | 11423493 | 3394 | + |
| Chr08 | 11420100 | 11424486 | 4387 | + |
| Chr08 | 11420100 | 11424515 | 4416 | + |
| Chr08 | 11420100 | 11425026 | 4927 | + |
| Chr08 | 11420100 | 11425533 | 5434 | + |
| Chr08 | 11420100 | 11427471 | 7372 | + |
| Chr08 | 11420100 | 11428380 | 8281 | + |
| Chr08 | 11420100 | 11428921 | 8822 | + |
| Chr08 | 11420100 | 11428921 | 8822 | + |
| Chr08 | 11420100 | 11429100 | 9001 | + |
| Chr08 | 11420100 | 11429459 | 9360 | + |
| Chr08 | 11420100 | 11429459 | 9360 | + |
| Chr08 | 11420100 | 11429639 | 9540 | + |
| Chr08 | 11420100 | 11429819 | 9720 | + |
| Chr08 | 11420100 | 11432025 | 11926 | + |
| Chr08 | 11420100 | 11433281 | 13182 | + |
| Chr08 | 11420100 | 11434514 | 14415 | + |
| Chr08 | 11420100 | 11435053 | 14954 | + |
| Chr08 | 11420100 | 11435592 | 15493 | + |
| Chr08 | 11420100 | 11436134 | 16035 | + |
| Chr08 | 11420100 | 11437030 | 16931 | + |
| Chr08 | 11420100 | 11437210 | 17111 | + |
| Chr08 | 11420100 | 11437569 | 17470 | + |
| Chr08 | 11420100 | 11437569 | 17470 | + |
| Chr08 | 11420100 | 11437749 | 17650 | + |
| Chr08 | 11420100 | 11438472 | 18373 | + |
| Chr08 | 11420100 | 11438833 | 18734 | + |
| Chr08 | 11420100 | 11438833 | 18734 | + |
| Chr08 | 11420100 | 11440583 | 20484 | + |
| Chr08 | 11420100 | 11440943 | 20844 | + |

| | | | | |
|---|---|---|---|---|
| Chr08 | 11420100 | 11442710 | 22611 | + |
| Chr08 | 11420100 | 11442739 | 22640 | + |
| Chr08 | 11420100 | 11442739 | 22640 | + |
| Chr08 | 11420100 | 11443101 | 23002 | + |
| Chr08 | 11420100 | 11443270 | 23171 | + |
| Chr08 | 11420100 | 11443422 | 23323 | + |
| Chr08 | 11420100 | 11444343 | 24244 | + |
| Chr08 | 11420100 | 11444495 | 24396 | + |
| Chr08 | 11420100 | 11444886 | 24787 | + |
| Chr08 | 11420100 | 11445040 | 24941 | + |
| Chr08 | 11420100 | 11445246 | 25147 | + |
| Chr08 | 11420100 | 11445246 | 25147 | + |
| Chr08 | 11420100 | 11445246 | 25147 | + |
| Chr08 | 11420100 | 11445578 | 25479 | + |
| Chr08 | 11420100 | 11445759 | 25660 | + |
| Chr08 | 11420100 | 11445759 | 25660 | + |
| Chr08 | 11420100 | 11445759 | 25660 | + |
| Chr08 | 11420100 | 11445759 | 25660 | + |
| Chr08 | 11420100 | 11445928 | 25829 | + |
| Chr08 | 11420100 | 11445957 | 25858 | + |
| Chr08 | 11420100 | 11445957 | 25858 | + |
| Chr08 | 11420100 | 11446283 | 26184 | + |
| Chr08 | 11420100 | 11446458 | 26359 | + |
| Chr08 | 11420100 | 11446458 | 26359 | + |
| Chr08 | 11420100 | 11446843 | 26744 | + |
| Chr08 | 11420100 | 11446843 | 26744 | + |
| Chr08 | 11420100 | 11447029 | 26930 | + |
| Chr08 | 11420100 | 11447358 | 27259 | + |
| Chr08 | 11420100 | 11447904 | 27805 | + |
| Chr08 | 11420100 | 11447904 | 27805 | + |
| Chr08 | 11420100 | 11448084 | 27985 | + |
| Chr08 | 11420100 | 11448601 | 28502 | + |
| Chr08 | 11420100 | 11448804 | 28705 | + |

| | | | | |
|---|---|---|---|---|
| Chr08 | 11420100 | 11448953 | 28854 | + |
| Chr08 | 11420100 | 11449303 | 29204 | + |
| Chr08 | 11420100 | 11449331 | 29232 | + |
| Chr08 | 11420100 | 11449480 | 29381 | + |
| Chr08 | 11420100 | 11449689 | 29590 | + |
| Chr08 | 11420100 | 11449869 | 29770 | + |
| Chr08 | 11420100 | 11450228 | 30129 | + |
| Chr08 | 11420100 | 11450755 | 30656 | + |
| Chr08 | 11420100 | 11450906 | 30807 | + |
| Chr08 | 11420100 | 11450935 | 30836 | + |
| Chr08 | 11420100 | 11451828 | 31729 | + |
| Chr08 | 11420608 | 11423493 | 2886 | + |
| Chr08 | 11420608 | 11424515 | 3908 | + |
| Chr08 | 11420608 | 11424515 | 3908 | + |
| Chr08 | 11420608 | 11424515 | 3908 | + |
| Chr08 | 11420608 | 11425026 | 4419 | + |
| Chr08 | 11420608 | 11425357 | 4750 | + |
| Chr08 | 11420608 | 11425406 | 4799 | + |
| Chr08 | 11420608 | 11425533 | 4926 | + |
| Chr08 | 11420608 | 11425714 | 5107 | + |
| Chr08 | 11420608 | 11426412 | 5805 | + |
| Chr08 | 11420608 | 11427639 | 7032 | + |
| Chr08 | 11420608 | 11428921 | 8314 | + |
| Chr08 | 11420608 | 11429100 | 8493 | + |
| Chr08 | 11420608 | 11429459 | 8852 | + |
| Chr08 | 11420608 | 11429819 | 9212 | + |
| Chr08 | 11420608 | 11432919 | 12312 | + |
| Chr08 | 11420608 | 11434693 | 14086 | + |
| Chr08 | 11420608 | 11435234 | 14627 | + |
| Chr08 | 11420608 | 11435743 | 15136 | + |
| Chr08 | 11420608 | 11436822 | 16215 | + |
| Chr08 | 11420608 | 11437210 | 16603 | + |
| Chr08 | 11420608 | 11437749 | 17142 | + |

| | | | | |
|---|---|---|---|---|
| Chr08 | 11420608 | 11438472 | 17865 | + |
| Chr08 | 11420608 | 11438624 | 18017 | + |
| Chr08 | 11420608 | 11438833 | 18226 | + |
| Chr08 | 11420608 | 11440583 | 19976 | + |
| Chr08 | 11420623 | 11447722 | 27100 | + |
| Chr08 | 11420963 | 11423464 | 2502 | + |
| Chr08 | 11421669 | 11423110 | 1442 | + |
| Chr08 | 11421817 | 11424486 | 2670 | + |
| Chr08 | 11421817 | 11425026 | 3210 | + |
| Chr08 | 11421817 | 11425177 | 3361 | + |
| Chr08 | 11421817 | 11425386 | 3570 | + |
| Chr08 | 11421817 | 11425533 | 3717 | + |
| Chr08 | 11421817 | 11425533 | 3717 | + |
| Chr08 | 11421817 | 11425714 | 3898 | + |
| Chr08 | 11421817 | 11426051 | 4235 | + |
| Chr08 | 11421817 | 11427668 | 5852 | + |
| Chr08 | 11421817 | 11428002 | 6186 | + |
| Chr08 | 11421817 | 11428199 | 6383 | + |
| Chr08 | 11421817 | 11428199 | 6383 | + |
| Chr08 | 11421817 | 11428351 | 6535 | + |
| Chr08 | 11421817 | 11428741 | 6925 | + |
| Chr08 | 11421817 | 11428921 | 7105 | + |
| Chr08 | 11421817 | 11429100 | 7284 | + |
| Chr08 | 11421817 | 11429100 | 7284 | + |
| Chr08 | 11421817 | 11429100 | 7284 | + |
| Chr08 | 11421817 | 11429639 | 7823 | + |
| Chr08 | 11421817 | 11429639 | 7823 | + |
| Chr08 | 11421817 | 11429819 | 8003 | + |
| Chr08 | 11421817 | 11432025 | 10209 | + |
| Chr08 | 11421817 | 11432378 | 10562 | + |
| Chr08 | 11421817 | 11432558 | 10742 | + |
| Chr08 | 11421817 | 11433100 | 11284 | + |
| Chr08 | 11421817 | 11433642 | 11826 | + |

| | | | | |
|---|---|---|---|---|
| Chr08 | 11421817 | 11433642 | 11826 | + |
| Chr08 | 11421817 | 11434167 | 12351 | + |
| Chr08 | 11421817 | 11434693 | 12877 | + |
| Chr08 | 11421817 | 11435414 | 13598 | + |
| Chr08 | 11421817 | 11435592 | 13776 | + |
| Chr08 | 11421817 | 11436134 | 14318 | + |
| Chr08 | 11421817 | 11436134 | 14318 | + |
| Chr08 | 11421817 | 11436134 | 14318 | + |
| Chr08 | 11421817 | 11436851 | 15035 | + |
| Chr08 | 11421817 | 11437030 | 15214 | + |
| Chr08 | 11421817 | 11437210 | 15394 | + |
| Chr08 | 11421817 | 11437210 | 15394 | + |
| Chr08 | 11421817 | 11437389 | 15573 | + |
| Chr08 | 11421817 | 11437569 | 15753 | + |
| Chr08 | 11421817 | 11437749 | 15933 | + |
| Chr08 | 11421817 | 11438472 | 16656 | + |
| Chr08 | 11421817 | 11438833 | 17017 | + |
| Chr08 | 11421817 | 11438833 | 17017 | + |
| Chr08 | 11421817 | 11438833 | 17017 | + |
| Chr08 | 11421817 | 11439319 | 17503 | + |
| Chr08 | 11421817 | 11439680 | 17864 | + |
| Chr08 | 11421817 | 11440401 | 18585 | + |
| Chr08 | 11421817 | 11440583 | 18767 | + |
| Chr08 | 11421817 | 11440943 | 19127 | + |
| Chr08 | 11421817 | 11441124 | 19308 | + |
| Chr08 | 11421817 | 11441124 | 19308 | + |
| Chr08 | 11421817 | 11441124 | 19308 | + |
| Chr08 | 11421817 | 11441473 | 19657 | + |
| Chr08 | 11421817 | 11441473 | 19657 | + |
| Chr08 | 11421817 | 11441835 | 20019 | + |
| Chr08 | 11421817 | 11441835 | 20019 | + |
| Chr08 | 11421817 | 11442197 | 20381 | + |
| Chr08 | 11421817 | 11442197 | 20381 | + |

| | | | | |
|---|---|---|---|---|
| Chr08 | 11421817 | 11442377 | 20561 | + |
| Chr08 | 11421817 | 11442739 | 20923 | + |
| Chr08 | 11421817 | 11443270 | 21454 | + |
| Chr08 | 11421817 | 11443270 | 21454 | + |
| Chr08 | 11421817 | 11443270 | 21454 | + |
| Chr08 | 11421817 | 11443422 | 21606 | + |
| Chr08 | 11421817 | 11443783 | 21967 | + |
| Chr08 | 11421817 | 11444343 | 22527 | + |
| Chr08 | 11421817 | 11444495 | 22679 | + |
| Chr08 | 11421817 | 11444495 | 22679 | + |
| Chr08 | 11421817 | 11444676 | 22860 | + |
| Chr08 | 11421817 | 11444886 | 23070 | + |
| Chr08 | 11421817 | 11445246 | 23430 | + |
| Chr08 | 11421817 | 11445246 | 23430 | + |
| Chr08 | 11421817 | 11445246 | 23430 | + |
| Chr08 | 11421817 | 11445246 | 23430 | + |
| Chr08 | 11421817 | 11445578 | 23762 | + |
| Chr08 | 11421817 | 11445759 | 23943 | + |
| Chr08 | 11421817 | 11445759 | 23943 | + |
| Chr08 | 11421817 | 11445759 | 23943 | + |
| Chr08 | 11421817 | 11445759 | 23943 | + |
| Chr08 | 11421817 | 11445928 | 24112 | + |
| Chr08 | 11421817 | 11445957 | 24141 | + |
| Chr08 | 11421817 | 11445957 | 24141 | + |
| Chr08 | 11421817 | 11446283 | 24467 | + |
| Chr08 | 11421817 | 11446458 | 24642 | + |
| Chr08 | 11421817 | 11446843 | 25027 | + |
| Chr08 | 11421817 | 11446843 | 25027 | + |
| Chr08 | 11421817 | 11447029 | 25213 | + |
| Chr08 | 11421817 | 11447029 | 25213 | + |
| Chr08 | 11421817 | 11447208 | 25392 | + |
| Chr08 | 11421817 | 11447208 | 25392 | + |
| Chr08 | 11421817 | 11447208 | 25392 | + |

| | | | | |
|---|---|---|---|---|
| Chr08 | 11421817 | 11447358 | 25542 | + |
| Chr08 | 11421817 | 11447904 | 26088 | + |
| Chr08 | 11421817 | 11448084 | 26268 | + |
| Chr08 | 11421817 | 11448804 | 26988 | + |
| Chr08 | 11421817 | 11448953 | 27137 | + |
| Chr08 | 11421817 | 11448953 | 27137 | + |
| Chr08 | 11421817 | 11448977 | 27161 | + |
| Chr08 | 11421817 | 11449303 | 27487 | + |
| Chr08 | 11421817 | 11449480 | 27664 | + |
| Chr08 | 11421817 | 11449689 | 27873 | + |
| Chr08 | 11421817 | 11449838 | 28022 | + |
| Chr08 | 11421817 | 11449869 | 28053 | + |
| Chr08 | 11421817 | 11450199 | 28383 | + |
| Chr08 | 11421817 | 11450228 | 28412 | + |
| Chr08 | 11421817 | 11450935 | 29119 | + |
| Chr08 | 11421817 | 11451111 | 29295 | + |
| Chr08 | 11421817 | 11451111 | 29295 | + |
| Chr08 | 11421817 | 11451439 | 29623 | + |
| Chr08 | 11421817 | 11451648 | 29832 | + |
| Chr08 | 11421817 | 11451828 | 30012 | + |
| Chr08 | 11421817 | 11452188 | 30372 | + |
| Chr08 | 11421977 | 11429639 | 7663 | + |
| Chr08 | 11422493 | 11430735 | 8243 | + |
| Chr08 | 11422493 | 11432378 | 9886 | + |
| Chr08 | 11422493 | 11435414 | 12922 | + |
| Chr08 | 11422493 | 11440943 | 18451 | + |
| Chr08 | 11422493 | 11441473 | 18981 | + |
| Chr08 | 11422493 | 11445040 | 22548 | + |
| Chr08 | 11422493 | 11445426 | 22934 | + |
| Chr08 | 11422997 | 11437210 | 14214 | + |
| Chr08 | 11422997 | 11444886 | 21890 | + |
| Chr08 | 11423007 | 11425357 | 2351 | + |
| Chr08 | 11423007 | 11425533 | 2527 | + |

| | | | | |
|---|---|---|---|---|
| Chr08 | 11423007 | 11426051 | 3045 | + |
| Chr08 | 11423007 | 11428532 | 5526 | + |
| Chr08 | 11423007 | 11429459 | 6453 | + |
| Chr08 | 11423007 | 11429459 | 6453 | + |
| Chr08 | 11423007 | 11432378 | 9372 | + |
| Chr08 | 11423007 | 11432919 | 9913 | + |
| Chr08 | 11423007 | 11433281 | 10275 | + |
| Chr08 | 11423007 | 11433642 | 10636 | + |
| Chr08 | 11423007 | 11434693 | 11687 | + |
| Chr08 | 11423007 | 11435234 | 12228 | + |
| Chr08 | 11423007 | 11435592 | 12586 | + |
| Chr08 | 11423007 | 11435592 | 12586 | + |
| Chr08 | 11423007 | 11437569 | 14563 | + |
| Chr08 | 11423007 | 11437569 | 14563 | + |
| Chr08 | 11423007 | 11438472 | 15466 | + |
| Chr08 | 11423007 | 11438833 | 15827 | + |
| Chr08 | 11423007 | 11440220 | 17214 | + |
| Chr08 | 11423007 | 11440943 | 17937 | + |
| Chr08 | 11423007 | 11440943 | 17937 | + |
| Chr08 | 11423007 | 11441124 | 18118 | + |
| Chr08 | 11423007 | 11441305 | 18299 | + |
| Chr08 | 11423007 | 11441473 | 18467 | + |
| Chr08 | 11423007 | 11441835 | 18829 | + |
| Chr08 | 11423007 | 11442016 | 19010 | + |
| Chr08 | 11423007 | 11442197 | 19191 | + |
| Chr08 | 11423007 | 11443270 | 20264 | + |
| Chr08 | 11423007 | 11444343 | 21337 | + |
| Chr08 | 11423007 | 11444495 | 21489 | + |
| Chr08 | 11423007 | 11444886 | 21880 | + |
| Chr08 | 11423007 | 11445578 | 22572 | + |
| Chr08 | 11423007 | 11445759 | 22753 | + |
| Chr08 | 11423007 | 11445759 | 22753 | + |
| Chr08 | 11423007 | 11445928 | 22922 | + |

| | | | | |
|---|---|---|---|---|
| Chr08 | 11423007 | 11445957 | 22951 | + |
| Chr08 | 11423007 | 11445957 | 22951 | + |
| Chr08 | 11423007 | 11446843 | 23837 | + |
| Chr08 | 11423007 | 11446843 | 23837 | + |
| Chr08 | 11423007 | 11447566 | 24560 | + |
| Chr08 | 11423007 | 11447904 | 24898 | + |
| Chr08 | 11423007 | 11447904 | 24898 | + |
| Chr08 | 11423007 | 11448601 | 25595 | + |
| Chr08 | 11423007 | 11448804 | 25798 | + |
| Chr08 | 11423007 | 11448953 | 25947 | + |
| Chr08 | 11423007 | 11448977 | 25971 | + |
| Chr08 | 11423007 | 11449303 | 26297 | + |
| Chr08 | 11423007 | 11449689 | 26683 | + |
| Chr08 | 11423007 | 11449838 | 26832 | + |
| Chr08 | 11423007 | 11449869 | 26863 | + |
| Chr08 | 11423007 | 11449869 | 26863 | + |
| Chr08 | 11423007 | 11450020 | 27014 | + |
| Chr08 | 11423007 | 11450228 | 27222 | + |
| Chr08 | 11423007 | 11450935 | 27929 | + |
| Chr08 | 11423007 | 11452188 | 29182 | + |
| Chr08 | 11423007 | 11453944 | 30938 | + |
| Chr08 | 11423022 | 11447722 | 24701 | + |
| Chr08 | 11423022 | 11447722 | 24701 | + |
| Chr08 | 11423362 | 11426051 | 2690 | + |
| Chr08 | 11424202 | 11449303 | 25102 | + |
| Chr08 | 11424202 | 11450228 | 26027 | + |
| Chr08 | 11424202 | 11453944 | 29743 | + |
| Chr08 | 11424530 | 11428741 | 4212 | + |
| Chr08 | 11424530 | 11453944 | 29415 | + |
| Chr08 | 11424545 | 11447722 | 23178 | + |
| Chr08 | 11425923 | 11453944 | 28022 | + |
| Chr08 | 11425923 | 11453944 | 28022 | + |
| Chr08 | 11425923 | 11453944 | 28022 | + |

| | | | | |
|---|---|---|---|---|
| Chr08 | 11425923 | 11453944 | 28022 | + |
| Chr08 | 11426279 | 11440734 | 14456 | + |
| Chr08 | 11430230 | 11453951 | 23722 | + |
| Chr08 | 11436720 | 11456151 | 19432 | + |
| Chr08 | 11437980 | 11454905 | 16926 | + |
| Chr08 | 11446895 | 11469712 | 22818 | + |
| Chr08 | 11449201 | 11473605 | 24405 | + |
| Chr08 | 11453124 | 11469712 | 16589 | + |
| Chr08 | 11584429 | 11598013 | 13585 | - |
| Chr08 | 12587856 | 12593500 | 5645 | - |
| Chr08 | 12724061 | 12729641 | 5581 | - |
| Chr08 | 12883127 | 12887820 | 4694 | - |
| Chr08 | 13275731 | 13286329 | 10599 | + |
| Chr08 | 13837067 | 13843203 | 6137 | + |
| Chr08 | 14200941 | 14206860 | 5920 | + |
| Chr08 | 14280685 | 14289323 | 8639 | - |
| Chr08 | 14808373 | 14813417 | 5045 | + |
| Chr08 | 14817839 | 14823327 | 5489 | - |
| Chr08 | 15055069 | 15060215 | 5147 | - |
| Chr08 | 15748342 | 15755476 | 7135 | + |
| Chr08 | 16112079 | 16132155 | 20077 | + |
| Chr08 | 16248000 | 16250457 | 2458 | - |
| Chr08 | 16296095 | 16301706 | 5612 | - |
| Chr08 | 16341229 | 16346472 | 5244 | + |
| Chr08 | 16357106 | 16362260 | 5155 | + |
| Chr08 | 16389644 | 16395047 | 5404 | + |
| Chr08 | 16402116 | 16416507 | 14392 | - |
| Chr08 | 16402116 | 16430039 | 27924 | - |
| Chr08 | 16588638 | 16597880 | 9243 | + |
| Chr08 | 16589323 | 16593779 | 4457 | + |
| Chr08 | 16603758 | 16613716 | 9959 | - |
| Chr08 | 16618802 | 16627739 | 8938 | + |
| Chr08 | 16672959 | 16678006 | 5048 | + |

| | | | | |
|---|---|---|---|---|
| Chr08 | 16707025 | 16712505 | 5481 | + |
| Chr08 | 17313863 | 17325958 | 12096 | - |
| Chr08 | 17744487 | 17750597 | 6111 | + |
| Chr08 | 18218201 | 18223436 | 5236 | + |
| Chr08 | 18260196 | 18267299 | 7104 | - |
| Chr08 | 18260196 | 18279979 | 19784 | - |
| Chr08 | 18621860 | 18626827 | 4968 | + |
| Chr08 | 18627933 | 18637460 | 9528 | - |
| Chr09 | 328370 | 333506 | 5137 | - |
| Chr09 | 635575 | 650296 | 14722 | - |
| Chr09 | 832733 | 838103 | 5371 | - |
| Chr09 | 1248516 | 1253614 | 5099 | - |
| Chr09 | 1307145 | 1312649 | 5505 | + |
| Chr09 | 1656467 | 1661715 | 5249 | + |
| Chr09 | 1950398 | 1955886 | 5489 | - |
| Chr09 | 2449775 | 2454984 | 5210 | + |
| Chr09 | 2483179 | 2506587 | 23409 | - |
| Chr09 | 2513174 | 2536480 | 23307 | - |
| Chr09 | 2846712 | 2852215 | 5504 | - |
| Chr09 | 2869235 | 2874633 | 5399 | + |
| Chr09 | 3162323 | 3175588 | 13266 | - |
| Chr09 | 3281955 | 3287366 | 5412 | + |
| Chr09 | 3555988 | 3561052 | 5065 | + |
| Chr09 | 3698224 | 3703635 | 5412 | + |
| Chr09 | 4139423 | 4145067 | 5645 | - |
| Chr09 | 4182320 | 4186019 | 3700 | + |
| Chr09 | 4984305 | 4993407 | 9103 | + |
| Chr09 | 5002746 | 5007979 | 5234 | - |
| Chr09 | 5177581 | 5180886 | 3306 | + |
| Chr09 | 5189730 | 5197443 | 7714 | + |
| Chr09 | 5555046 | 5562147 | 7102 | - |
| Chr09 | 5594120 | 5599667 | 5548 | - |
| Chr09 | 6007203 | 6012473 | 5271 | - |

| | | | | |
|---|---|---|---|---|
| Chr09 | 6073155 | 6078535 | 5381 | + |
| Chr09 | 6257203 | 6262243 | 5041 | - |
| Chr09 | 6567676 | 6578666 | 10991 | + |
| Chr09 | 6749324 | 6754385 | 5062 | - |
| Chr09 | 6784403 | 6796653 | 12251 | + |
| Chr09 | 6786725 | 6794309 | 7585 | + |
| Chr09 | 7023206 | 7044443 | 21238 | - |
| Chr09 | 7401403 | 7406041 | 4639 | + |
| Chr09 | 7505378 | 7512215 | 6838 | - |
| Chr09 | 8083090 | 8089203 | 6114 | + |
| Chr09 | 8308582 | 8310987 | 2406 | + |
| Chr09 | 8478486 | 8485290 | 6805 | + |
| Chr09 | 8584538 | 8589936 | 5399 | - |
| Chr09 | 8584538 | 8609425 | 24888 | - |
| Chr09 | 8722748 | 8729023 | 6276 | - |
| Chr09 | 8810980 | 8814404 | 3425 | + |
| Chr09 | 8875549 | 8877543 | 1995 | + |
| Chr09 | 9092899 | 9101741 | 8843 | - |
| Chr09 | 9119109 | 9125996 | 6888 | - |
| Chr09 | 9218231 | 9230241 | 12011 | + |
| Chr09 | 9410287 | 9415412 | 5126 | + |
| Chr09 | 9419745 | 9425859 | 6115 | + |
| Chr09 | 9487427 | 9492302 | 4876 | - |
| Chr09 | 9742237 | 9750214 | 7978 | - |
| Chr09 | 9798000 | 9807701 | 9702 | - |
| Chr09 | 9962907 | 9967957 | 5051 | + |
| Chr09 | 9976791 | 9981824 | 5034 | - |
| Chr09 | 10178941 | 10184367 | 5427 | + |
| Chr09 | 10243249 | 10253876 | 10628 | - |
| Chr09 | 10380400 | 10385614 | 5215 | - |
| Chr09 | 10380400 | 10389688 | 9289 | - |
| Chr09 | 11373550 | 11377547 | 3998 | - |
| Chr09 | 11391285 | 11396668 | 5384 | - |

| | | | | |
|---|---|---|---|---|
| Chr09 | 11576101 | 11581823 | 5723 | + |
| Chr09 | 11689756 | 11691907 | 2152 | + |
| Chr09 | 11731917 | 11742708 | 10792 | - |
| Chr09 | 11736368 | 11741794 | 5427 | - |
| Chr09 | 11889507 | 11899244 | 9738 | + |
| Chr09 | 12082636 | 12088744 | 6109 | + |
| Chr09 | 12414104 | 12416048 | 1945 | + |
| Chr09 | 12523868 | 12527897 | 4030 | - |
| Chr09 | 12523868 | 12529335 | 5468 | - |
| Chr09 | 12523868 | 12530597 | 6730 | - |
| Chr09 | 12523868 | 12532018 | 8151 | - |
| Chr09 | 12523868 | 12532920 | 9053 | - |
| Chr09 | 12523868 | 12533644 | 9777 | - |
| Chr09 | 12523868 | 12540832 | 16965 | - |
| Chr09 | 12523868 | 12556816 | 32949 | - |
| Chr09 | 12523868 | 12556816 | 32949 | - |
| Chr09 | 12527613 | 12561722 | 34110 | - |
| Chr09 | 12530675 | 12561722 | 31048 | - |
| Chr09 | 12531367 | 12561722 | 30356 | - |
| Chr09 | 12532458 | 12561722 | 29265 | - |
| Chr09 | 12532638 | 12561722 | 29085 | - |
| Chr09 | 12533722 | 12561722 | 28001 | - |
| Chr09 | 12533901 | 12561722 | 27822 | - |
| Chr09 | 12534081 | 12561722 | 27642 | - |
| Chr09 | 12534980 | 12561722 | 26743 | - |
| Chr09 | 12535132 | 12561722 | 26591 | - |
| Chr09 | 12535882 | 12561722 | 25841 | - |
| Chr09 | 12536061 | 12561722 | 25662 | - |
| Chr09 | 12548023 | 12561722 | 13700 | - |
| Chr09 | 12548924 | 12556816 | 7893 | - |
| Chr09 | 12548924 | 12561722 | 12799 | - |
| Chr09 | 12548924 | 12561722 | 12799 | - |
| Chr09 | 12548924 | 12561722 | 12799 | - |

| | | | | |
|---|---|---|---|---|
| Chr09 | 12548924 | 12561722 | 12799 | - |
| Chr09 | 12549066 | 12561722 | 12657 | - |
| Chr09 | 12549573 | 12556816 | 7244 | - |
| Chr09 | 12549573 | 12561722 | 12150 | - |
| Chr09 | 12549573 | 12561722 | 12150 | - |
| Chr09 | 12549573 | 12561722 | 12150 | - |
| Chr09 | 12549573 | 12561722 | 12150 | - |
| Chr09 | 12549573 | 12561722 | 12150 | - |
| Chr09 | 12550806 | 12561722 | 10917 | - |
| Chr09 | 12553852 | 12561722 | 7871 | - |
| Chr09 | 12554747 | 12561722 | 6976 | - |
| Chr09 | 12556864 | 12561722 | 4859 | - |
| Chr09 | 12558601 | 12561722 | 3122 | - |
| Chr09 | 12558601 | 12561722 | 3122 | + |
| Chr09 | 12613036 | 12633094 | 20059 | + |
| Chr09 | 12614388 | 12631975 | 17588 | - |
| Chr09 | 13593821 | 13605961 | 12141 | - |
| Chr09 | 13718021 | 13730769 | 12749 | + |
| Chr09 | 14225214 | 14235228 | 10015 | + |
| Chr09 | 14541622 | 14553577 | 11956 | - |
| Chr09 | 14697960 | 14705506 | 7547 | - |
| Chr09 | 14806709 | 14816519 | 9811 | + |
| Chr09 | 14807717 | 14815531 | 7815 | + |
| Chr09 | 15008921 | 15012963 | 4043 | + |
| Chr09 | 15037489 | 15042783 | 5295 | - |
| Chr09 | 15296083 | 15297696 | 1614 | - |
| Chr09 | 15311296 | 15316747 | 5452 | - |
| Chr09 | 15714549 | 15720051 | 5503 | + |
| Chr09 | 16001227 | 16008034 | 6808 | - |
| Chr09 | 16238246 | 16255452 | 17207 | - |
| Chr09 | 16262668 | 16264635 | 1968 | - |
| Chr09 | 16373592 | 16375390 | 1799 | + |
| Chr09 | 16505957 | 16507754 | 1798 | + |

| | | | | |
|---|---|---|---|---|
| Chr09 | 16643084 | 16647586 | 4503 | - |
| Chr09 | 17104700 | 17111064 | 6365 | + |
| Chr09 | 17415310 | 17420496 | 5187 | + |
| Chr09 | 17888921 | 17901513 | 12593 | + |
| Chr09 | 18429455 | 18450353 | 20899 | - |
| Chr09 | 18501022 | 18505643 | 4622 | - |
| Chr09 | 18559823 | 18566789 | 6967 | + |
| Chr09 | 18898380 | 18908870 | 10491 | + |
| Chr09 | 19156620 | 19175287 | 18668 | - |
| Chr09 | 19162805 | 19184834 | 22030 | + |
| Chr09 | 19220063 | 19225083 | 5021 | + |
| Chr09 | 19493383 | 19498405 | 5023 | + |
| Chr09 | 19498631 | 19504495 | 5865 | - |
| Chr09 | 20126317 | 20151208 | 24892 | + |
| Chr09 | 20201158 | 20214439 | 13282 | + |
| Chr09 | 20380108 | 20386831 | 6724 | + |
| Chr09 | 20750300 | 20773716 | 23417 | + |
| Chr09 | 20750316 | 20765163 | 14848 | + |
| Chr09 | 21033762 | 21037344 | 3583 | - |
| Chr09 | 21083014 | 21088430 | 5417 | - |
| Chr09 | 21283132 | 21288483 | 5352 | + |
| Chr09 | 21354018 | 21355930 | 1913 | + |
| Chr09 | 21363878 | 21373836 | 9959 | - |
| Chr09 | 21428642 | 21434422 | 5781 | + |
| Chr09 | 21668212 | 21673638 | 5427 | - |
| Chr09 | 21742019 | 21748025 | 6007 | - |
| Chr09 | 21748189 | 21759012 | 10824 | - |
| Chr09 | 21809706 | 21821166 | 11461 | + |
| Chr09 | 21893465 | 21898427 | 4963 | + |
| Chr09 | 22006556 | 22033103 | 26548 | + |
| Chr09 | 22011727 | 22017340 | 5614 | + |
| Chr09 | 22285737 | 22291068 | 5332 | - |
| Chr09 | 22349818 | 22355432 | 5615 | - |

| | | | | |
|---|---|---|---|---|
| Chr09 | 22439143 | 22444817 | 5675 | - |
| Chr09 | 23023332 | 23028360 | 5029 | - |
| Chr09 | 23577117 | 23581914 | 4798 | + |
| Chr09 | 24002887 | 24008390 | 5504 | + |
| Chr09 | 24195987 | 24201494 | 5508 | - |
| Chr09 | 24317586 | 24328195 | 10610 | - |
| Chr09 | 24317586 | 24328208 | 10623 | - |
| Chr09 | 24361631 | 24367076 | 5446 | + |
| Chr09 | 24500789 | 24510305 | 9517 | - |
| Chr09 | 24530593 | 24536095 | 5503 | + |
| Chr09 | 24587522 | 24592767 | 5246 | - |
| Chr09 | 24735149 | 24741140 | 5992 | + |
| Chr09 | 24885000 | 24892069 | 7070 | + |
| Chr09 | 24899268 | 24908535 | 9268 | + |
| Chr09 | 25189715 | 25198143 | 8429 | + |
| Chr09 | 25309188 | 25315592 | 6405 | + |
| Chr09 | 25358501 | 25372985 | 14485 | + |
| Chr09 | 25424691 | 25448462 | 23772 | - |
| Chr09 | 25873585 | 25878795 | 5211 | + |
| Chr09 | 26057849 | 26062724 | 4876 | - |
| Chr09 | 26057849 | 26079136 | 21288 | - |
| Chr09 | 26337595 | 26346645 | 9051 | + |
| Chr09 | 26432967 | 26440031 | 7065 | - |
| Chr09 | 26474783 | 26479414 | 4632 | + |
| Chr09 | 26587385 | 26592771 | 5387 | - |
| Chr09 | 26625094 | 26632811 | 7718 | + |
| Chr09 | 26711883 | 26716288 | 4406 | + |
| Chr09 | 26718120 | 26724525 | 6406 | - |
| Chr09 | 26719271 | 26724525 | 5255 | - |
| Chr09 | 27295136 | 27300069 | 4934 | + |
| Chr09 | 27407195 | 27413403 | 6209 | - |
| Chr09 | 28302303 | 28307308 | 5006 | + |
| Chr09 | 28328087 | 28337078 | 8992 | + |

| | | | | |
|---|---|---|---|---|
| Chr09 | 28411726 | 28416950 | 5225 | + |
| Chr09 | 28425229 | 28430325 | 5097 | - |
| Chr09 | 28868903 | 28873959 | 5057 | + |
| Chr09 | 28879229 | 28884846 | 5618 | + |
| Chr09 | 29034290 | 29040445 | 6156 | - |
| Chr09 | 29110295 | 29117762 | 7468 | - |
| Chr09 | 29376926 | 29383524 | 6599 | + |
| Chr09 | 29572415 | 29595246 | 22832 | - |
| Chr09 | 29589185 | 29594594 | 5410 | - |
| Chr09 | 30183093 | 30189359 | 6267 | - |
| Chr09 | 30183093 | 30189359 | 6267 | - |

**Supplementary Table 4: Coordinate of the long trasposable elements (LTR) on the alternative haplotype**

| Chromosome | Start | End | Length | Strand |
|---|---|---|---|---|
| Chr01 | 18955 | 29568 | 10614 | - |
| Chr01 | 476764 | 481740 | 4977 | + |
| Chr01 | 1014478 | 1019840 | 5363 | - |
| Chr01 | 1252015 | 1257263 | 5249 | + |
| Chr01 | 1352657 | 1357261 | 4605 | + |
| Chr01 | 1373177 | 1379528 | 6352 | - |
| Chr01 | 1667617 | 1669390 | 1774 | + |
| Chr01 | 1902022 | 1906756 | 4735 | + |
| Chr01 | 2041774 | 2048848 | 7075 | + |
| Chr01 | 2070655 | 2080298 | 9644 | - |
| Chr01 | 2656906 | 2675115 | 18210 | + |
| Chr01 | 2675260 | 2680183 | 4924 | + |
| Chr01 | 3056713 | 3061747 | 5035 | - |
| Chr01 | 3075226 | 3080352 | 5127 | + |
| Chr01 | 3461871 | 3472654 | 10784 | - |
| Chr01 | 4288467 | 4296717 | 8251 | + |
| Chr01 | 4994063 | 4999698 | 5636 | + |
| Chr01 | 6886811 | 6898871 | 12061 | + |
| Chr01 | 6937580 | 6944866 | 7287 | - |
| Chr01 | 7563569 | 7570805 | 7237 | + |
| Chr01 | 8848371 | 8856371 | 8001 | - |
| Chr01 | 9048824 | 9054255 | 5432 | + |
| Chr01 | 9079644 | 9091516 | 11873 | - |
| Chr01 | 9464444 | 9469992 | 5549 | - |
| Chr01 | 9738729 | 9743309 | 4581 | + |
| Chr01 | 9753340 | 9758697 | 5358 | - |
| Chr01 | 9776206 | 9781606 | 5401 | - |
| Chr01 | 9826029 | 9831309 | 5281 | + |

| | | | | |
|---|---|---|---|---|
| Chr01 | 9891838 | 9897167 | 5330 | + |
| Chr01 | 9929613 | 9939954 | 10342 | - |
| Chr01 | 9944361 | 9949370 | 5010 | + |
| Chr01 | 9981013 | 9990668 | 9656 | + |
| Chr01 | 10167109 | 10172514 | 5406 | + |
| Chr01 | 10200681 | 10211099 | 10419 | - |
| Chr01 | 10346807 | 10353437 | 6631 | + |
| Chr01 | 10724213 | 10729288 | 5076 | - |
| Chr01 | 10724350 | 10728469 | 4120 | - |
| Chr01 | 10957001 | 10962644 | 5644 | + |
| Chr01 | 12561548 | 12567910 | 6363 | - |
| Chr01 | 12807898 | 12820127 | 12230 | - |
| Chr01 | 12825518 | 12835167 | 9650 | - |
| Chr01 | 12918759 | 12924137 | 5379 | + |
| Chr01 | 12938778 | 12942575 | 3798 | - |
| Chr01 | 13093848 | 13117249 | 23402 | - |
| Chr01 | 13192104 | 13204379 | 12276 | + |
| Chr01 | 13582014 | 13587151 | 5138 | + |
| Chr01 | 13611885 | 13614972 | 3088 | + |
| Chr01 | 13941155 | 13948005 | 6851 | + |
| Chr01 | 14355429 | 14366482 | 11054 | - |
| Chr01 | 14398868 | 14406452 | 7585 | - |
| Chr01 | 14456491 | 14461154 | 4664 | - |
| Chr01 | 14739523 | 14754829 | 15307 | + |
| Chr01 | 15037772 | 15048703 | 10932 | - |
| Chr01 | 15252982 | 15275372 | 22391 | + |
| Chr01 | 15348611 | 15355081 | 6471 | - |
| Chr01 | 15418756 | 15423810 | 5055 | - |
| Chr01 | 15524963 | 15530226 | 5264 | + |
| Chr01 | 15537054 | 15543241 | 6188 | - |
| Chr01 | 15711821 | 15717149 | 5329 | + |
| Chr01 | 15803328 | 15815775 | 12448 | + |

| | | | | |
|---|---|---|---|---|
| Chr01 | 15987607 | 15995467 | 7861 | + |
| Chr01 | 16037722 | 16042572 | 4851 | + |
| Chr01 | 16112269 | 16117325 | 5057 | + |
| Chr01 | 16164802 | 16170056 | 5255 | - |
| Chr01 | 16223409 | 16230085 | 6677 | + |
| Chr01 | 16280694 | 16282630 | 1937 | + |
| Chr01 | 16386717 | 16393732 | 7016 | - |
| Chr01 | 16639897 | 16644933 | 5037 | + |
| Chr01 | 16862192 | 16867213 | 5022 | + |
| Chr01 | 16963678 | 16968688 | 5011 | + |
| Chr01 | 17098505 | 17104131 | 5627 | + |
| Chr01 | 17209066 | 17213881 | 4816 | - |
| Chr01 | 17361852 | 17372717 | 10866 | + |
| Chr01 | 17494686 | 17501985 | 7300 | - |
| Chr01 | 17608029 | 17625392 | 17364 | + |
| Chr01 | 17846382 | 17872037 | 25656 | - |
| Chr01 | 18282356 | 18289466 | 7111 | - |
| Chr01 | 18429319 | 18439647 | 10329 | + |
| Chr01 | 18578336 | 18606420 | 28085 | + |
| Chr01 | 18795907 | 18810834 | 14928 | - |
| Chr01 | 18936258 | 18941315 | 5058 | - |
| Chr01 | 18949351 | 18960047 | 10697 | + |
| Chr01 | 19244936 | 19256210 | 11275 | - |
| Chr01 | 19275139 | 19277089 | 1951 | + |
| Chr01 | 19548256 | 19554848 | 6593 | + |
| Chr01 | 19639300 | 19641264 | 1965 | + |
| Chr01 | 19838733 | 19843400 | 4668 | - |
| Chr01 | 20015662 | 20026284 | 10623 | - |
| Chr01 | 20603855 | 20614533 | 10679 | - |
| Chr01 | 20621366 | 20625025 | 3660 | + |
| Chr01 | 20962245 | 20967557 | 5313 | - |
| Chr01 | 21078253 | 21088889 | 10637 | - |

| | | | | |
|---|---|---|---|---|
| Chr01 | 21260244 | 21266109 | 5866 | + |
| Chr01 | 21418194 | 21441203 | 23010 | + |
| Chr01 | 21586432 | 21591887 | 5456 | - |
| Chr01 | 21640848 | 21650798 | 9951 | - |
| Chr01 | 21713795 | 21719239 | 5445 | - |
| Chr01 | 21955983 | 21962920 | 6938 | + |
| Chr01 | 22292236 | 22315640 | 23405 | + |
| Chr01 | 22295492 | 22309764 | 14273 | - |
| Chr01 | 22295746 | 22310288 | 14543 | + |
| Chr01 | 22296287 | 22306754 | 10468 | + |
| Chr01 | 22296287 | 22314700 | 18414 | + |
| Chr01 | 22296690 | 22307095 | 10406 | + |
| Chr01 | 22296690 | 22314981 | 18292 | + |
| Chr01 | 22296882 | 22323114 | 26233 | + |
| Chr01 | 22306383 | 22315640 | 9258 | + |
| Chr01 | 22325294 | 22331170 | 5877 | + |
| Chr01 | 22325294 | 22338785 | 13492 | + |
| Chr01 | 22332914 | 22338785 | 5872 | + |
| Chr01 | 22384395 | 22387869 | 3475 | + |
| Chr01 | 22488795 | 22495676 | 6882 | + |
| Chr01 | 22630975 | 22643699 | 12725 | + |
| Chr01 | 22681234 | 22689622 | 8389 | - |
| Chr01 | 22693373 | 22711098 | 17726 | - |
| Chr01 | 22819462 | 22832091 | 12630 | + |
| Chr01 | 22851770 | 22856654 | 4885 | + |
| Chr01 | 23176192 | 23181692 | 5501 | + |
| Chr01 | 23450852 | 23456789 | 5938 | + |
| Chr01 | 23490167 | 23499855 | 9689 | + |
| Chr01 | 23754205 | 23764380 | 10176 | + |
| Chr01 | 23780926 | 23786415 | 5490 | + |
| Chr01 | 23878582 | 23890156 | 11575 | - |
| Chr01 | 24395942 | 24400948 | 5007 | + |

| | | | | |
|---|---|---|---|---|
| Chr01 | 24477873 | 24481221 | 3349 | + |
| Chr01 | 24515985 | 24522958 | 6974 | - |
| Chr01 | 24655057 | 24659985 | 4929 | + |
| Chr01 | 24787751 | 24797745 | 9995 | + |
| Chr01 | 24909824 | 24924063 | 14240 | - |
| Chr01 | 25139911 | 25144668 | 4758 | + |
| Chr01 | 25147771 | 25154698 | 6928 | - |
| Chr01 | 25161185 | 25166203 | 5019 | - |
| Chr01 | 25173501 | 25181314 | 7814 | + |
| Chr01 | 25273699 | 25284003 | 10305 | + |
| Chr01 | 25672089 | 25675035 | 2947 | + |
| Chr01 | 25680160 | 25685008 | 4849 | + |
| Chr01 | 26006770 | 26018958 | 12189 | + |
| Chr01 | 26110429 | 26115715 | 5287 | + |
| Chr01 | 26125566 | 26133312 | 7747 | + |
| Chr01 | 26386677 | 26391949 | 5273 | + |
| Chr01 | 26458289 | 26463268 | 4980 | - |
| Chr01 | 26758252 | 26765374 | 7123 | - |
| Chr01 | 26799164 | 26804575 | 5412 | + |
| Chr01 | 26840570 | 26850409 | 9840 | + |
| Chr01 | 26862114 | 26888617 | 26504 | - |
| Chr01 | 27111757 | 27127097 | 15341 | - |
| Chr01 | 27112704 | 27128071 | 15368 | - |
| Chr01 | 27657970 | 27663493 | 5524 | + |
| Chr01 | 27756565 | 27763222 | 6658 | - |
| Chr01 | 27782489 | 27791851 | 9363 | - |
| Chr01 | 27871851 | 27884158 | 12308 | + |
| Chr01 | 27898123 | 27903939 | 5817 | - |
| Chr01 | 27963069 | 27973400 | 10332 | - |
| Chr01 | 28041155 | 28050904 | 9750 | - |
| Chr01 | 28468522 | 28475063 | 6542 | + |
| Chr01 | 28674526 | 28679185 | 4660 | + |

| | | | | |
|---|---|---|---|---|
| Chr01 | 28960110 | 28964888 | 4779 | - |
| Chr01 | 29005367 | 29018952 | 13586 | + |
| Chr01 | 29005367 | 29019292 | 13926 | + |
| Chr01 | 29005367 | 29020906 | 15540 | + |
| Chr01 | 29005367 | 29020906 | 15540 | + |
| Chr01 | 29005371 | 29018113 | 12743 | + |
| Chr01 | 29005421 | 29018113 | 12693 | + |
| Chr01 | 29005421 | 29018952 | 13532 | + |
| Chr01 | 29005421 | 29019292 | 13872 | + |
| Chr01 | 29090443 | 29095890 | 5448 | + |
| Chr01 | 29187997 | 29189341 | 1345 | + |
| Chr01 | 29187997 | 29189708 | 1712 | - |
| Chr01 | 29187997 | 29189708 | 1712 | + |
| Chr01 | 29187997 | 29190108 | 2112 | - |
| Chr01 | 29187997 | 29190108 | 2112 | + |
| Chr01 | 29187997 | 29190848 | 2852 | - |
| Chr01 | 29187997 | 29190848 | 2852 | + |
| Chr01 | 29187997 | 29191216 | 3220 | - |
| Chr01 | 29187997 | 29191216 | 3220 | + |
| Chr01 | 29187997 | 29191628 | 3632 | - |
| Chr01 | 29187997 | 29191628 | 3632 | + |
| Chr01 | 29187997 | 29192361 | 4365 | + |
| Chr01 | 29187997 | 29192361 | 4365 | + |
| Chr01 | 29187997 | 29192756 | 4760 | + |
| Chr01 | 29187997 | 29192756 | 4760 | + |
| Chr01 | 29187997 | 29192756 | 4760 | + |
| Chr01 | 29187997 | 29192756 | 4760 | + |
| Chr01 | 29187997 | 29193522 | 5526 | + |
| Chr01 | 29187997 | 29193522 | 5526 | + |
| Chr01 | 29187997 | 29193920 | 5924 | + |
| Chr01 | 29190704 | 29192361 | 1658 | + |
| Chr01 | 29190704 | 29192756 | 2053 | + |

| | | | | |
|---|---|---|---|---|
| Chr01 | 29190704 | 29193522 | 2819 | + |
| Chr01 | 29190704 | 29193920 | 3217 | + |
| Chr01 | 29191072 | 29192361 | 1290 | + |
| Chr01 | 29191072 | 29193522 | 2451 | + |
| Chr01 | 29191072 | 29193920 | 2849 | + |
| Chr01 | 29247308 | 29265942 | 18635 | - |
| Chr01 | 29370929 | 29376244 | 5316 | - |
| Chr01 | 29424914 | 29431864 | 6951 | + |
| Chr01 | 29431980 | 29436921 | 4942 | + |
| Chr01 | 29537488 | 29542681 | 5194 | - |
| Chr01 | 29544881 | 29549776 | 4896 | + |
| Chr01 | 29572355 | 29577859 | 5505 | + |
| Chr01 | 29607978 | 29613444 | 5467 | - |
| Chr01 | 29671502 | 29681370 | 9869 | + |
| Chr01 | 29930361 | 29935177 | 4817 | - |
| Chr01 | 30052433 | 30056518 | 4086 | + |
| Chr01 | 30053736 | 30056518 | 2783 | + |
| Chr01 | 30286751 | 30292134 | 5384 | + |
| Chr02 | 165786 | 171302 | 5517 | - |
| Chr02 | 1022422 | 1035770 | 13349 | + |
| Chr02 | 1261315 | 1267592 | 6278 | + |
| Chr02 | 1484204 | 1495031 | 10828 | - |
| Chr02 | 1548326 | 1553547 | 5222 | + |
| Chr02 | 1626593 | 1631770 | 5178 | + |
| Chr02 | 1855537 | 1860966 | 5430 | + |
| Chr02 | 1956375 | 1983544 | 27170 | - |
| Chr02 | 1961867 | 1976060 | 14194 | - |
| Chr02 | 1965478 | 1972126 | 6649 | - |
| Chr02 | 2024268 | 2028758 | 4491 | + |
| Chr02 | 2092470 | 2098821 | 6352 | - |
| Chr02 | 2094509 | 2098711 | 4203 | - |
| Chr02 | 2095379 | 2098821 | 3443 | - |

| | | | | |
|---|---|---|---|---|
| Chr02 | 2146297 | 2158065 | 11769 | - |
| Chr02 | 2233001 | 2238612 | 5612 | - |
| Chr02 | 2355733 | 2366003 | 10271 | + |
| Chr02 | 2378452 | 2403241 | 24790 | + |
| Chr02 | 3463794 | 3469331 | 5538 | + |
| Chr02 | 3539405 | 3544669 | 5265 | + |
| Chr02 | 3657442 | 3662492 | 5051 | + |
| Chr02 | 3692375 | 3697303 | 4929 | + |
| Chr02 | 4749007 | 4765040 | 16034 | - |
| Chr02 | 4942495 | 4949298 | 6804 | + |
| Chr02 | 5070357 | 5077433 | 7077 | - |
| Chr02 | 5927444 | 5932759 | 5316 | - |
| Chr02 | 5944751 | 5952022 | 7272 | + |
| Chr02 | 6242147 | 6247082 | 4936 | + |
| Chr02 | 6259197 | 6271179 | 11983 | + |
| Chr02 | 6783873 | 6793875 | 10003 | - |
| Chr02 | 6825448 | 6846137 | 20690 | + |
| Chr02 | 6826544 | 6847940 | 21397 | - |
| Chr02 | 6855726 | 6861255 | 5530 | + |
| Chr02 | 6901765 | 6909017 | 7253 | + |
| Chr02 | 6928353 | 6933393 | 5041 | + |
| Chr02 | 6974664 | 6998745 | 24082 | - |
| Chr02 | 8014111 | 8020845 | 6735 | - |
| Chr02 | 8152442 | 8162345 | 9904 | - |
| Chr02 | 8477208 | 8483131 | 5924 | - |
| Chr02 | 8553928 | 8560560 | 6633 | + |
| Chr02 | 8597772 | 8607821 | 10050 | - |
| Chr02 | 9173205 | 9185496 | 12292 | + |
| Chr02 | 9497462 | 9516914 | 19453 | + |
| Chr02 | 9504213 | 9510344 | 6132 | - |
| Chr02 | 9557715 | 9562853 | 5139 | + |
| Chr02 | 9688813 | 9693239 | 4427 | + |

| | | | | |
|---|---|---|---|---|
| Chr02 | 9848656 | 9850656 | 2001 | + |
| Chr02 | 9945090 | 9947830 | 2741 | + |
| Chr02 | 10057368 | 10064412 | 7045 | - |
| Chr02 | 10073654 | 10081826 | 8173 | - |
| Chr02 | 10260432 | 10262805 | 2374 | + |
| Chr02 | 10414611 | 10421481 | 6871 | - |
| Chr02 | 10962133 | 10972727 | 10595 | - |
| Chr02 | 11007360 | 11012014 | 4655 | + |
| Chr02 | 11078050 | 11086896 | 8847 | + |
| Chr02 | 11096455 | 11106388 | 9934 | - |
| Chr02 | 11152459 | 11163124 | 10666 | + |
| Chr02 | 11152774 | 11163124 | 10351 | + |
| Chr02 | 11276655 | 11282010 | 5356 | - |
| Chr02 | 11587477 | 11592825 | 5349 | - |
| Chr02 | 11697046 | 11707536 | 10491 | + |
| Chr02 | 12057765 | 12069170 | 11406 | + |
| Chr02 | 12198528 | 12202832 | 4305 | + |
| Chr02 | 12201531 | 12210572 | 9042 | + |
| Chr02 | 12209092 | 12218804 | 9713 | + |
| Chr02 | 12525972 | 12532721 | 6750 | + |
| Chr02 | 12589035 | 12599666 | 10632 | + |
| Chr02 | 13048100 | 13066004 | 17905 | + |
| Chr02 | 13098119 | 13102863 | 4745 | - |
| Chr02 | 13633311 | 13638185 | 4875 | + |
| Chr02 | 13854843 | 13856806 | 1964 | + |
| Chr02 | 13944175 | 13950441 | 6267 | + |
| Chr02 | 14208848 | 14217596 | 8749 | + |
| Chr02 | 14625316 | 14633683 | 8368 | - |
| Chr02 | 15013929 | 15020512 | 6584 | - |
| Chr02 | 15066043 | 15070504 | 4462 | + |
| Chr02 | 15093962 | 15097065 | 3104 | - |
| Chr02 | 15118375 | 15123903 | 5529 | + |

| | | | | |
|---|---|---|---|---|
| Chr02 | 15132651 | 15139531 | 6881 | + |
| Chr02 | 15268909 | 15285889 | 16981 | - |
| Chr02 | 15271917 | 15279234 | 7318 | + |
| Chr02 | 15480804 | 15489175 | 8372 | + |
| Chr02 | 15612617 | 15619116 | 6500 | + |
| Chr02 | 15881924 | 15890863 | 8940 | + |
| Chr02 | 16495883 | 16507701 | 11819 | + |
| Chr02 | 16599055 | 16604232 | 5178 | + |
| Chr02 | 16971286 | 16975524 | 4239 | + |
| Chr02 | 17186391 | 17191847 | 5457 | + |
| Chr02 | 17284861 | 17293032 | 8172 | + |
| Chr02 | 17326927 | 17335010 | 8084 | - |
| Chr02 | 17339092 | 17347433 | 8342 | + |
| Chr02 | 17704891 | 17707725 | 2835 | + |
| Chr02 | 17994109 | 18004593 | 10485 | - |
| Chr02 | 18392299 | 18397731 | 5433 | - |
| Chr02 | 18408453 | 18415176 | 6724 | + |
| Chr02 | 18490632 | 18496025 | 5394 | + |
| Chr02 | 18605397 | 18614520 | 9124 | + |
| Chr02 | 18605397 | 18615704 | 10308 | + |
| Chr02 | 19305556 | 19312453 | 6898 | + |
| Chr02 | 19447371 | 19453014 | 5644 | + |
| Chr02 | 19460783 | 19473828 | 13046 | - |
| Chr02 | 19463398 | 19472062 | 8665 | + |
| Chr02 | 19621128 | 19626126 | 4999 | + |
| Chr02 | 19677873 | 19684479 | 6607 | + |
| Chr02 | 19855109 | 19862098 | 6990 | - |
| Chr02 | 19865311 | 19872312 | 7002 | + |
| Chr02 | 19877252 | 19884085 | 6834 | + |
| Chr02 | 19917945 | 19921029 | 3085 | + |
| Chr02 | 20024317 | 20033268 | 8952 | - |
| Chr02 | 20089891 | 20095996 | 6106 | + |

| | | | | |
|---|---|---|---|---|
| Chr02 | 20175405 | 20181517 | 6113 | + |
| Chr02 | 20327256 | 20333374 | 6119 | + |
| Chr02 | 20564796 | 20571017 | 6222 | + |
| Chr02 | 20606519 | 20627270 | 20752 | + |
| Chr02 | 20758647 | 20763821 | 5175 | + |
| Chr02 | 20794846 | 20799977 | 5132 | - |
| Chr02 | 20872647 | 20877649 | 5003 | + |
| Chr02 | 20925402 | 20927388 | 1987 | + |
| Chr02 | 20928736 | 20933842 | 5107 | + |
| Chr02 | 21067765 | 21073283 | 5519 | + |
| Chr02 | 21077773 | 21083181 | 5409 | + |
| Chr02 | 21167076 | 21172405 | 5330 | + |
| Chr02 | 21187531 | 21201562 | 14032 | + |
| Chr02 | 21257239 | 21262683 | 5445 | + |
| Chr02 | 21278068 | 21282848 | 4781 | + |
| Chr02 | 21339152 | 21344626 | 5475 | + |
| Chr02 | 21354062 | 21359601 | 5540 | + |
| Chr02 | 21475777 | 21481119 | 5343 | + |
| Chr02 | 21554433 | 21560035 | 5603 | + |
| Chr02 | 21574979 | 21592897 | 17919 | + |
| Chr02 | 21675952 | 21682635 | 6684 | + |
| Chr02 | 21709362 | 21714989 | 5628 | - |
| Chr02 | 21870918 | 21876267 | 5350 | + |
| Chr02 | 22127482 | 22136180 | 8699 | - |
| Chr02 | 22503056 | 22511843 | 8788 | + |
| Chr02 | 22519100 | 22521558 | 2459 | + |
| Chr02 | 22641585 | 22645446 | 3862 | - |
| Chr02 | 22798326 | 22804434 | 6109 | + |
| Chr02 | 22864098 | 22868569 | 4472 | - |
| Chr02 | 23198474 | 23203457 | 4984 | - |
| Chr02 | 23290550 | 23293815 | 3266 | + |
| Chr02 | 23290550 | 23294207 | 3658 | + |

| | | | | |
|---|---|---|---|---|
| Chr02 | 23290550 | 23294599 | 4050 | + |
| Chr02 | 23346300 | 23348376 | 2077 | + |
| Chr02 | 23446189 | 23452966 | 6778 | + |
| Chr02 | 23669758 | 23674950 | 5193 | + |
| Chr02 | 23689797 | 23695402 | 5606 | + |
| Chr02 | 23769872 | 23775134 | 5263 | + |
| Chr02 | 23801262 | 23808840 | 7579 | + |
| Chr02 | 23877170 | 23882356 | 5187 | + |
| Chr02 | 24157669 | 24164742 | 7074 | - |
| Chr02 | 24247749 | 24256273 | 8525 | - |
| Chr02 | 24248395 | 24255664 | 7270 | + |
| Chr02 | 24256959 | 24264383 | 7425 | - |
| Chr02 | 24367782 | 24373310 | 5529 | + |
| Chr02 | 24420758 | 24425152 | 4395 | + |
| Chr02 | 24421536 | 24425152 | 3617 | + |
| Chr02 | 24750650 | 24764580 | 13931 | - |
| Chr02 | 24753918 | 24781064 | 27147 | - |
| Chr02 | 24764746 | 24778707 | 13962 | + |
| Chr02 | 24966556 | 24971918 | 5363 | + |
| Chr02 | 25130415 | 25142308 | 11894 | - |
| Chr02 | 25414854 | 25420191 | 5338 | + |
| Chr02 | 25421589 | 25426861 | 5273 | + |
| Chr02 | 25466508 | 25478395 | 11888 | - |
| Chr02 | 25512870 | 25519573 | 6704 | - |
| Chr02 | 25756381 | 25763774 | 7394 | - |
| Chr02 | 25822233 | 25831374 | 9142 | + |
| Chr02 | 26138696 | 26148434 | 9739 | + |
| Chr02 | 26164073 | 26173583 | 9511 | + |
| Chr02 | 26184713 | 26190178 | 5466 | + |
| Chr02 | 26262704 | 26271027 | 8324 | + |
| Chr02 | 26306536 | 26317176 | 10641 | - |
| Chr02 | 26327032 | 26331712 | 4681 | - |

| | | | | |
|---|---|---|---|---|
| Chr02 | 26478037 | 26483017 | 4981 | + |
| Chr02 | 26501094 | 26521087 | 19994 | + |
| Chr02 | 26509031 | 26518152 | 9122 | + |
| Chr02 | 26583702 | 26590860 | 7159 | - |
| Chr02 | 26615041 | 26624608 | 9568 | - |
| Chr02 | 26837062 | 26843285 | 6224 | - |
| Chr02 | 27932100 | 27938638 | 6539 | - |
| Chr02 | 27947338 | 27952846 | 5509 | - |
| Chr02 | 27998677 | 28004549 | 5873 | + |
| Chr02 | 28040447 | 28051763 | 11317 | - |
| Chr02 | 28053670 | 28058152 | 4483 | - |
| Chr02 | 28073811 | 28079320 | 5510 | + |
| Chr02 | 28321633 | 28328122 | 6490 | + |
| Chr02 | 28545305 | 28550179 | 4875 | - |
| Chr02 | 29148973 | 29161489 | 12517 | - |
| Chr02 | 29499673 | 29504594 | 4922 | + |
| Chr02 | 29687224 | 29696528 | 9305 | + |
| Chr02 | 29916122 | 29923409 | 7288 | - |
| Chr02 | 29965996 | 29976131 | 10136 | + |
| Chr02 | 30000291 | 30004749 | 4459 | + |
| Chr02 | 30034751 | 30040224 | 5474 | + |
| Chr02 | 30320513 | 30345455 | 24943 | + |
| Chr02 | 30386248 | 30400530 | 14283 | + |
| Chr02 | 30391323 | 30395950 | 4628 | - |
| Chr02 | 30532370 | 30556023 | 23654 | + |
| Chr02 | 30662187 | 30668030 | 5844 | + |
| Chr02 | 30933619 | 30940709 | 7091 | + |
| Chr02 | 31005211 | 31010812 | 5602 | + |
| Chr02 | 31217774 | 31222951 | 5178 | - |
| Chr02 | 31299980 | 31307519 | 7540 | + |
| Chr02 | 31894214 | 31899889 | 5676 | + |
| Chr02 | 32058062 | 32064992 | 6931 | + |

| | | | | |
|---|---|---|---|---|
| Chr02 | 32127621 | 32134023 | 6403 | + |
| Chr02 | 32304223 | 32309312 | 5090 | - |
| Chr02 | 32736347 | 32741774 | 5428 | - |
| Chr02 | 32784484 | 32799458 | 14975 | + |
| Chr02 | 32829626 | 32835162 | 5537 | + |
| Chr02 | 32961391 | 32976905 | 15515 | - |
| Chr02 | 32962550 | 32983396 | 20847 | - |
| Chr02 | 33073237 | 33079552 | 6316 | + |
| Chr02 | 34025679 | 34030775 | 5097 | + |
| Chr02 | 34041289 | 34046813 | 5525 | + |
| Chr02 | 34080821 | 34086002 | 5182 | - |
| Chr02 | 34247813 | 34263318 | 15506 | + |
| Chr02 | 34252377 | 34262288 | 9912 | - |
| Chr02 | 34712062 | 34717084 | 5023 | + |
| Chr02 | 34732468 | 34734271 | 1804 | - |
| Chr02 | 34865314 | 34875828 | 10515 | - |
| Chr02 | 34978948 | 34984355 | 5408 | - |
| Chr02 | 35275914 | 35280670 | 4757 | + |
| Chr02 | 35478142 | 35483432 | 5291 | - |
| Chr02 | 35716354 | 35720821 | 4468 | + |
| Chr02 | 35826024 | 35834748 | 8725 | + |
| Chr02 | 35884184 | 35889621 | 5438 | - |
| Chr02 | 35904847 | 35910102 | 5256 | + |
| Chr02 | 35914955 | 35924278 | 9324 | + |
| Chr02 | 36473411 | 36484084 | 10674 | - |
| Chr02 | 36569511 | 36574572 | 5062 | + |
| Chr02 | 36595841 | 36606398 | 10558 | - |
| Chr02 | 36626469 | 36643167 | 16699 | + |
| Chr02 | 37389060 | 37393555 | 4496 | - |
| Chr02 | 37948180 | 37953186 | 5007 | - |
| Chr02 | 37974419 | 37979468 | 5050 | - |
| Chr02 | 38121392 | 38124030 | 2639 | + |

| | | | | |
|---|---|---|---|---|
| Chr02 | 38174549 | 38177776 | 3228 | + |
| Chr02 | 38554510 | 38556729 | 2220 | - |
| Chr02 | 38607939 | 38613296 | 5358 | - |
| Chr02 | 39014583 | 39019936 | 5354 | - |
| Chr02 | 39121608 | 39132898 | 11291 | + |
| Chr02 | 39138538 | 39144852 | 6315 | - |
| Chr02 | 39280137 | 39301314 | 21178 | + |
| Chr02 | 39286628 | 39296453 | 9826 | - |
| Chr02 | 39377364 | 39384953 | 7590 | + |
| Chr02 | 39434035 | 39438657 | 4623 | + |
| Chr02 | 39562071 | 39567212 | 5142 | - |
| Chr02 | 39625371 | 39636169 | 10799 | + |
| Chr02 | 39739423 | 39741327 | 1905 | - |
| Chr02 | 39743299 | 39749548 | 6250 | + |
| Chr02 | 39797686 | 39803119 | 5434 | + |
| Chr02 | 39812091 | 39819100 | 7010 | - |
| Chr02 | 39866638 | 39869890 | 3253 | - |
| Chr02 | 39907106 | 39912622 | 5517 | - |
| Chr02 | 39923614 | 39926900 | 3287 | - |
| Chr02 | 40087666 | 40111579 | 23914 | + |
| Chr02 | 40298584 | 40315187 | 16604 | + |
| Chr02 | 40352725 | 40357917 | 5193 | + |
| Chr02 | 40360914 | 40366421 | 5508 | - |
| Chr02 | 40640917 | 40654289 | 13373 | + |
| Chr02 | 40883901 | 40894619 | 10719 | + |
| Chr02 | 41532689 | 41537997 | 5309 | + |
| Chr02 | 42335380 | 42340907 | 5528 | - |
| Chr02 | 42335411 | 42340907 | 5497 | - |
| Chr02 | 43468361 | 43478030 | 9670 | - |
| Chr02 | 43536540 | 43548731 | 12192 | + |
| Chr02 | 43799861 | 43821583 | 21723 | + |
| Chr02 | 44120103 | 44131977 | 11875 | - |

| | | | | |
|---|---|---|---|---|
| Chr02 | 44121736 | 44131980 | 10245 | - |
| Chr02 | 44380120 | 44385283 | 5164 | + |
| Chr02 | 44890449 | 44892108 | 1660 | - |
| Chr02 | 44892428 | 44897315 | 4888 | - |
| Chr02 | 44901399 | 44906938 | 5540 | + |
| Chr02 | 45356752 | 45361761 | 5010 | + |
| Chr02 | 45567627 | 45572906 | 5280 | + |
| Chr02 | 45765030 | 45769264 | 4235 | + |
| Chr02 | 45872222 | 45876801 | 4580 | + |
| Chr02 | 45939576 | 45949054 | 9479 | + |
| Chr02 | 46112431 | 46117617 | 5187 | - |
| Chr02 | 46234497 | 46240148 | 5652 | - |
| Chr02 | 46246179 | 46254654 | 8476 | + |
| Chr02 | 46685133 | 46690326 | 5194 | - |
| Chr02 | 46695204 | 46698530 | 3327 | + |
| Chr02 | 47057068 | 47063671 | 6604 | + |
| Chr02 | 47216832 | 47242589 | 25758 | + |
| Chr02 | 47229052 | 47241111 | 12060 | - |
| Chr02 | 47418351 | 47424845 | 6495 | - |
| Chr02 | 47485892 | 47490371 | 4480 | + |
| Chr02 | 47679193 | 47684404 | 5212 | + |
| Chr02 | 47908213 | 47919264 | 11052 | + |
| Chr02 | 48407834 | 48413115 | 5282 | - |
| Chr02 | 49469874 | 49471888 | 2015 | + |
| Chr02 | 49471456 | 49473847 | 2392 | + |
| Chr03 | 201673 | 207207 | 5535 | - |
| Chr03 | 242983 | 248833 | 5851 | + |
| Chr03 | 343854 | 349121 | 5268 | - |
| Chr03 | 368866 | 371221 | 2356 | + |
| Chr03 | 542563 | 546625 | 4063 | + |
| Chr03 | 558264 | 565456 | 7193 | - |
| Chr03 | 622947 | 644781 | 21835 | - |

| | | | | |
|---|---|---|---|---|
| Chr03 | 718857 | 733128 | 14272 | - |
| Chr03 | 825966 | 830974 | 5009 | + |
| Chr03 | 909500 | 925761 | 16262 | + |
| Chr03 | 989409 | 995027 | 5619 | + |
| Chr03 | 1074708 | 1079738 | 5031 | + |
| Chr03 | 1719564 | 1725070 | 5507 | + |
| Chr03 | 1831050 | 1840937 | 9888 | - |
| Chr03 | 1909734 | 1920332 | 10599 | + |
| Chr03 | 2345228 | 2355999 | 10772 | + |
| Chr03 | 2399828 | 2405297 | 5470 | - |
| Chr03 | 2441987 | 2461394 | 19408 | + |
| Chr03 | 2443062 | 2449154 | 6093 | - |
| Chr03 | 2580751 | 2586164 | 5414 | + |
| Chr03 | 2619121 | 2645721 | 26601 | + |
| Chr03 | 2854560 | 2871669 | 17110 | + |
| Chr03 | 2953078 | 2967706 | 14629 | + |
| Chr03 | 2979004 | 2994600 | 15597 | - |
| Chr03 | 2982832 | 2988386 | 5555 | - |
| Chr03 | 3098809 | 3104192 | 5384 | + |
| Chr03 | 3236631 | 3241997 | 5367 | - |
| Chr03 | 3251278 | 3276896 | 25619 | - |
| Chr03 | 3386517 | 3391820 | 5304 | + |
| Chr03 | 3802371 | 3807602 | 5232 | + |
| Chr03 | 3857548 | 3867606 | 10059 | + |
| Chr03 | 3868610 | 3877467 | 8858 | + |
| Chr03 | 3880644 | 3903032 | 22389 | + |
| Chr03 | 3938294 | 3953053 | 14760 | - |
| Chr03 | 4179151 | 4189811 | 10661 | + |
| Chr03 | 4202054 | 4207489 | 5436 | + |
| Chr03 | 4311913 | 4315301 | 3389 | + |
| Chr03 | 4316061 | 4333696 | 17636 | + |
| Chr03 | 4408188 | 4431070 | 22883 | + |

| | | | | |
|---|---|---|---|---|
| Chr03 | 4495018 | 4501422 | 6405 | + |
| Chr03 | 4653751 | 4657979 | 4229 | - |
| Chr03 | 4853974 | 4879521 | 25548 | + |
| Chr03 | 4946747 | 4973328 | 26582 | - |
| Chr03 | 4965754 | 4973328 | 7575 | - |
| Chr03 | 5032608 | 5042631 | 10024 | - |
| Chr03 | 5250007 | 5254284 | 4278 | + |
| Chr03 | 5602161 | 5612037 | 9877 | - |
| Chr03 | 5877993 | 5883220 | 5228 | + |
| Chr03 | 5890938 | 5896170 | 5233 | - |
| Chr03 | 6155542 | 6160478 | 4937 | + |
| Chr03 | 6306347 | 6311508 | 5162 | - |
| Chr03 | 6458157 | 6476611 | 18455 | - |
| Chr03 | 6561105 | 6567801 | 6697 | - |
| Chr03 | 6607906 | 6612896 | 4991 | + |
| Chr03 | 6633535 | 6652311 | 18777 | - |
| Chr03 | 6637229 | 6651874 | 14646 | - |
| Chr03 | 6641348 | 6648913 | 7566 | - |
| Chr03 | 6685659 | 6698397 | 12739 | - |
| Chr03 | 6777420 | 6783999 | 6580 | + |
| Chr03 | 6789873 | 6796041 | 6169 | - |
| Chr03 | 6940579 | 6954802 | 14224 | - |
| Chr03 | 6945245 | 6954802 | 9558 | - |
| Chr03 | 7018185 | 7024609 | 6425 | + |
| Chr03 | 7198661 | 7204119 | 5459 | - |
| Chr03 | 7390410 | 7401027 | 10618 | + |
| Chr03 | 7504366 | 7510024 | 5659 | + |
| Chr03 | 7699265 | 7704655 | 5391 | + |
| Chr03 | 7711764 | 7717372 | 5609 | - |
| Chr03 | 8050797 | 8056121 | 5325 | - |
| Chr03 | 8231544 | 8235169 | 3626 | - |
| Chr03 | 8332366 | 8337589 | 5224 | + |

| | | | | |
|---|---|---|---|---|
| Chr03 | 8386127 | 8389892 | 3766 | + |
| Chr03 | 8902239 | 8908266 | 6028 | + |
| Chr03 | 9009452 | 9019953 | 10502 | + |
| Chr03 | 9045448 | 9050687 | 5240 | + |
| Chr03 | 9224762 | 9234754 | 9993 | - |
| Chr03 | 9314313 | 9325217 | 10905 | + |
| Chr03 | 9382777 | 9392482 | 9706 | + |
| Chr03 | 9414424 | 9424080 | 9657 | - |
| Chr03 | 9564139 | 9594885 | 30747 | + |
| Chr03 | 9708365 | 9714892 | 6528 | - |
| Chr03 | 9737309 | 9740690 | 3382 | - |
| Chr03 | 9815916 | 9832766 | 16851 | + |
| Chr03 | 9898494 | 9903739 | 5246 | + |
| Chr03 | 10489441 | 10494444 | 5004 | + |
| Chr03 | 10606940 | 10616918 | 9979 | - |
| Chr03 | 10783020 | 10789503 | 6484 | - |
| Chr03 | 10811522 | 10818670 | 7149 | + |
| Chr03 | 10849808 | 10854725 | 4918 | - |
| Chr03 | 10851125 | 10853236 | 2112 | + |
| Chr03 | 11034621 | 11036798 | 2178 | - |
| Chr03 | 11109750 | 11115490 | 5741 | - |
| Chr03 | 11109750 | 11116435 | 6686 | - |
| Chr03 | 11120779 | 11129667 | 8889 | - |
| Chr03 | 11249347 | 11250934 | 1588 | - |
| Chr03 | 11254054 | 11261061 | 7008 | + |
| Chr03 | 11696902 | 11702194 | 5293 | - |
| Chr03 | 11847246 | 11852693 | 5448 | + |
| Chr03 | 11876200 | 11898612 | 22413 | - |
| Chr03 | 11882907 | 11889689 | 6783 | + |
| Chr03 | 12164180 | 12169348 | 5169 | - |
| Chr03 | 12177127 | 12182393 | 5267 | + |
| Chr03 | 12688401 | 12693678 | 5278 | - |

| | | | | |
|---|---|---|---|---|
| Chr03 | 12773673 | 12788313 | 14641 | - |
| Chr03 | 12780229 | 12787181 | 6953 | + |
| Chr03 | 12802427 | 12811675 | 9249 | - |
| Chr03 | 12886490 | 12896914 | 10425 | - |
| Chr03 | 12886490 | 12905083 | 18594 | - |
| Chr03 | 12886490 | 12905413 | 18924 | - |
| Chr03 | 12886490 | 12905413 | 18924 | - |
| Chr03 | 12886573 | 12896914 | 10342 | - |
| Chr03 | 12886783 | 12896914 | 10132 | - |
| Chr03 | 12886783 | 12905413 | 18631 | - |
| Chr03 | 12896543 | 12905413 | 8871 | - |
| Chr03 | 13378096 | 13383701 | 5606 | + |
| Chr03 | 13391419 | 13397795 | 6377 | + |
| Chr03 | 13410902 | 13416266 | 5365 | - |
| Chr03 | 13443039 | 13445445 | 2407 | - |
| Chr03 | 13491822 | 13497024 | 5203 | + |
| Chr03 | 13547986 | 13552654 | 4669 | - |
| Chr03 | 13552711 | 13554432 | 1722 | - |
| Chr03 | 13802680 | 13809582 | 6903 | + |
| Chr03 | 13830598 | 13840709 | 10112 | - |
| Chr03 | 14096593 | 14101783 | 5191 | - |
| Chr03 | 14203399 | 14208889 | 5491 | + |
| Chr03 | 14277288 | 14283485 | 6198 | + |
| Chr03 | 14301918 | 14306512 | 4595 | + |
| Chr03 | 14369901 | 14375574 | 5674 | + |
| Chr03 | 14551840 | 14557620 | 5781 | + |
| Chr03 | 14607423 | 14612928 | 5506 | + |
| Chr03 | 14648200 | 14658735 | 10536 | + |
| Chr03 | 14697900 | 14708499 | 10600 | + |
| Chr03 | 15114102 | 15119637 | 5536 | + |
| Chr03 | 15122480 | 15128232 | 5753 | + |
| Chr03 | 15154350 | 15159883 | 5534 | + |

| | | | | |
|---|---|---|---|---|
| Chr03 | 15253682 | 15258575 | 4894 | - |
| Chr03 | 15445222 | 15453779 | 8558 | + |
| Chr03 | 15459689 | 15464870 | 5182 | - |
| Chr03 | 15674352 | 15682851 | 8500 | + |
| Chr03 | 16168170 | 16180882 | 12713 | - |
| Chr03 | 16451589 | 16457121 | 5533 | - |
| Chr03 | 16531805 | 16536923 | 5119 | + |
| Chr03 | 16538105 | 16567765 | 29661 | + |
| Chr03 | 16555615 | 16560455 | 4841 | + |
| Chr03 | 16623677 | 16630646 | 6970 | + |
| Chr03 | 16656124 | 16657922 | 1799 | - |
| Chr03 | 16674769 | 16685448 | 10680 | - |
| Chr03 | 16825202 | 16830640 | 5439 | - |
| Chr03 | 16839231 | 16849015 | 9785 | + |
| Chr03 | 16855781 | 16861194 | 5414 | - |
| Chr03 | 17011106 | 17015400 | 4295 | - |
| Chr03 | 17050020 | 17076033 | 26014 | - |
| Chr03 | 17057186 | 17063524 | 6339 | + |
| Chr03 | 17065866 | 17075415 | 9550 | - |
| Chr03 | 17276252 | 17282110 | 5859 | - |
| Chr03 | 17290497 | 17304882 | 14386 | - |
| Chr03 | 17472716 | 17481652 | 8937 | - |
| Chr03 | 17664358 | 17670026 | 5669 | + |
| Chr03 | 17692928 | 17707161 | 14234 | - |
| Chr03 | 17742264 | 17746636 | 4373 | + |
| Chr03 | 17838613 | 17846567 | 7955 | + |
| Chr03 | 18027849 | 18043619 | 15771 | - |
| Chr03 | 18351603 | 18356999 | 5397 | + |
| Chr03 | 18521798 | 18532695 | 10898 | + |
| Chr03 | 18566399 | 18574889 | 8491 | - |
| Chr03 | 20641938 | 20646737 | 4800 | + |
| Chr03 | 21555699 | 21561572 | 5874 | + |

| | | | | |
|---|---|---|---|---|
| Chr03 | 21981537 | 21986991 | 5455 | - |
| Chr03 | 23028642 | 23035336 | 6695 | + |
| Chr03 | 25455010 | 25458009 | 3000 | + |
| Chr03 | 27584849 | 27590672 | 5824 | - |
| Chr04 | 353109 | 358118 | 5010 | - |
| Chr04 | 668935 | 673875 | 4941 | - |
| Chr04 | 726552 | 733781 | 7230 | - |
| Chr04 | 749253 | 756368 | 7116 | - |
| Chr04 | 844948 | 855718 | 10771 | - |
| Chr04 | 1444787 | 1450263 | 5477 | + |
| Chr04 | 2228672 | 2234218 | 5547 | + |
| Chr04 | 2309054 | 2319309 | 10256 | + |
| Chr04 | 2361666 | 2367042 | 5377 | + |
| Chr04 | 2615661 | 2620477 | 4817 | - |
| Chr04 | 2615661 | 2633682 | 18022 | - |
| Chr04 | 2620229 | 2628990 | 8762 | + |
| Chr04 | 2628723 | 2633682 | 4960 | - |
| Chr04 | 3030630 | 3035856 | 5227 | - |
| Chr04 | 3388992 | 3395950 | 6959 | - |
| Chr04 | 3975700 | 3985651 | 9952 | + |
| Chr04 | 3996046 | 4002451 | 6406 | + |
| Chr04 | 4235795 | 4244246 | 8452 | + |
| Chr04 | 4267517 | 4272956 | 5440 | - |
| Chr04 | 4366181 | 4371074 | 4894 | + |
| Chr04 | 4601411 | 4607986 | 6576 | + |
| Chr04 | 4799728 | 4805550 | 5823 | + |
| Chr04 | 5853315 | 5863262 | 9948 | + |
| Chr04 | 5926655 | 5936498 | 9844 | + |
| Chr04 | 6431447 | 6438514 | 7068 | + |
| Chr04 | 6461062 | 6466264 | 5203 | + |
| Chr04 | 6495424 | 6500911 | 5488 | - |
| Chr04 | 6595009 | 6605282 | 10274 | + |

| | | | | |
|---|---|---|---|---|
| Chr04 | 6853466 | 6871224 | 17759 | - |
| Chr04 | 6873781 | 6880804 | 7024 | - |
| Chr04 | 7021203 | 7035043 | 13841 | + |
| Chr04 | 7022498 | 7033464 | 10967 | - |
| Chr04 | 7266820 | 7272042 | 5223 | - |
| Chr04 | 7473178 | 7480185 | 7008 | - |
| Chr04 | 7495823 | 7500846 | 5024 | + |
| Chr04 | 7692811 | 7699812 | 7002 | + |
| Chr04 | 7791141 | 7802683 | 11543 | + |
| Chr04 | 7827403 | 7838429 | 11027 | - |
| Chr04 | 8071270 | 8088762 | 17493 | + |
| Chr04 | 8576731 | 8588860 | 12130 | + |
| Chr04 | 8601241 | 8606281 | 5041 | + |
| Chr04 | 8609929 | 8616651 | 6723 | - |
| Chr04 | 9009548 | 9017150 | 7603 | - |
| Chr04 | 9945576 | 9958177 | 12602 | + |
| Chr04 | 10016793 | 10027816 | 11024 | + |
| Chr04 | 10182904 | 10192634 | 9731 | + |
| Chr04 | 10302495 | 10309582 | 7088 | + |
| Chr04 | 10374643 | 10379851 | 5209 | + |
| Chr04 | 10379905 | 10384477 | 4573 | + |
| Chr04 | 10421330 | 10427925 | 6596 | + |
| Chr04 | 10459886 | 10465465 | 5580 | + |
| Chr04 | 10522400 | 10527501 | 5102 | - |
| Chr04 | 10547718 | 10552620 | 4903 | + |
| Chr04 | 10915673 | 10925747 | 10075 | + |
| Chr04 | 11045199 | 11052131 | 6933 | + |
| Chr04 | 11093339 | 11118394 | 25056 | - |
| Chr04 | 11306801 | 11318688 | 11888 | + |
| Chr04 | 11983890 | 11994403 | 10514 | + |
| Chr04 | 12160050 | 12168839 | 8790 | + |
| Chr04 | 12286747 | 12299448 | 12702 | + |

| | | | | |
|---|---|---|---|---|
| Chr04 | 12376000 | 12383336 | 7337 | + |
| Chr04 | 12433841 | 12443133 | 9293 | - |
| Chr04 | 12518789 | 12531597 | 12809 | + |
| Chr04 | 12595274 | 12601771 | 6498 | - |
| Chr04 | 12700116 | 12708224 | 8109 | + |
| Chr04 | 12702006 | 12718718 | 16713 | + |
| Chr04 | 12708255 | 12724244 | 15990 | + |
| Chr04 | 12897212 | 12909482 | 12271 | + |
| Chr04 | 13095524 | 13105449 | 9926 | - |
| Chr04 | 13116716 | 13124723 | 8008 | + |
| Chr04 | 13143601 | 13149426 | 5826 | + |
| Chr04 | 13249108 | 13261136 | 12029 | - |
| Chr04 | 13476059 | 13482374 | 6316 | - |
| Chr04 | 14225982 | 14231571 | 5590 | + |
| Chr04 | 14316826 | 14327244 | 10419 | + |
| Chr04 | 14375286 | 14380526 | 5241 | + |
| Chr04 | 14601261 | 14603218 | 1958 | + |
| Chr04 | 14650209 | 14655220 | 5012 | + |
| Chr04 | 14755385 | 14768299 | 12915 | + |
| Chr04 | 14822906 | 14832573 | 9668 | + |
| Chr04 | 14973301 | 14982821 | 9521 | + |
| Chr04 | 15228820 | 15237335 | 8516 | + |
| Chr04 | 15400478 | 15406420 | 5943 | + |
| Chr04 | 15435612 | 15446228 | 10617 | - |
| Chr04 | 15515081 | 15536960 | 21880 | - |
| Chr04 | 15519135 | 15529909 | 10775 | + |
| Chr04 | 15590481 | 15600654 | 10174 | - |
| Chr04 | 15591559 | 15596418 | 4860 | - |
| Chr04 | 15623876 | 15628485 | 4610 | - |
| Chr04 | 15666793 | 15678643 | 11851 | + |
| Chr04 | 15733839 | 15737839 | 4001 | - |
| Chr04 | 15913195 | 15917895 | 4701 | - |

| | | | | |
|---|---|---|---|---|
| Chr04 | 16083629 | 16093977 | 10349 | - |
| Chr04 | 16979003 | 16992827 | 13825 | + |
| Chr04 | 17272726 | 17279064 | 6339 | + |
| Chr04 | 17408665 | 17414017 | 5353 | - |
| Chr04 | 17631274 | 17635757 | 4484 | + |
| Chr04 | 17641352 | 17643666 | 2315 | - |
| Chr04 | 17671655 | 17676650 | 4996 | + |
| Chr04 | 17690743 | 17695485 | 4743 | - |
| Chr04 | 17712197 | 17717209 | 5013 | - |
| Chr04 | 17787920 | 17793170 | 5251 | + |
| Chr04 | 17862957 | 17870373 | 7417 | - |
| Chr04 | 17901960 | 17915618 | 13659 | - |
| Chr04 | 18105331 | 18112333 | 7003 | + |
| Chr04 | 18602454 | 18612491 | 10038 | - |
| Chr04 | 18630834 | 18643801 | 12968 | - |
| Chr04 | 18660995 | 18665995 | 5001 | + |
| Chr04 | 18692666 | 18696242 | 3577 | - |
| Chr04 | 18752361 | 18757336 | 4976 | - |
| Chr04 | 18788603 | 18794713 | 6111 | + |
| Chr04 | 18806898 | 18812215 | 5318 | - |
| Chr04 | 18838203 | 18842866 | 4664 | + |
| Chr04 | 18945589 | 18953055 | 7467 | - |
| Chr04 | 19157451 | 19167003 | 9553 | - |
| Chr04 | 19386681 | 19393977 | 7297 | + |
| Chr04 | 20066451 | 20072195 | 5745 | - |
| Chr04 | 20111949 | 20119409 | 7461 | - |
| Chr04 | 20301737 | 20307627 | 5891 | + |
| Chr04 | 20413397 | 20414958 | 1562 | - |
| Chr04 | 20561899 | 20566994 | 5096 | - |
| Chr04 | 20609147 | 20610712 | 1566 | - |
| Chr04 | 20659473 | 20664780 | 5308 | - |
| Chr04 | 20709241 | 20714337 | 5097 | - |

| | | | | |
|---|---|---|---|---|
| Chr04 | 21037757 | 21046486 | 8730 | - |
| Chr04 | 21053057 | 21058173 | 5117 | + |
| Chr04 | 21079934 | 21085540 | 5607 | + |
| Chr04 | 21164444 | 21176505 | 12062 | - |
| Chr04 | 21202502 | 21215285 | 12784 | + |
| Chr04 | 21263926 | 21271440 | 7515 | + |
| Chr04 | 21277086 | 21283606 | 6521 | + |
| Chr04 | 21311628 | 21317281 | 5654 | + |
| Chr04 | 21511231 | 21516803 | 5573 | + |
| Chr04 | 21812508 | 21832059 | 19552 | - |
| Chr04 | 21867715 | 21872670 | 4956 | - |
| Chr04 | 21914496 | 21919445 | 4950 | - |
| Chr04 | 22161474 | 22166733 | 5260 | + |
| Chr04 | 23702813 | 23726844 | 24032 | + |
| Chr04 | 23894960 | 23905182 | 10223 | - |
| Chr04 | 23910563 | 23915658 | 5096 | - |
| Chr04 | 23992411 | 24016623 | 24213 | + |
| Chr04 | 24019992 | 24025425 | 5434 | - |
| Chr04 | 24278984 | 24283262 | 4279 | - |
| Chr04 | 24283328 | 24297070 | 13743 | - |
| Chr04 | 24399851 | 24410721 | 10871 | - |
| Chr04 | 24425471 | 24427711 | 2241 | - |
| Chr04 | 24676907 | 24682507 | 5601 | - |
| Chr04 | 24713025 | 24719340 | 6316 | + |
| Chr04 | 24743107 | 24748618 | 5512 | - |
| Chr04 | 24754595 | 24763346 | 8752 | - |
| Chr04 | 24793455 | 24800232 | 6778 | - |
| Chr04 | 24859171 | 24865205 | 6035 | - |
| Chr04 | 25046190 | 25051385 | 5196 | - |
| Chr04 | 25092665 | 25098160 | 5496 | + |
| Chr04 | 25607420 | 25614685 | 7266 | + |
| Chr04 | 25645909 | 25650895 | 4987 | + |

| | | | | |
|---|---|---|---|---|
| Chr04 | 25667016 | 25672035 | 5020 | - |
| Chr04 | 25895611 | 25905057 | 9447 | + |
| Chr04 | 26440554 | 26447249 | 6696 | + |
| Chr04 | 26923426 | 26928054 | 4629 | + |
| Chr04 | 27005038 | 27009646 | 4609 | + |
| Chr04 | 27237262 | 27239004 | 1743 | + |
| Chr04 | 27344359 | 27352711 | 8353 | - |
| Chr04 | 27344359 | 27354340 | 9982 | - |
| Chr04 | 27344359 | 27354340 | 9982 | - |
| Chr04 | 27344359 | 27354340 | 9982 | - |
| Chr04 | 27600570 | 27603597 | 3028 | - |
| Chr04 | 28336595 | 28359328 | 22734 | - |
| Chr04 | 28345599 | 28371779 | 26181 | - |
| Chr04 | 28353507 | 28359328 | 5822 | - |
| Chr04 | 28396804 | 28414332 | 17529 | - |
| Chr04 | 28401333 | 28415429 | 14097 | - |
| Chr04 | 28402672 | 28424791 | 22120 | - |
| Chr04 | 28414881 | 28424791 | 9911 | - |
| Chr04 | 28415296 | 28424422 | 9127 | - |
| Chr05 | 229422 | 238180 | 8759 | + |
| Chr05 | 273670 | 278691 | 5022 | - |
| Chr05 | 335851 | 346829 | 10979 | + |
| Chr05 | 480579 | 486044 | 5466 | + |
| Chr05 | 1046170 | 1050735 | 4566 | - |
| Chr05 | 1224900 | 1230465 | 5566 | - |
| Chr05 | 1419295 | 1423898 | 4604 | - |
| Chr05 | 2111593 | 2114944 | 3352 | - |
| Chr05 | 2194945 | 2200602 | 5658 | - |
| Chr05 | 2202796 | 2210878 | 8083 | + |
| Chr05 | 2364248 | 2369760 | 5513 | - |
| Chr05 | 2867191 | 2874456 | 7266 | + |
| Chr05 | 2913317 | 2918828 | 5512 | + |

| | | | | |
|---|---|---|---|---|
| Chr05 | 2964741 | 2970086 | 5346 | - |
| Chr05 | 3481671 | 3486616 | 4946 | + |
| Chr05 | 3666062 | 3673348 | 7287 | + |
| Chr05 | 4254413 | 4259935 | 5523 | - |
| Chr05 | 4725882 | 4734291 | 8410 | + |
| Chr05 | 4953782 | 4957455 | 3674 | + |
| Chr05 | 5327891 | 5332626 | 4736 | + |
| Chr05 | 5372815 | 5378200 | 5386 | + |
| Chr05 | 5758774 | 5764338 | 5565 | + |
| Chr05 | 5829379 | 5854798 | 25420 | - |
| Chr05 | 5834293 | 5841597 | 7305 | - |
| Chr05 | 6874414 | 6879217 | 4804 | - |
| Chr05 | 6931374 | 6943428 | 12055 | + |
| Chr05 | 6981433 | 6983504 | 2072 | - |
| Chr05 | 7093413 | 7098840 | 5428 | - |
| Chr05 | 7104251 | 7109539 | 5289 | + |
| Chr05 | 7514395 | 7518784 | 4390 | - |
| Chr05 | 8598702 | 8603866 | 5165 | - |
| Chr05 | 8878389 | 8883756 | 5368 | + |
| Chr05 | 9075515 | 9084140 | 8626 | - |
| Chr05 | 9342184 | 9349130 | 6947 | + |
| Chr05 | 9611652 | 9628920 | 17269 | + |
| Chr05 | 10502101 | 10507475 | 5375 | - |
| Chr05 | 10768402 | 10776005 | 7604 | + |
| Chr05 | 10876295 | 10881375 | 5081 | + |
| Chr05 | 11047791 | 11054848 | 7058 | + |
| Chr05 | 11190008 | 11197717 | 7710 | + |
| Chr05 | 11257254 | 11269436 | 12183 | + |
| Chr05 | 11522980 | 11541008 | 18029 | - |
| Chr05 | 11618199 | 11631723 | 13525 | + |
| Chr05 | 11866390 | 11884839 | 18450 | + |
| Chr05 | 11962240 | 11972351 | 10112 | + |

| | | | | |
|---|---|---|---|---|
| Chr05 | 12006920 | 12016336 | 9417 | + |
| Chr05 | 12199877 | 12209635 | 9759 | + |
| Chr05 | 12210221 | 12212036 | 1816 | + |
| Chr05 | 12233157 | 12238662 | 5506 | + |
| Chr05 | 12287627 | 12298362 | 10736 | - |
| Chr05 | 12339604 | 12347839 | 8236 | + |
| Chr05 | 12401868 | 12406339 | 4472 | + |
| Chr05 | 12501817 | 12506949 | 5133 | - |
| Chr05 | 12515550 | 12520866 | 5317 | + |
| Chr05 | 12695201 | 12700271 | 5071 | - |
| Chr05 | 12875017 | 12880990 | 5974 | + |
| Chr05 | 13200996 | 13206399 | 5404 | + |
| Chr05 | 13454325 | 13459765 | 5441 | - |
| Chr05 | 13777095 | 13785302 | 8208 | + |
| Chr05 | 13799400 | 13804696 | 5297 | - |
| Chr05 | 14212492 | 14224354 | 11863 | + |
| Chr05 | 14232834 | 14245227 | 12394 | - |
| Chr05 | 14269606 | 14281648 | 12043 | + |
| Chr05 | 14282971 | 14288364 | 5394 | - |
| Chr05 | 14333646 | 14340705 | 7060 | + |
| Chr05 | 14363448 | 14382832 | 19385 | + |
| Chr05 | 14435764 | 14443582 | 7819 | + |
| Chr05 | 15013446 | 15032539 | 19094 | - |
| Chr05 | 15034772 | 15057322 | 22551 | - |
| Chr05 | 15185398 | 15190884 | 5487 | - |
| Chr05 | 15408968 | 15414329 | 5362 | + |
| Chr05 | 15417260 | 15422689 | 5430 | + |
| Chr05 | 15436112 | 15441700 | 5589 | + |
| Chr05 | 15446618 | 15453212 | 6595 | + |
| Chr05 | 15624797 | 15645029 | 20233 | - |
| Chr05 | 15639772 | 15656689 | 16918 | - |
| Chr05 | 15761807 | 15767111 | 5305 | - |

| | | | | |
|---|---|---|---|---|
| Chr05 | 15904678 | 15913229 | 8552 | + |
| Chr05 | 15922074 | 15927126 | 5053 | - |
| Chr05 | 15976021 | 15981498 | 5478 | - |
| Chr05 | 16139843 | 16151952 | 12110 | + |
| Chr05 | 16452066 | 16453837 | 1772 | - |
| Chr05 | 16562824 | 16569273 | 6450 | - |
| Chr05 | 16884128 | 16893823 | 9696 | + |
| Chr05 | 17005850 | 17010060 | 4211 | - |
| Chr05 | 17140520 | 17150133 | 9614 | - |
| Chr05 | 17391468 | 17396791 | 5324 | - |
| Chr05 | 17570627 | 17577193 | 6567 | + |
| Chr05 | 17757136 | 17759637 | 2502 | + |
| Chr05 | 17776671 | 17779410 | 2740 | + |
| Chr05 | 17776671 | 17779814 | 3144 | + |
| Chr05 | 17776671 | 17781896 | 5226 | + |
| Chr05 | 18239966 | 18250674 | 10709 | - |
| Chr05 | 18799647 | 18816765 | 17119 | + |
| Chr05 | 18802595 | 18806234 | 3640 | - |
| Chr05 | 18821187 | 18822897 | 1711 | - |
| Chr05 | 18854291 | 18859630 | 5340 | - |
| Chr05 | 19545389 | 19566421 | 21033 | + |
| Chr05 | 19545389 | 19567418 | 22030 | + |
| Chr05 | 19545883 | 19567418 | 21536 | + |
| Chr05 | 19576342 | 19581654 | 5313 | - |
| Chr05 | 19936956 | 19947358 | 10403 | + |
| Chr05 | 19960433 | 19965616 | 5184 | - |
| Chr05 | 19981088 | 19986474 | 5387 | + |
| Chr05 | 20104069 | 20115025 | 10957 | - |
| Chr05 | 20108540 | 20115025 | 6486 | - |
| Chr05 | 20157858 | 20168838 | 10981 | - |
| Chr05 | 20164828 | 20168838 | 4011 | - |
| Chr05 | 20280780 | 20287253 | 6474 | - |

| | | | | |
|---|---|---|---|---|
| Chr05 | 20297336 | 20306455 | 9120 | + |
| Chr05 | 20830892 | 20842062 | 11171 | - |
| Chr05 | 20832730 | 20840177 | 7448 | - |
| Chr05 | 20849167 | 20854560 | 5394 | - |
| Chr05 | 21290417 | 21308840 | 18424 | - |
| Chr05 | 21471357 | 21484886 | 13530 | - |
| Chr05 | 21527685 | 21537373 | 9689 | - |
| Chr05 | 21527877 | 21536322 | 8446 | + |
| Chr05 | 21571115 | 21583167 | 12053 | - |
| Chr05 | 21640544 | 21653114 | 12571 | + |
| Chr05 | 21699657 | 21716278 | 16622 | - |
| Chr05 | 21732436 | 21735647 | 3212 | - |
| Chr05 | 21876647 | 21878827 | 2181 | + |
| Chr05 | 21882874 | 21898305 | 15432 | - |
| Chr05 | 21913306 | 21916759 | 3454 | - |
| Chr05 | 21977777 | 21990498 | 12722 | + |
| Chr05 | 21983309 | 21990409 | 7101 | - |
| Chr05 | 21984280 | 21990498 | 6219 | - |
| Chr05 | 22056501 | 22068017 | 11517 | + |
| Chr05 | 22092024 | 22113026 | 21003 | + |
| Chr05 | 22099357 | 22113158 | 13802 | + |
| Chr05 | 22102084 | 22116480 | 14397 | + |
| Chr05 | 22102091 | 22115276 | 13186 | + |
| Chr05 | 22102431 | 22116480 | 14050 | + |
| Chr05 | 22103718 | 22116960 | 13243 | + |
| Chr05 | 22118903 | 22122244 | 3342 | + |
| Chr05 | 22156637 | 22160096 | 3460 | + |
| Chr05 | 22225823 | 22240499 | 14677 | - |
| Chr05 | 22262133 | 22277436 | 15304 | - |
| Chr05 | 22316925 | 22321539 | 4615 | + |
| Chr05 | 22317503 | 22331821 | 14319 | - |
| Chr05 | 22317503 | 22331821 | 14319 | - |

| | | | | |
|---|---|---|---|---|
| Chr05 | 22317520 | 22322781 | 5262 | + |
| Chr05 | 22317520 | 22323716 | 6197 | + |
| Chr05 | 22317520 | 22327524 | 10005 | + |
| Chr05 | 22317520 | 22327858 | 10339 | + |
| Chr05 | 22317520 | 22329146 | 11627 | - |
| Chr05 | 22317520 | 22329776 | 12257 | + |
| Chr05 | 22317520 | 22331390 | 13871 | + |
| Chr05 | 22317786 | 22320662 | 2877 | + |
| Chr05 | 22317786 | 22327884 | 10099 | + |
| Chr05 | 22317786 | 22328185 | 10400 | + |
| Chr05 | 22317786 | 22329172 | 11387 | + |
| Chr05 | 22317807 | 22331821 | 14015 | + |
| Chr05 | 22317807 | 22331821 | 14015 | - |
| Chr05 | 22317807 | 22331821 | 14015 | - |
| Chr05 | 22317824 | 22319349 | 1526 | + |
| Chr05 | 22317824 | 22327858 | 10035 | + |
| Chr05 | 22317824 | 22329146 | 11323 | + |
| Chr05 | 22318154 | 22331692 | 13539 | - |
| Chr05 | 22318768 | 22331821 | 13054 | - |
| Chr05 | 22318768 | 22331821 | 13054 | - |
| Chr05 | 22318785 | 22321870 | 3086 | + |
| Chr05 | 22318785 | 22327858 | 9074 | - |
| Chr05 | 22319100 | 22331821 | 12722 | - |
| Chr05 | 22319117 | 22330077 | 10961 | - |
| Chr05 | 22319117 | 22331692 | 12576 | - |
| Chr05 | 22319379 | 22329172 | 9794 | + |
| Chr05 | 22319379 | 22329802 | 10424 | + |
| Chr05 | 22319400 | 22331821 | 12422 | - |
| Chr05 | 22319400 | 22331821 | 12422 | + |
| Chr05 | 22319400 | 22331821 | 12422 | + |
| Chr05 | 22319400 | 22331821 | 12422 | - |
| Chr05 | 22319417 | 22322172 | 2756 | + |

| | | | | |
|---|---|---|---|---|
| Chr05 | 22319417 | 22327858 | 8442 | + |
| Chr05 | 22319417 | 22329776 | 10360 | + |
| Chr05 | 22319417 | 22329776 | 10360 | + |
| Chr05 | 22319417 | 22330077 | 10661 | + |
| Chr05 | 22319746 | 22322473 | 2728 | - |
| Chr05 | 22319746 | 22322781 | 3036 | - |
| Chr05 | 22319746 | 22330077 | 10332 | + |
| Chr05 | 22320667 | 22328513 | 7847 | + |
| Chr05 | 22320688 | 22331821 | 11134 | - |
| Chr05 | 22320705 | 22327194 | 6490 | - |
| Chr05 | 22321901 | 22329802 | 7902 | + |
| Chr05 | 22321901 | 22330431 | 8531 | + |
| Chr05 | 22321901 | 22330759 | 8859 | + |
| Chr05 | 22321901 | 22330759 | 8859 | + |
| Chr05 | 22321922 | 22331821 | 9900 | - |
| Chr05 | 22321922 | 22331821 | 9900 | + |
| Chr05 | 22321922 | 22331821 | 9900 | + |
| Chr05 | 22321922 | 22331821 | 9900 | - |
| Chr05 | 22321939 | 22329446 | 7508 | + |
| Chr05 | 22321939 | 22329776 | 7838 | + |
| Chr05 | 22321939 | 22330077 | 8139 | + |
| Chr05 | 22321939 | 22330405 | 8467 | + |
| Chr05 | 22322203 | 22331088 | 8886 | + |
| Chr05 | 22322203 | 22331088 | 8886 | + |
| Chr05 | 22322203 | 22331416 | 9214 | + |
| Chr05 | 22322224 | 22331821 | 9598 | + |
| Chr05 | 22322224 | 22331821 | 9598 | + |
| Chr05 | 22322241 | 22327194 | 4954 | + |
| Chr05 | 22322241 | 22327524 | 5284 | + |
| Chr05 | 22322241 | 22328816 | 6576 | + |
| Chr05 | 22322241 | 22331390 | 9150 | + |
| Chr05 | 22323158 | 22331821 | 8664 | - |

| | | | | |
|---|---|---|---|---|
| Chr05 | 22323180 | 22328159 | 4980 | + |
| Chr05 | 22323180 | 22329446 | 6267 | + |
| Chr05 | 22323180 | 22329776 | 6597 | + |
| Chr05 | 22323180 | 22330077 | 6898 | + |
| Chr05 | 22323484 | 22328816 | 5333 | + |
| Chr05 | 22323747 | 22328185 | 4439 | + |
| Chr05 | 22323747 | 22328842 | 5096 | + |
| Chr05 | 22323747 | 22329172 | 5426 | + |
| Chr05 | 22323747 | 22329802 | 6056 | + |
| Chr05 | 22323747 | 22330759 | 7013 | + |
| Chr05 | 22323747 | 22331088 | 7342 | + |
| Chr05 | 22323747 | 22331718 | 7972 | + |
| Chr05 | 22323747 | 22331718 | 7972 | + |
| Chr05 | 22323768 | 22331821 | 8054 | + |
| Chr05 | 22323768 | 22331821 | 8054 | + |
| Chr05 | 22323768 | 22331821 | 8054 | - |
| Chr05 | 22323768 | 22331821 | 8054 | - |
| Chr05 | 22323785 | 22325579 | 1795 | + |
| Chr05 | 22323785 | 22327524 | 3740 | + |
| Chr05 | 22323785 | 22327858 | 4074 | + |
| Chr05 | 22323785 | 22328159 | 4375 | + |
| Chr05 | 22323785 | 22328816 | 5032 | + |
| Chr05 | 22323785 | 22329146 | 5362 | + |
| Chr05 | 22323785 | 22329146 | 5362 | + |
| Chr05 | 22323785 | 22329776 | 5992 | + |
| Chr05 | 22323785 | 22330077 | 6293 | + |
| Chr05 | 22323785 | 22330405 | 6621 | + |
| Chr05 | 22324377 | 22329172 | 4796 | + |
| Chr05 | 22324377 | 22329802 | 5426 | + |
| Chr05 | 22324377 | 22331718 | 7342 | + |
| Chr05 | 22324398 | 22331821 | 7424 | + |
| Chr05 | 22324398 | 22331821 | 7424 | + |

| | | | | |
|---|---|---|---|---|
| Chr05 | 22324398 | 22331821 | 7424 | - |
| Chr05 | 22324415 | 22326209 | 1795 | + |
| Chr05 | 22324415 | 22327524 | 3110 | + |
| Chr05 | 22324415 | 22329146 | 4732 | + |
| Chr05 | 22324415 | 22329776 | 5362 | + |
| Chr05 | 22324415 | 22330405 | 5991 | + |
| Chr05 | 22324677 | 22328185 | 3509 | + |
| Chr05 | 22324698 | 22331821 | 7124 | + |
| Chr05 | 22324698 | 22331821 | 7124 | - |
| Chr05 | 22324715 | 22326209 | 1495 | + |
| Chr05 | 22324715 | 22327524 | 2810 | + |
| Chr05 | 22324715 | 22327858 | 3144 | + |
| Chr05 | 22324715 | 22329146 | 4432 | + |
| Chr05 | 22324715 | 22329776 | 5062 | + |
| Chr05 | 22324715 | 22331390 | 6676 | + |
| Chr05 | 22324715 | 22331390 | 6676 | + |
| Chr05 | 22324978 | 22327884 | 2907 | + |
| Chr05 | 22324978 | 22329172 | 4195 | + |
| Chr05 | 22324978 | 22329802 | 4825 | + |
| Chr05 | 22324999 | 22331821 | 6823 | - |
| Chr05 | 22325016 | 22327524 | 2509 | + |
| Chr05 | 22325016 | 22327858 | 2843 | + |
| Chr05 | 22325016 | 22331390 | 6375 | + |
| Chr05 | 22325347 | 22330077 | 4731 | - |
| Chr05 | 22325610 | 22327884 | 2275 | + |
| Chr05 | 22325610 | 22329802 | 4193 | + |
| Chr05 | 22325631 | 22331821 | 6191 | + |
| Chr05 | 22325631 | 22331821 | 6191 | + |
| Chr05 | 22325648 | 22329776 | 4129 | + |
| Chr05 | 22327292 | 22330405 | 3114 | - |
| Chr05 | 22327292 | 22331390 | 4099 | - |
| Chr05 | 22328567 | 22331821 | 3255 | - |

| | | | | |
|---|---|---|---|---|
| Chr05 | 22338802 | 22360198 | 21397 | - |
| Chr05 | 22341478 | 22360198 | 18721 | - |
| Chr05 | 22342250 | 22366752 | 24503 | - |
| Chr05 | 22342272 | 22359313 | 17042 | - |
| Chr05 | 22847272 | 22852701 | 5430 | + |
| Chr05 | 22935121 | 22940708 | 5588 | + |
| Chr05 | 22970372 | 22985230 | 14859 | + |
| Chr05 | 23054936 | 23060382 | 5447 | + |
| Chr05 | 23428110 | 23434125 | 6016 | + |
| Chr05 | 23484153 | 23495472 | 11320 | - |
| Chr05 | 23522437 | 23548796 | 26360 | + |
| Chr05 | 23625463 | 23635029 | 9567 | + |
| Chr05 | 23711868 | 23718789 | 6922 | + |
| Chr05 | 23742964 | 23755839 | 12876 | + |
| Chr05 | 23895061 | 23901175 | 6115 | - |
| Chr05 | 23981916 | 23987211 | 5296 | - |
| Chr05 | 24429657 | 24442520 | 12864 | - |
| Chr05 | 24898698 | 24904100 | 5403 | + |
| Chr05 | 24950792 | 24957465 | 6674 | + |
| Chr05 | 25307231 | 25318453 | 11223 | - |
| Chr05 | 25363246 | 25368626 | 5381 | + |
| Chr05 | 25455737 | 25462494 | 6758 | + |
| Chr05 | 25611025 | 25622029 | 11005 | - |
| Chr05 | 25663335 | 25675463 | 12129 | - |
| Chr05 | 25862547 | 25887454 | 24908 | + |
| Chr05 | 26226908 | 26233883 | 6976 | + |
| Chr05 | 26302254 | 26321501 | 19248 | - |
| Chr05 | 26355532 | 26372491 | 16960 | - |
| Chr05 | 26418378 | 26423113 | 4736 | + |
| Chr05 | 26646373 | 26651909 | 5537 | + |
| Chr05 | 26721738 | 26728931 | 7194 | + |
| Chr05 | 26855502 | 26858429 | 2928 | - |

| | | | | |
|---|---|---|---|---|
| Chr05 | 27141277 | 27150884 | 9608 | + |
| Chr05 | 27256539 | 27262182 | 5644 | + |
| Chr05 | 27499133 | 27509189 | 10057 | - |
| Chr05 | 27555960 | 27565855 | 9896 | - |
| Chr05 | 27874928 | 27878555 | 3628 | + |
| Chr05 | 27890126 | 27895260 | 5135 | + |
| Chr05 | 27975468 | 27980894 | 5427 | + |
| Chr05 | 28048863 | 28053412 | 4550 | + |
| Chr05 | 28060130 | 28068743 | 8614 | + |
| Chr05 | 28633407 | 28641470 | 8064 | + |
| Chr05 | 28949443 | 28953820 | 4378 | - |
| Chr05 | 28959155 | 28966165 | 7011 | - |
| Chr05 | 28971641 | 28978202 | 6562 | + |
| Chr05 | 29056400 | 29062831 | 6432 | + |
| Chr05 | 29267311 | 29272315 | 5005 | + |
| Chr05 | 29664023 | 29669405 | 5383 | - |
| Chr05 | 29832915 | 29840005 | 7091 | - |
| Chr05 | 29850842 | 29858180 | 7339 | + |
| Chr05 | 30028748 | 30040488 | 11741 | - |
| Chr05 | 30098334 | 30106450 | 8117 | + |
| Chr05 | 30444884 | 30454469 | 9586 | - |
| Chr05 | 30700457 | 30709106 | 8650 | + |
| Chr05 | 30750673 | 30768595 | 17923 | + |
| Chr05 | 30759894 | 30765713 | 5820 | + |
| Chr05 | 30761864 | 30767574 | 5711 | + |
| Chr05 | 30761864 | 30767857 | 5994 | + |
| Chr05 | 30805767 | 30819880 | 14114 | + |
| Chr05 | 30805767 | 30820162 | 14396 | + |
| Chr05 | 30834142 | 30841420 | 7279 | + |
| Chr05 | 30880608 | 30883425 | 2818 | + |
| Chr05 | 30920810 | 30925564 | 4755 | - |
| Chr05 | 30949730 | 30954255 | 4526 | + |

| | | | | |
|---|---|---|---|---|
| Chr05 | 31383147 | 31400590 | 17444 | - |
| Chr05 | 31399476 | 31412735 | 13260 | - |
| Chr05 | 31412096 | 31420485 | 8390 | - |
| Chr05 | 31590309 | 31600510 | 10202 | + |
| Chr05 | 31595105 | 31600510 | 5406 | + |
| Chr05 | 31595105 | 31600733 | 5629 | + |
| Chr05 | 31660492 | 31671971 | 11480 | + |
| Chr05 | 31725661 | 31732382 | 6722 | - |
| Chr05 | 31800615 | 31806521 | 5907 | + |
| Chr05 | 32053195 | 32058278 | 5084 | + |
| Chr05 | 32061247 | 32065728 | 4482 | + |
| Chr05 | 32103671 | 32109244 | 5574 | + |
| Chr05 | 32144364 | 32149616 | 5253 | + |
| Chr05 | 32168071 | 32173331 | 5261 | - |
| Chr05 | 32248474 | 32253065 | 4592 | + |
| Chr05 | 32279483 | 32284970 | 5488 | + |
| Chr05 | 32468032 | 32472777 | 4746 | + |
| Chr05 | 32519084 | 32527957 | 8874 | + |
| Chr05 | 32567027 | 32571608 | 4582 | - |
| Chr05 | 32622402 | 32632254 | 9853 | + |
| Chr05 | 32694923 | 32699924 | 5002 | + |
| Chr05 | 32752061 | 32757230 | 5170 | + |
| Chr05 | 32802594 | 32808852 | 6259 | - |
| Chr05 | 33037791 | 33042953 | 5163 | - |
| Chr05 | 33037791 | 33049187 | 11397 | - |
| Chr05 | 33053826 | 33066755 | 12930 | + |
| Chr05 | 33709433 | 33715056 | 5624 | + |
| Chr05 | 34156250 | 34164604 | 8355 | + |
| Chr05 | 34265814 | 34291930 | 26117 | + |
| Chr05 | 34267808 | 34269419 | 1612 | + |
| Chr05 | 34268655 | 34286769 | 18115 | + |
| Chr05 | 34269429 | 34278778 | 9350 | + |

| | | | | |
|---|---|---|---|---|
| Chr05 | 34327644 | 34337127 | 9484 | + |
| Chr05 | 34345947 | 34361619 | 15673 | - |
| Chr05 | 34383920 | 34389098 | 5179 | + |
| Chr05 | 34460578 | 34469959 | 9382 | + |
| Chr05 | 34886716 | 34891726 | 5011 | - |
| Chr05 | 34947069 | 34952640 | 5572 | + |
| Chr05 | 35035956 | 35041537 | 5582 | - |
| Chr05 | 35202820 | 35209131 | 6312 | + |
| Chr05 | 35388721 | 35416070 | 27350 | + |
| Chr05 | 35421553 | 35447912 | 26360 | + |
| Chr05 | 35439495 | 35444872 | 5378 | + |
| Chr05 | 35615539 | 35621176 | 5638 | + |
| Chr05 | 35838534 | 35840916 | 2383 | - |
| Chr05 | 35840483 | 35844665 | 4183 | + |
| Chr05 | 35840483 | 35845319 | 4837 | + |
| Chr05 | 35889276 | 35894827 | 5552 | - |
| Chr05 | 35940720 | 35948803 | 8084 | - |
| Chr05 | 36427690 | 36433121 | 5432 | - |
| Chr05 | 36592893 | 36598150 | 5258 | - |
| Chr05 | 36610234 | 36617918 | 7685 | + |
| Chr05 | 36753037 | 36758571 | 5535 | + |
| Chr05 | 36799151 | 36804366 | 5216 | - |
| Chr05 | 36819798 | 36825735 | 5938 | - |
| Chr05 | 36843894 | 36864784 | 20891 | - |
| Chr05 | 36869375 | 36874747 | 5373 | - |
| Chr05 | 36879771 | 36884350 | 4580 | - |
| Chr05 | 36942882 | 36949101 | 6220 | + |
| Chr05 | 37188028 | 37193696 | 5669 | + |
| Chr05 | 37635818 | 37641962 | 6145 | + |
| Chr05 | 37786873 | 37793322 | 6450 | - |
| Chr05 | 37915763 | 37921087 | 5325 | - |
| Chr05 | 38865182 | 38872405 | 7224 | + |

| | | | | |
|---|---|---|---|---|
| Chr05 | 38929613 | 38932019 | 2407 | - |
| Chr05 | 40137729 | 40142827 | 5099 | - |
| Chr05 | 40159212 | 40168857 | 9646 | + |
| Chr05 | 40975390 | 40978634 | 3245 | - |
| Chr05 | 41198393 | 41201702 | 3310 | - |
| Chr05 | 42614260 | 42624097 | 9838 | + |
| Chr05 | 42618740 | 42623666 | 4927 | + |
| Chr05 | 44579637 | 44585066 | 5430 | - |
| Chr05 | 45085429 | 45091007 | 5579 | + |
| Chr05 | 45176409 | 45181930 | 5522 | - |
| Chr06 | 24685 | 34340 | 9656 | - |
| Chr06 | 63485 | 72396 | 8912 | + |
| Chr06 | 87994 | 93240 | 5247 | + |
| Chr06 | 131850 | 143250 | 11401 | - |
| Chr06 | 253357 | 258673 | 5317 | + |
| Chr06 | 285840 | 294209 | 8370 | + |
| Chr06 | 298453 | 301449 | 2997 | - |
| Chr06 | 422974 | 428248 | 5275 | + |
| Chr06 | 451210 | 455149 | 3940 | - |
| Chr06 | 653391 | 659529 | 6139 | + |
| Chr06 | 733340 | 741620 | 8281 | - |
| Chr06 | 736022 | 741205 | 5184 | + |
| Chr06 | 869365 | 872208 | 2844 | + |
| Chr06 | 869831 | 872208 | 2378 | - |
| Chr06 | 873009 | 874924 | 1916 | + |
| Chr06 | 1176104 | 1182982 | 6879 | - |
| Chr06 | 1216777 | 1222000 | 5224 | - |
| Chr06 | 1340777 | 1351805 | 11029 | + |
| Chr06 | 1535763 | 1541889 | 6127 | + |
| Chr06 | 1581275 | 1585711 | 4437 | - |
| Chr06 | 1861032 | 1867409 | 6378 | + |
| Chr06 | 1883784 | 1888731 | 4948 | - |

| | | | | |
|---|---|---|---|---|
| Chr06 | 1991119 | 1997473 | 6355 | + |
| Chr06 | 2061308 | 2069697 | 8390 | + |
| Chr06 | 2317609 | 2330335 | 12727 | - |
| Chr06 | 2505079 | 2531630 | 26552 | - |
| Chr06 | 2576088 | 2581809 | 5722 | - |
| Chr06 | 2782140 | 2790946 | 8807 | - |
| Chr06 | 2854718 | 2858515 | 3798 | + |
| Chr06 | 2897821 | 2903318 | 5498 | - |
| Chr06 | 2978275 | 2988130 | 9856 | - |
| Chr06 | 3075181 | 3082937 | 7757 | + |
| Chr06 | 3258812 | 3264250 | 5439 | + |
| Chr06 | 3269782 | 3279280 | 9499 | + |
| Chr06 | 3290476 | 3295363 | 4888 | - |
| Chr06 | 3299293 | 3305448 | 6156 | - |
| Chr06 | 3305464 | 3315325 | 9862 | + |
| Chr06 | 3327026 | 3337281 | 10256 | - |
| Chr06 | 3628446 | 3635658 | 7213 | + |
| Chr06 | 3760823 | 3767531 | 6709 | + |
| Chr06 | 4295548 | 4302242 | 6695 | - |
| Chr06 | 4304947 | 4318088 | 13142 | + |
| Chr06 | 4335049 | 4344283 | 9235 | + |
| Chr06 | 4465626 | 4472596 | 6971 | - |
| Chr06 | 4500001 | 4508968 | 8968 | + |
| Chr06 | 4858691 | 4863900 | 5210 | + |
| Chr06 | 5125799 | 5132631 | 6833 | + |
| Chr06 | 5268402 | 5278511 | 10110 | - |
| Chr06 | 5284174 | 5295677 | 11504 | + |
| Chr06 | 5284174 | 5306998 | 22825 | + |
| Chr06 | 5446639 | 5453680 | 7042 | + |
| Chr06 | 5457842 | 5462279 | 4438 | + |
| Chr06 | 5540568 | 5546703 | 6136 | - |
| Chr06 | 5599298 | 5606607 | 7310 | + |

| | | | | |
|---|---|---|---|---|
| Chr06 | 5663160 | 5668168 | 5009 | + |
| Chr06 | 5803479 | 5808913 | 5435 | + |
| Chr06 | 5811898 | 5818859 | 6962 | - |
| Chr06 | 5865869 | 5878538 | 12670 | + |
| Chr06 | 6047593 | 6052953 | 5361 | - |
| Chr06 | 6387740 | 6399525 | 11786 | - |
| Chr06 | 6414113 | 6421104 | 6992 | + |
| Chr06 | 6478874 | 6491435 | 12562 | + |
| Chr06 | 6556276 | 6562748 | 6473 | + |
| Chr06 | 6595792 | 6600752 | 4961 | - |
| Chr06 | 6613736 | 6619029 | 5294 | - |
| Chr06 | 6719729 | 6725104 | 5376 | + |
| Chr06 | 6742654 | 6750216 | 7563 | + |
| Chr06 | 6851534 | 6862114 | 10581 | + |
| Chr06 | 7028269 | 7039010 | 10742 | - |
| Chr06 | 7266016 | 7277291 | 11276 | - |
| Chr06 | 7291321 | 7298476 | 7156 | - |
| Chr06 | 7371457 | 7382663 | 11207 | - |
| Chr06 | 7474641 | 7481121 | 6481 | - |
| Chr06 | 7609782 | 7616593 | 6812 | + |
| Chr06 | 7788672 | 7799236 | 10565 | + |
| Chr06 | 7823501 | 7833004 | 9504 | - |
| Chr06 | 8142115 | 8151622 | 9508 | + |
| Chr06 | 8203081 | 8223433 | 20353 | + |
| Chr06 | 8432078 | 8438991 | 6914 | - |
| Chr06 | 8434145 | 8438991 | 4847 | - |
| Chr06 | 8572281 | 8577192 | 4912 | - |
| Chr06 | 8705002 | 8710606 | 5605 | + |
| Chr06 | 8756988 | 8762202 | 5215 | - |
| Chr06 | 8952292 | 8957016 | 4725 | - |
| Chr06 | 9004374 | 9011091 | 6718 | - |
| Chr06 | 9221335 | 9231555 | 10221 | - |

| | | | | |
|---|---|---|---|---|
| Chr06 | 9264274 | 9269700 | 5427 | - |
| Chr06 | 9538736 | 9551276 | 12541 | - |
| Chr06 | 9629879 | 9638659 | 8781 | + |
| Chr06 | 9772338 | 9777529 | 5192 | + |
| Chr06 | 9974983 | 9983891 | 8909 | - |
| Chr06 | 10357950 | 10368462 | 10513 | + |
| Chr06 | 10357950 | 10381308 | 23359 | + |
| Chr06 | 10389207 | 10394583 | 5377 | - |
| Chr06 | 10399954 | 10405126 | 5173 | + |
| Chr06 | 10720260 | 10727079 | 6820 | + |
| Chr06 | 11106272 | 11111933 | 5662 | + |
| Chr06 | 11172504 | 11174428 | 1925 | - |
| Chr06 | 11280748 | 11286272 | 5525 | - |
| Chr06 | 11369440 | 11374129 | 4690 | + |
| Chr06 | 11418952 | 11424337 | 5386 | - |
| Chr06 | 11464920 | 11477415 | 12496 | + |
| Chr06 | 11568685 | 11575832 | 7148 | + |
| Chr06 | 11733918 | 11739337 | 5420 | - |
| Chr06 | 12051156 | 12056361 | 5206 | - |
| Chr06 | 12059721 | 12064715 | 4995 | + |
| Chr06 | 12070462 | 12082640 | 12179 | + |
| Chr06 | 12071022 | 12076299 | 5278 | - |
| Chr06 | 12190976 | 12196142 | 5167 | + |
| Chr06 | 12571792 | 12587639 | 15848 | + |
| Chr06 | 12709989 | 12721434 | 11446 | - |
| Chr06 | 12864864 | 12881297 | 16434 | - |
| Chr06 | 12944163 | 12949502 | 5340 | - |
| Chr06 | 13312016 | 13317434 | 5419 | - |
| Chr06 | 13426695 | 13433393 | 6699 | + |
| Chr06 | 13702311 | 13712843 | 10533 | + |
| Chr06 | 13798497 | 13803676 | 5180 | + |
| Chr06 | 13861340 | 13866016 | 4677 | - |

| | | | | |
|---|---|---|---|---|
| Chr06 | 14338537 | 14351215 | 12679 | + |
| Chr06 | 14463611 | 14468509 | 4899 | + |
| Chr06 | 14760934 | 14773167 | 12234 | - |
| Chr06 | 14761892 | 14768866 | 6975 | + |
| Chr06 | 14903989 | 14914174 | 10186 | - |
| Chr06 | 15547656 | 15554723 | 7068 | + |
| Chr06 | 15556609 | 15563246 | 6638 | + |
| Chr06 | 15629900 | 15641136 | 11237 | + |
| Chr06 | 16453802 | 16459360 | 5559 | - |
| Chr06 | 16809731 | 16815027 | 5297 | + |
| Chr06 | 16819930 | 16825293 | 5364 | - |
| Chr06 | 16867084 | 16877403 | 10320 | + |
| Chr06 | 17673144 | 17678675 | 5532 | + |
| Chr06 | 18064195 | 18069728 | 5534 | + |
| Chr06 | 18447907 | 18453340 | 5434 | + |
| Chr06 | 18718282 | 18723463 | 5182 | - |
| Chr06 | 19351092 | 19357788 | 6697 | - |
| Chr06 | 19572076 | 19577495 | 5420 | - |
| Chr06 | 20358128 | 20365828 | 7701 | - |
| Chr06 | 20720377 | 20725444 | 5068 | + |
| Chr06 | 21165697 | 21171296 | 5600 | - |
| Chr06 | 21323628 | 21329040 | 5413 | - |
| Chr06 | 21405240 | 21410749 | 5510 | - |
| Chr06 | 21735494 | 21745018 | 9525 | - |
| Chr06 | 21859311 | 21866315 | 7005 | + |
| Chr06 | 21923907 | 21929925 | 6019 | + |
| Chr06 | 22275638 | 22283110 | 7473 | + |
| Chr06 | 22378174 | 22387524 | 9351 | + |
| Chr06 | 22707435 | 22712558 | 5124 | - |
| Chr06 | 23525793 | 23531258 | 5466 | - |
| Chr06 | 23697356 | 23707972 | 10617 | - |
| Chr06 | 23805916 | 23811932 | 6017 | + |

| | | | | |
|---|---|---|---|---|
| Chr06 | 24037466 | 24047352 | 9887 | - |
| Chr06 | 24063971 | 24069278 | 5308 | + |
| Chr06 | 24073638 | 24083199 | 9562 | + |
| Chr06 | 24095122 | 24107184 | 12063 | + |
| Chr06 | 24177213 | 24182477 | 5265 | + |
| Chr07 | 25846 | 32147 | 6302 | + |
| Chr07 | 407075 | 411535 | 4461 | + |
| Chr07 | 752845 | 757904 | 5060 | + |
| Chr07 | 1376936 | 1382950 | 6015 | + |
| Chr07 | 1411163 | 1418486 | 7324 | - |
| Chr07 | 1790137 | 1795541 | 5405 | + |
| Chr07 | 1816601 | 1821970 | 5370 | - |
| Chr07 | 2224576 | 2236501 | 11926 | - |
| Chr07 | 2661252 | 2667134 | 5883 | + |
| Chr07 | 2674622 | 2679135 | 4514 | - |
| Chr07 | 2705536 | 2715678 | 10143 | - |
| Chr07 | 2767715 | 2773127 | 5413 | - |
| Chr07 | 2774654 | 2780249 | 5596 | - |
| Chr07 | 2893808 | 2899077 | 5270 | - |
| Chr07 | 3091736 | 3101403 | 9668 | + |
| Chr07 | 3203714 | 3212480 | 8767 | + |
| Chr07 | 3440388 | 3450011 | 9624 | + |
| Chr07 | 3469128 | 3475534 | 6407 | + |
| Chr07 | 3666761 | 3671794 | 5034 | + |
| Chr07 | 4020968 | 4026158 | 5191 | - |
| Chr07 | 4098428 | 4105260 | 6833 | - |
| Chr07 | 4402932 | 4409396 | 6465 | + |
| Chr07 | 5401421 | 5407342 | 5922 | + |
| Chr07 | 5666792 | 5671835 | 5044 | - |
| Chr07 | 5997756 | 6012901 | 15146 | - |
| Chr07 | 6167960 | 6171794 | 3835 | - |
| Chr07 | 6506800 | 6510653 | 3854 | + |

| | | | | |
|---|---|---|---|---|
| Chr07 | 7349956 | 7358850 | 8895 | + |
| Chr07 | 7915340 | 7917725 | 2386 | + |
| Chr07 | 8124547 | 8129555 | 5009 | + |
| Chr07 | 9296523 | 9303611 | 7089 | + |
| Chr07 | 9449269 | 9462896 | 13628 | + |
| Chr07 | 9659560 | 9666654 | 7095 | - |
| Chr07 | 9735293 | 9740489 | 5197 | + |
| Chr07 | 9763862 | 9773644 | 9783 | + |
| Chr07 | 9806384 | 9815999 | 9616 | - |
| Chr07 | 9842519 | 9854527 | 12009 | + |
| Chr07 | 9953999 | 9960491 | 6493 | + |
| Chr07 | 10016575 | 10023856 | 7282 | - |
| Chr07 | 10108056 | 10120846 | 12791 | - |
| Chr07 | 10141806 | 10144075 | 2270 | + |
| Chr07 | 10248693 | 10258349 | 9657 | + |
| Chr07 | 10289452 | 10295689 | 6238 | + |
| Chr07 | 10378718 | 10389408 | 10691 | - |
| Chr07 | 10597216 | 10607595 | 10380 | + |
| Chr07 | 10631177 | 10637669 | 6493 | + |
| Chr07 | 10643282 | 10652201 | 8920 | + |
| Chr07 | 10688428 | 10694084 | 5657 | - |
| Chr07 | 10702177 | 10714364 | 12188 | + |
| Chr07 | 10789863 | 10799811 | 9949 | + |
| Chr07 | 10900129 | 10904282 | 4154 | - |
| Chr07 | 11293134 | 11306051 | 12918 | + |
| Chr07 | 11423842 | 11427995 | 4154 | + |
| Chr07 | 11813974 | 11818739 | 4766 | + |
| Chr07 | 11883049 | 11888352 | 5304 | + |
| Chr07 | 11999594 | 12014363 | 14770 | - |
| Chr07 | 12042004 | 12047350 | 5347 | - |
| Chr07 | 12277756 | 12283055 | 5300 | - |
| Chr07 | 12299025 | 12312499 | 13475 | - |

| | | | | |
|---|---|---|---|---|
| Chr07 | 12299229 | 12311474 | 12246 | + |
| Chr07 | 12299229 | 12312499 | 13271 | - |
| Chr07 | 12301843 | 12305718 | 3876 | - |
| Chr07 | 12459459 | 12464130 | 4672 | + |
| Chr07 | 12648158 | 12656473 | 8316 | + |
| Chr07 | 12765466 | 12774463 | 8998 | + |
| Chr07 | 12792368 | 12797298 | 4931 | - |
| Chr07 | 13067700 | 13083183 | 15484 | - |
| Chr07 | 13068743 | 13081783 | 13041 | - |
| Chr07 | 13077275 | 13083204 | 5930 | + |
| Chr07 | 13098176 | 13107823 | 9648 | + |
| Chr07 | 13190823 | 13197351 | 6529 | - |
| Chr07 | 13614234 | 13620465 | 6232 | + |
| Chr07 | 13636876 | 13642082 | 5207 | - |
| Chr07 | 13762568 | 13770082 | 7515 | + |
| Chr07 | 13875079 | 13886817 | 11739 | + |
| Chr07 | 13892688 | 13905448 | 12761 | + |
| Chr07 | 13946916 | 13953030 | 6115 | + |
| Chr07 | 14003642 | 14009895 | 6254 | - |
| Chr07 | 14006271 | 14009895 | 3625 | - |
| Chr07 | 14064071 | 14074767 | 10697 | - |
| Chr07 | 14206537 | 14220113 | 13577 | - |
| Chr07 | 14288501 | 14295032 | 6532 | - |
| Chr07 | 14403245 | 14408115 | 4871 | + |
| Chr07 | 14412367 | 14417534 | 5168 | + |
| Chr07 | 14672813 | 14676621 | 3809 | - |
| Chr07 | 14699727 | 14705196 | 5470 | - |
| Chr07 | 14915715 | 14920633 | 4919 | + |
| Chr07 | 14921096 | 14926397 | 5302 | + |
| Chr07 | 16031133 | 16036177 | 5045 | + |
| Chr07 | 16557447 | 16559904 | 2458 | - |
| Chr07 | 17195440 | 17200200 | 4761 | + |

| | | | | |
|---|---|---|---|---|
| Chr07 | 17456346 | 17462616 | 6271 | + |
| Chr07 | 18233014 | 18241941 | 8928 | - |
| Chr07 | 18685664 | 18691781 | 6118 | + |
| Chr07 | 18732803 | 18737787 | 4985 | - |
| Chr07 | 18889423 | 18895008 | 5586 | + |
| Chr07 | 19793767 | 19799153 | 5387 | + |
| Chr07 | 20031982 | 20037129 | 5148 | - |
| Chr07 | 20163134 | 20173295 | 10162 | + |
| Chr07 | 20381625 | 20387816 | 6192 | + |
| Chr07 | 21724234 | 21730648 | 6415 | - |
| Chr07 | 22032817 | 22039079 | 6263 | + |
| Chr07 | 22192358 | 22203284 | 10927 | - |
| Chr07 | 22547004 | 22552623 | 5620 | + |
| Chr07 | 22575635 | 22581256 | 5622 | - |
| Chr07 | 22603317 | 22608959 | 5643 | + |
| Chr07 | 22728891 | 22734239 | 5349 | - |
| Chr08 | 49109 | 60049 | 10941 | - |
| Chr08 | 88647 | 101426 | 12780 | - |
| Chr08 | 255217 | 260850 | 5634 | + |
| Chr08 | 349992 | 357224 | 7233 | + |
| Chr08 | 380099 | 387101 | 7003 | - |
| Chr08 | 578204 | 589509 | 11306 | + |
| Chr08 | 578346 | 584772 | 6427 | + |
| Chr08 | 584774 | 589604 | 4831 | - |
| Chr08 | 945740 | 948122 | 2383 | + |
| Chr08 | 1158715 | 1164203 | 5489 | - |
| Chr08 | 1184729 | 1195151 | 10423 | + |
| Chr08 | 1184729 | 1195753 | 11025 | + |
| Chr08 | 1184760 | 1195151 | 10392 | + |
| Chr08 | 1184760 | 1195299 | 10540 | + |
| Chr08 | 1184760 | 1195343 | 10584 | + |
| Chr08 | 1457908 | 1463087 | 5180 | - |

| | | | | |
|---|---|---|---|---|
| Chr08 | 1518692 | 1525741 | 7050 | + |
| Chr08 | 1585137 | 1590683 | 5547 | + |
| Chr08 | 1597521 | 1602659 | 5139 | + |
| Chr08 | 1597596 | 1602659 | 5064 | + |
| Chr08 | 1642129 | 1646798 | 4670 | + |
| Chr08 | 1804612 | 1810236 | 5625 | - |
| Chr08 | 1936741 | 1941721 | 4981 | - |
| Chr08 | 2013675 | 2018800 | 5126 | + |
| Chr08 | 2086677 | 2089076 | 2400 | + |
| Chr08 | 2145432 | 2150699 | 5268 | + |
| Chr08 | 2395150 | 2399760 | 4611 | + |
| Chr08 | 2466453 | 2480124 | 13672 | + |
| Chr08 | 2496476 | 2502073 | 5598 | + |
| Chr08 | 2518429 | 2523853 | 5425 | - |
| Chr08 | 2678745 | 2683960 | 5216 | + |
| Chr08 | 2685142 | 2689959 | 4818 | - |
| Chr08 | 2718203 | 2723197 | 4995 | - |
| Chr08 | 2812105 | 2815223 | 3119 | + |
| Chr08 | 2867725 | 2876449 | 8725 | - |
| Chr08 | 3045165 | 3053445 | 8281 | + |
| Chr08 | 3074780 | 3077788 | 3009 | - |
| Chr08 | 3181729 | 3186262 | 4534 | - |
| Chr08 | 3219851 | 3230291 | 10441 | - |
| Chr08 | 3463838 | 3474438 | 10601 | + |
| Chr08 | 3579710 | 3584665 | 4956 | - |
| Chr08 | 3600233 | 3610953 | 10721 | - |
| Chr08 | 3668911 | 3673041 | 4131 | - |
| Chr08 | 3809495 | 3814688 | 5194 | - |
| Chr08 | 3883930 | 3895207 | 11278 | + |
| Chr08 | 3955690 | 3962962 | 7273 | + |
| Chr08 | 4057475 | 4068505 | 11031 | - |
| Chr08 | 4091612 | 4097175 | 5564 | + |

| | | | | |
|---|---|---|---|---|
| Chr08 | 4103997 | 4109128 | 5132 | + |
| Chr08 | 4158625 | 4163368 | 4744 | + |
| Chr08 | 4165850 | 4171231 | 5382 | + |
| Chr08 | 4228679 | 4234431 | 5753 | + |
| Chr08 | 4238689 | 4240322 | 1634 | + |
| Chr08 | 4311369 | 4316403 | 5035 | - |
| Chr08 | 4385614 | 4387710 | 2097 | - |
| Chr08 | 5329125 | 5334704 | 5580 | - |
| Chr08 | 5828897 | 5834293 | 5397 | + |
| Chr08 | 6500827 | 6511296 | 10470 | + |
| Chr08 | 7066666 | 7068569 | 1904 | + |
| Chr08 | 7150579 | 7157944 | 7366 | + |
| Chr08 | 7222685 | 7228192 | 5508 | + |
| Chr08 | 7397031 | 7399402 | 2372 | - |
| Chr08 | 7407697 | 7414693 | 6997 | + |
| Chr08 | 7423283 | 7428889 | 5607 | - |
| Chr08 | 7969386 | 7991157 | 21772 | + |
| Chr08 | 8067736 | 8077533 | 9798 | + |
| Chr08 | 8147715 | 8153268 | 5554 | + |
| Chr08 | 8163171 | 8168756 | 5586 | + |
| Chr08 | 8221061 | 8227926 | 6866 | + |
| Chr08 | 8247332 | 8249338 | 2007 | + |
| Chr08 | 8306860 | 8313518 | 6659 | - |
| Chr08 | 8381192 | 8386779 | 5588 | + |
| Chr08 | 8454804 | 8460315 | 5512 | + |
| Chr08 | 8708768 | 8714211 | 5444 | + |
| Chr08 | 8752662 | 8778513 | 25852 | - |
| Chr08 | 8956402 | 8961825 | 5424 | - |
| Chr08 | 9203785 | 9215484 | 11700 | - |
| Chr08 | 9302151 | 9307362 | 5212 | + |
| Chr08 | 9322859 | 9329981 | 7123 | - |
| Chr08 | 9353862 | 9363786 | 9925 | + |

| | | | | |
|---|---|---|---|---|
| Chr08 | 9532906 | 9534413 | 1508 | + |
| Chr08 | 9572640 | 9577175 | 4536 | + |
| Chr08 | 9651995 | 9663404 | 11410 | + |
| Chr08 | 9951731 | 9958008 | 6278 | + |
| Chr08 | 10101041 | 10103899 | 2859 | + |
| Chr08 | 10117275 | 10137828 | 20554 | + |
| Chr08 | 10222206 | 10227530 | 5325 | + |
| Chr08 | 10278167 | 10283373 | 5207 | + |
| Chr08 | 10392656 | 10401456 | 8801 | - |
| Chr08 | 10742594 | 10747908 | 5315 | + |
| Chr08 | 11204076 | 11206847 | 2772 | + |
| Chr08 | 11401902 | 11407472 | 5571 | - |
| Chr08 | 11473052 | 11478446 | 5395 | + |
| Chr08 | 11478808 | 11489726 | 10919 | + |
| Chr08 | 11545909 | 11553285 | 7377 | + |
| Chr08 | 11816816 | 11826309 | 9494 | - |
| Chr08 | 11816816 | 11826309 | 9494 | - |
| Chr08 | 11816816 | 11826309 | 9494 | - |
| Chr08 | 12031566 | 12037131 | 5566 | + |
| Chr08 | 12211486 | 12219610 | 8125 | - |
| Chr08 | 12369017 | 12373921 | 4905 | + |
| Chr08 | 12533368 | 12547630 | 14263 | + |
| Chr08 | 12534772 | 12561933 | 27162 | + |
| Chr08 | 12563988 | 12585158 | 21171 | + |
| Chr08 | 12565986 | 12570962 | 4977 | + |
| Chr08 | 12565986 | 12573274 | 7289 | + |
| Chr08 | 12565986 | 12573635 | 7650 | + |
| Chr08 | 12565986 | 12574899 | 8914 | + |
| Chr08 | 12565986 | 12577790 | 11805 | + |
| Chr08 | 12565986 | 12578689 | 12704 | + |
| Chr08 | 12565986 | 12581587 | 15602 | + |
| Chr08 | 12565986 | 12581944 | 15959 | + |

| | | | | |
|---|---|---|---|---|
| Chr08 | 12565986 | 12582659 | 16674 | + |
| Chr08 | 12565986 | 12582838 | 16853 | + |
| Chr08 | 12565986 | 12583018 | 17033 | + |
| Chr08 | 12565986 | 12584798 | 18813 | + |
| Chr08 | 12565986 | 12585391 | 19406 | + |
| Chr08 | 12565986 | 12589000 | 23015 | + |
| Chr08 | 12565986 | 12590439 | 24454 | + |
| Chr08 | 12565986 | 12591882 | 25897 | + |
| Chr08 | 12565986 | 12592059 | 26074 | + |
| Chr08 | 12565986 | 12592776 | 26791 | + |
| Chr08 | 12565986 | 12594399 | 28414 | + |
| Chr08 | 12566348 | 12584798 | 18451 | + |
| Chr08 | 12580892 | 12612142 | 31251 | + |
| Chr08 | 12581263 | 12611783 | 30521 | + |
| Chr08 | 12582169 | 12610888 | 28720 | + |
| Chr08 | 12582348 | 12610710 | 28363 | + |
| Chr08 | 12582529 | 12610529 | 28001 | + |
| Chr08 | 12600934 | 12611521 | 10588 | + |
| Chr08 | 12607664 | 12626271 | 18608 | + |
| Chr08 | 12607664 | 12633064 | 25401 | + |
| Chr08 | 12607664 | 12633064 | 25401 | + |
| Chr08 | 12607664 | 12633064 | 25401 | + |
| Chr08 | 12607664 | 12633064 | 25401 | + |
| Chr08 | 12607664 | 12633064 | 25401 | + |
| Chr08 | 12607664 | 12633064 | 25401 | + |
| Chr08 | 12735045 | 12749042 | 13998 | - |
| Chr08 | 13038866 | 13045586 | 6721 | - |
| Chr08 | 13738730 | 13744300 | 5571 | + |
| Chr08 | 13861517 | 13866421 | 4905 | + |
| Chr08 | 14014731 | 14019721 | 4991 | - |
| Chr08 | 14056009 | 14061464 | 5456 | - |
| Chr08 | 14297899 | 14300910 | 3012 | - |

| | | | | |
|---|---|---|---|---|
| Chr08 | 14505278 | 14510917 | 5640 | + |
| Chr08 | 15700848 | 15703465 | 2618 | - |
| Chr08 | 15738607 | 15740654 | 2048 | + |
| Chr08 | 17060402 | 17066010 | 5609 | + |
| Chr08 | 17092534 | 17101463 | 8930 | + |
| Chr08 | 17280121 | 17283599 | 3479 | - |
| Chr08 | 17322825 | 17328347 | 5523 | + |
| Chr08 | 17350280 | 17353720 | 3441 | + |
| Chr08 | 17416080 | 17420965 | 4886 | + |
| Chr08 | 17475940 | 17495037 | 19098 | + |
| Chr08 | 17495053 | 17511607 | 16555 | + |
| Chr08 | 17612203 | 17617779 | 5577 | - |
| Chr08 | 17636148 | 17641570 | 5423 | + |
| Chr08 | 17657770 | 17663001 | 5232 | + |
| Chr08 | 17946042 | 17957042 | 11001 | + |
| Chr08 | 17947315 | 17952794 | 5480 | - |
| Chr08 | 18219648 | 18224800 | 5153 | + |
| Chr08 | 18464973 | 18469921 | 4949 | + |
| Chr08 | 18684439 | 18689475 | 5037 | + |
| Chr08 | 19062170 | 19067428 | 5259 | + |
| Chr08 | 19334130 | 19339537 | 5408 | + |
| Chr08 | 19348772 | 19353616 | 4845 | - |
| Chr08 | 19543561 | 19558290 | 14730 | + |
| Chr08 | 19701299 | 19714965 | 13667 | - |
| Chr08 | 19729535 | 19738123 | 8589 | + |
| Chr08 | 19729535 | 19739174 | 9640 | - |
| Chr08 | 19846625 | 19851858 | 5234 | + |
| Chr08 | 20002184 | 20009541 | 7358 | + |
| Chr08 | 20062047 | 20067520 | 5474 | - |
| Chr09 | 258519 | 263729 | 5211 | - |
| Chr09 | 1328310 | 1333794 | 5485 | + |
| Chr09 | 1406145 | 1408520 | 2376 | - |

| | | | | |
|---|---|---|---|---|
| Chr09 | 1574516 | 1579708 | 5193 | - |
| Chr09 | 1626405 | 1629622 | 3218 | + |
| Chr09 | 1626405 | 1630949 | 4545 | + |
| Chr09 | 1880355 | 1896388 | 16034 | + |
| Chr09 | 2724162 | 2731869 | 7708 | - |
| Chr09 | 2844267 | 2849461 | 5195 | + |
| Chr09 | 2972795 | 2978321 | 5527 | - |
| Chr09 | 3021526 | 3027414 | 5889 | + |
| Chr09 | 3754559 | 3759867 | 5309 | - |
| Chr09 | 3958676 | 3969648 | 10973 | - |
| Chr09 | 4078608 | 4083976 | 5369 | + |
| Chr09 | 4514184 | 4519628 | 5445 | - |
| Chr09 | 4559727 | 4564726 | 5000 | + |
| Chr09 | 4703593 | 4709089 | 5497 | - |
| Chr09 | 4724771 | 4728599 | 3829 | + |
| Chr09 | 5236276 | 5247024 | 10749 | - |
| Chr09 | 5315206 | 5321317 | 6112 | - |
| Chr09 | 5363589 | 5369302 | 5714 | + |
| Chr09 | 5405872 | 5411288 | 5417 | - |
| Chr09 | 5749394 | 5754653 | 5260 | - |
| Chr09 | 5969494 | 5975380 | 5887 | + |
| Chr09 | 5996764 | 6020372 | 23609 | - |
| Chr09 | 6602119 | 6607093 | 4975 | - |
| Chr09 | 6706107 | 6711016 | 4910 | + |
| Chr09 | 6813406 | 6818444 | 5039 | + |
| Chr09 | 6844694 | 6850036 | 5343 | + |
| Chr09 | 7042565 | 7053047 | 10483 | + |
| Chr09 | 7777151 | 7783660 | 6510 | - |
| Chr09 | 7833311 | 7838794 | 5484 | - |
| Chr09 | 7990278 | 7995537 | 5260 | + |
| Chr09 | 8071288 | 8076630 | 5343 | + |
| Chr09 | 8177066 | 8178951 | 1886 | + |

| | | | | |
|---|---|---|---|---|
| Chr09 | 8348668 | 8359253 | 10586 | + |
| Chr09 | 8422057 | 8428893 | 6837 | - |
| Chr09 | 8430877 | 8435357 | 4481 | - |
| Chr09 | 8442234 | 8448510 | 6277 | - |
| Chr09 | 8442522 | 8448475 | 5954 | - |
| Chr09 | 8459911 | 8464981 | 5071 | + |
| Chr09 | 8527362 | 8537638 | 10277 | - |
| Chr09 | 8748309 | 8755122 | 6814 | - |
| Chr09 | 8801284 | 8806514 | 5231 | - |
| Chr09 | 8836422 | 8851825 | 15404 | - |
| Chr09 | 9112765 | 9118636 | 5872 | + |
| Chr09 | 9169450 | 9180521 | 11072 | + |
| Chr09 | 9169537 | 9180497 | 10961 | + |
| Chr09 | 9183661 | 9189008 | 5348 | - |
| Chr09 | 9251440 | 9260857 | 9418 | + |
| Chr09 | 9267168 | 9272502 | 5335 | + |
| Chr09 | 9337218 | 9340282 | 3065 | + |
| Chr09 | 9717448 | 9728306 | 10859 | + |
| Chr09 | 9752029 | 9754334 | 2306 | - |
| Chr09 | 9803354 | 9813179 | 9826 | - |
| Chr09 | 9881043 | 9891663 | 10621 | - |
| Chr09 | 10173047 | 10177941 | 4895 | + |
| Chr09 | 10262920 | 10273257 | 10338 | + |
| Chr09 | 10843880 | 10855667 | 11788 | + |
| Chr09 | 10949892 | 10955082 | 5191 | + |
| Chr09 | 11132227 | 11142430 | 10204 | + |
| Chr09 | 11533556 | 11538278 | 4723 | - |
| Chr09 | 11789656 | 11795235 | 5580 | + |
| Chr09 | 12235745 | 12255412 | 19668 | + |
| Chr09 | 12237726 | 12245295 | 7570 | + |
| Chr09 | 12240464 | 12245295 | 4832 | + |
| Chr09 | 12350791 | 12364389 | 13599 | - |

| | | | | |
|---|---|---|---|---|
| Chr09 | 12761062 | 12775215 | 14154 | - |
| Chr09 | 13039495 | 13046569 | 7075 | - |
| Chr09 | 13534351 | 13539351 | 5001 | - |
| Chr09 | 13602189 | 13618089 | 15901 | - |
| Chr09 | 13609694 | 13618089 | 8396 | - |
| Chr09 | 14001379 | 14010868 | 9490 | - |
| Chr09 | 14027716 | 14044011 | 16296 | + |
| Chr09 | 14031150 | 14038145 | 6996 | + |
| Chr09 | 14145208 | 14160183 | 14976 | + |
| Chr09 | 14273214 | 14278738 | 5525 | + |
| Chr09 | 14425125 | 14426941 | 1817 | + |
| Chr09 | 14524012 | 14529576 | 5565 | - |
| Chr09 | 14918359 | 14928303 | 9945 | - |
| Chr09 | 14942661 | 14953216 | 10556 | + |
| Chr09 | 15109537 | 15114660 | 5124 | - |
| Chr09 | 15133552 | 15138469 | 4918 | + |
| Chr09 | 15222444 | 15228017 | 5574 | + |
| Chr09 | 15689686 | 15705512 | 15827 | + |
| Chr09 | 15819563 | 15826957 | 7395 | - |
| Chr09 | 15844274 | 15854213 | 9940 | + |
| Chr09 | 16148099 | 16158052 | 9954 | + |
| Chr09 | 16181144 | 16183357 | 2214 | + |
| Chr09 | 16317288 | 16332324 | 15037 | + |
| Chr09 | 16361815 | 16363782 | 1968 | + |
| Chr09 | 16362378 | 16363782 | 1405 | + |
| Chr09 | 16591004 | 16597010 | 6007 | - |
| Chr09 | 16755245 | 16760378 | 5134 | - |
| Chr09 | 17065696 | 17070920 | 5225 | - |
| Chr09 | 17198626 | 17203746 | 5121 | - |
| Chr09 | 17893129 | 17899645 | 6517 | - |
| Chr09 | 17971730 | 17987221 | 15492 | + |
| Chr09 | 18059189 | 18064317 | 5129 | + |

| | | | | |
|---|---|---|---|---|
| Chr09 | 18077185 | 18082569 | 5385 | - |
| Chr09 | 18178356 | 18187221 | 8866 | - |
| Chr09 | 18305076 | 18310516 | 5441 | - |
| Chr09 | 18389678 | 18395460 | 5783 | + |
| Chr09 | 18460865 | 18466466 | 5602 | - |
| Chr09 | 18773102 | 18780757 | 7656 | - |
| Chr09 | 19309217 | 19318250 | 9034 | - |
| Chr09 | 19616909 | 19627050 | 10142 | + |
| Chr09 | 19640647 | 19645940 | 5294 | + |
| Chr09 | 19669649 | 19676112 | 6464 | - |
| Chr09 | 19894503 | 19907229 | 12727 | + |
| Chr09 | 19999626 | 20018203 | 18578 | + |
| Chr09 | 20051994 | 20057274 | 5281 | + |
| Chr09 | 20146018 | 20148651 | 2634 | + |
| Chr09 | 20285531 | 20290757 | 5227 | + |
| Chr09 | 20359553 | 20364922 | 5370 | + |
| Chr09 | 20482281 | 20492223 | 9943 | + |
| Chr09 | 20734550 | 20739512 | 4963 | + |
| Chr09 | 20799815 | 20810754 | 10940 | - |
| Chr09 | 20926946 | 20932294 | 5349 | - |
| Chr09 | 20999692 | 21005060 | 5369 | + |
| Chr09 | 21148995 | 21158627 | 9633 | - |
| Chr09 | 21210851 | 21217292 | 6442 | + |
| Chr09 | 21350059 | 21355571 | 5513 | + |
| Chr09 | 21588830 | 21597273 | 8444 | - |
| Chr09 | 21615133 | 21628124 | 12992 | + |
| Chr09 | 21908981 | 21918475 | 9495 | - |
| Chr09 | 22002909 | 22007720 | 4812 | + |
| Chr09 | 22672640 | 22678128 | 5489 | + |
| Chr09 | 22734830 | 22740244 | 5415 | - |
| Chr09 | 22893048 | 22900374 | 7327 | + |
| Chr09 | 22920566 | 22930408 | 9843 | + |

| | | | | |
|---|---|---|---|---|
| Chr09 | 22941961 | 22948567 | 6607 | - |
| Chr09 | 22967044 | 22976937 | 9894 | + |
| Chr09 | 23378709 | 23388047 | 9339 | + |
| Chr09 | 23482256 | 23489620 | 7365 | - |
| Chr09 | 23594997 | 23601342 | 6346 | + |
| Chr09 | 23920279 | 23930060 | 9782 | + |
| Chr09 | 23987552 | 23989165 | 1614 | + |
| Chr09 | 24184262 | 24195315 | 11054 | + |
| Chr09 | 24243481 | 24256703 | 13223 | - |
| Chr09 | 24347186 | 24369596 | 22411 | + |
| Chr09 | 24374665 | 24379823 | 5159 | + |
| Chr09 | 24456212 | 24467304 | 11093 | - |
| Chr09 | 24547323 | 24561087 | 13765 | + |
| Chr09 | 24555218 | 24560284 | 5067 | + |
| Chr09 | 24555351 | 24561087 | 5737 | + |
| Chr09 | 24622481 | 24624448 | 1968 | + |
| Chr09 | 24699062 | 24705852 | 6791 | - |
| Chr09 | 25311446 | 25316465 | 5020 | - |
| Chr09 | 25588790 | 25596012 | 7223 | + |
| Chr09 | 25702950 | 25708237 | 5288 | + |
| Chr09 | 26069167 | 26078779 | 9613 | - |
| Chr09 | 26243336 | 26249449 | 6114 | + |
| Chr09 | 26299153 | 26301917 | 2765 | - |
| Chr09 | 26299153 | 26322396 | 23244 | - |
| Chr09 | 26344841 | 26346822 | 1982 | + |
| Chr09 | 26386352 | 26391503 | 5152 | - |
| Chr09 | 26633019 | 26641007 | 7989 | + |
| Chr09 | 26751456 | 26772618 | 21163 | - |
| Chr09 | 26775466 | 26786956 | 11491 | + |
| Chr09 | 26935162 | 26942053 | 6892 | + |
| Chr09 | 27258403 | 27263744 | 5342 | + |
| Chr09 | 27341710 | 27347365 | 5656 | + |

| | | | | |
|---|---|---|---|---|
| Chr09 | 27692055 | 27712787 | 20733 | - |
| Chr09 | 27763176 | 27780601 | 17426 | + |
| Chr09 | 27821009 | 27824280 | 3272 | + |
| Chr09 | 27821009 | 27824280 | 3272 | + |
| Chr09 | 27821081 | 27824208 | 3128 | + |
| Chr09 | 27844879 | 27870875 | 25997 | - |
| Chr09 | 27863837 | 27870894 | 7058 | - |
| Chr09 | 27888021 | 27912292 | 24272 | + |
| Chr09 | 27905298 | 27912295 | 6998 | + |
| Chr09 | 27942510 | 27952357 | 9848 | - |
| Chr09 | 28050372 | 28066281 | 15910 | + |
| Chr09 | 28191443 | 28211123 | 19681 | + |
| Chr09 | 28215479 | 28229722 | 14244 | + |
| Chr09 | 28246707 | 28265888 | 19182 | + |
| Chr09 | 28404193 | 28410804 | 6612 | + |
| Chr09 | 28404193 | 28411095 | 6903 | + |
| Chr09 | 28404566 | 28410804 | 6239 | + |
| Chr09 | 28404566 | 28411095 | 6530 | + |
| Chr09 | 28485733 | 28499803 | 14071 | + |
| Chr09 | 28488160 | 28496292 | 8133 | + |
| Chr09 | 28488160 | 28497759 | 9600 | + |
| Chr09 | 28625045 | 28629599 | 4555 | + |
| Chr09 | 28694070 | 28698699 | 4630 | + |
| Chr09 | 28704357 | 28715024 | 10668 | - |
| Chr09 | 28742265 | 28751088 | 8824 | + |
| Chr09 | 28761785 | 28767222 | 5438 | - |
| Chr09 | 28770380 | 28775535 | 5156 | - |
| Chr09 | 28786999 | 28792639 | 5641 | + |
| Chr09 | 28922434 | 28927309 | 4876 | - |
| Chr09 | 28950510 | 28962867 | 12358 | + |
| Chr09 | 28994155 | 29004982 | 10828 | - |
| Chr09 | 29081707 | 29091015 | 9309 | - |

| | | | | |
|---|---|---|---|---|
| Chr09 | 29118063 | 29130994 | 12932 | - |
| Chr09 | 29229097 | 29241139 | 12043 | + |
| Chr09 | 29483439 | 29491883 | 8445 | + |
| Chr09 | 29565172 | 29570796 | 5625 | + |
| Chr09 | 29620040 | 29625667 | 5628 | + |
| Chr09 | 29644098 | 29649717 | 5620 | - |
| Chr09 | 29652814 | 29657484 | 4671 | + |
| Chr09 | 29703850 | 29708326 | 4477 | - |
| Chr09 | 29823402 | 29826526 | 3125 | + |
| Chr09 | 29823402 | 29832147 | 8746 | + |
| Chr09 | 29840365 | 29845456 | 5092 | + |
| Chr09 | 29852738 | 29857999 | 5262 | - |
| Chr09 | 29953837 | 29960297 | 6461 | + |
| Chr09 | 30324120 | 30329498 | 5379 | - |
| Chr09 | 30505792 | 30513571 | 7780 | + |
| Chr09 | 31319260 | 31324258 | 4999 | + |
| Chr09 | 31816282 | 31821932 | 5651 | + |
| Chr09 | 31954103 | 31960253 | 6151 | - |
| Chr09 | 32186929 | 32198081 | 11153 | - |
| Chr09 | 32373608 | 32379228 | 5621 | - |
| Chr09 | 32753510 | 32758765 | 5256 | + |
| Chr09 | 32761314 | 32763278 | 1965 | + |
| Chr09 | 32764836 | 32769752 | 4917 | + |
| Chr09 | 32790322 | 32794609 | 4288 | + |
| Chr09 | 33094967 | 33100160 | 5194 | + |
| Chr09 | 33128382 | 33135638 | 7257 | - |
| Chr09 | 33388975 | 33390990 | 2016 | - |
